# Marine viral particles reveal an expansive repertoire of phage-parasitizing mobile elements

**DOI:** 10.1101/2022.07.26.501625

**Authors:** John M. Eppley, Steven J. Biller, Elaine Luo, Andrew Burger, Edward F. DeLong

## Abstract

Phage satellites are mobile genetic elements that propagate by parasitizing bacteriophage replication. We report here the discovery of abundant and diverse phage satellites that were packaged as concatemeric repeats within naturally occurring bacteriophage particles in seawater. These same phage-parasitizing mobile elements were found integrated in the genomes of dominant co-occurring bacterioplankton species. Like known phage satellites, many of the marine phage satellites encoded genes for integration, DNA replication, phage interference, and capsid assembly. Many also contained distinctive gene suites indicative of unique virus hijacking, phage interference and mobilization mechanisms. Marine phage satellite sequences were widespread in local and global oceanic virioplankton populations, reflecting their ubiquity, abundance, and temporal persistence in marine planktonic communities worldwide. Their gene content and putative life cycles suggest they may impact host-cell phage immunity and defense, lateral gene transfer, and bacteriophage-induced cell mortality and host and virus productivity. These previously unrecognized marine phage satellites therefore have potential to impact the ecology and evolution of bacteria and their bacteriophages in the ocean, and similar phage parasites likely thrive in many other microbial habitats as well.

**Significance statement:** Phage satellites are mobile genetic elements that parasitize bacteriophage, thereby exerting profound biological and ecological impacts. To date however, phage satellites have been found primarily in Gram-positive cocci and a few Gram-negative bacteria, many of which are human pathogens. Direct inspection of “wild” marine virus particles however, revealed that phage satellites are widely distributed in the sea, and that their genetic diversity, gene repertoires, and host ranges are much greater than previously supposed. Our analyses provide insight into their parasitic life cycles, potential satellite-helper-phage interactions, and reproductive strategies of these newly recognized phage-parasitizing mobile elements. Their properties, diversity and environmental distributions suggest they exert pervasive influence on marine plankton ecology and bacterial and virus evolution in the sea.

## Introduction

Bacterial viruses (aka bacteriophages) can impact host-cell phenotype, population structure, pathogenicity, host immunity, lateral gene transfer, and other diverse aspects of microbial physiology, ecology and evolution (1). Viruses can themselves be parasitized by small mobile genetic elements (MGEs) called phage satellites, that propagate by exploiting co-occurring bacteriophage reproduction machinery. Comprising a few known types - most notably satellite phage (e.g., the *Escherichia coli* phage P4/P2 system; (2, 3)), phage-inducible chromosomal islands (PICIs like those found in *Staphylococcus aureus* and other Gram-positive and Gramnegative bacteria; (4–6)), and PICI-like elements (PLEs in *Vibrio cholerae*; (7–9)) - such MGEs are genetically diverse. They do, however, share a common life cycle strategy which depends on propagation via the hijacking of “helper phage”, redirecting phage capsid assembly towards producing phage-like particles containing mobile elements instead of phage DNA (10).

Phage satellites can influence host cell phage immunity (*10,14*), the frequency of host gene transduction (9, 11, 12–14), host cell pathogenicity (4, 14, 15), and host cell and helper phage genome evolution (13, 16). While evidence suggests that some habitat and host diversity may exist among known phage satellites (6, 12, 15–18), they have been demonstrated to date only in Gram positive bacteria (18) and a few Enterobacteriaceae and Pastuerellaceae pathogens (6, 8, 10). The extant diversity, cellular host range, helper phage-satellite interactions, and ecology of phage satellites in other diverse habitats, however, remains largely unexplored.

### Many “wild” marine phage particles contain phage satellite concatemeric DNAs, instead of phage genomes

Previous work had shown that single-molecule genome sequencing of naturally occurring virus particles efficiently recovers complete phage genomes in single DNA reads (19). To further refine these analyses, we used density gradient fractionation to separate tailed phages from other types of marine particles, such as membrane vesicles and tailless phages, as well as cellular DNA contamination (Fig. 1A; (20)). Following single-molecule nanopore sequencing of virion encapsidated high molecular weight DNA, we discovered that a considerable fraction (0.6 %) of virion particles collected from 25m depth in the North Pacific Subtropical Gyre (NPSG), contained DNA encoding concatemeric repeats (Fig 1A; *SI Appendix*, Fig. S1; Dataset S2). Virion encapsidated concatemeric DNA lengths exceeded 100 kbp (Fig. 1B) and overlapped with the prevailing genome sizes of co-occurring bacteriophages (Fig. 1B; Dataset S2). We also observed DNA concatemers in the lower density membrane vesicle-enriched fraction, albeit at lower frequencies and with much shorter lengths, mostly < 20 kbp (Fig. 1B).

**Fig. 1.**
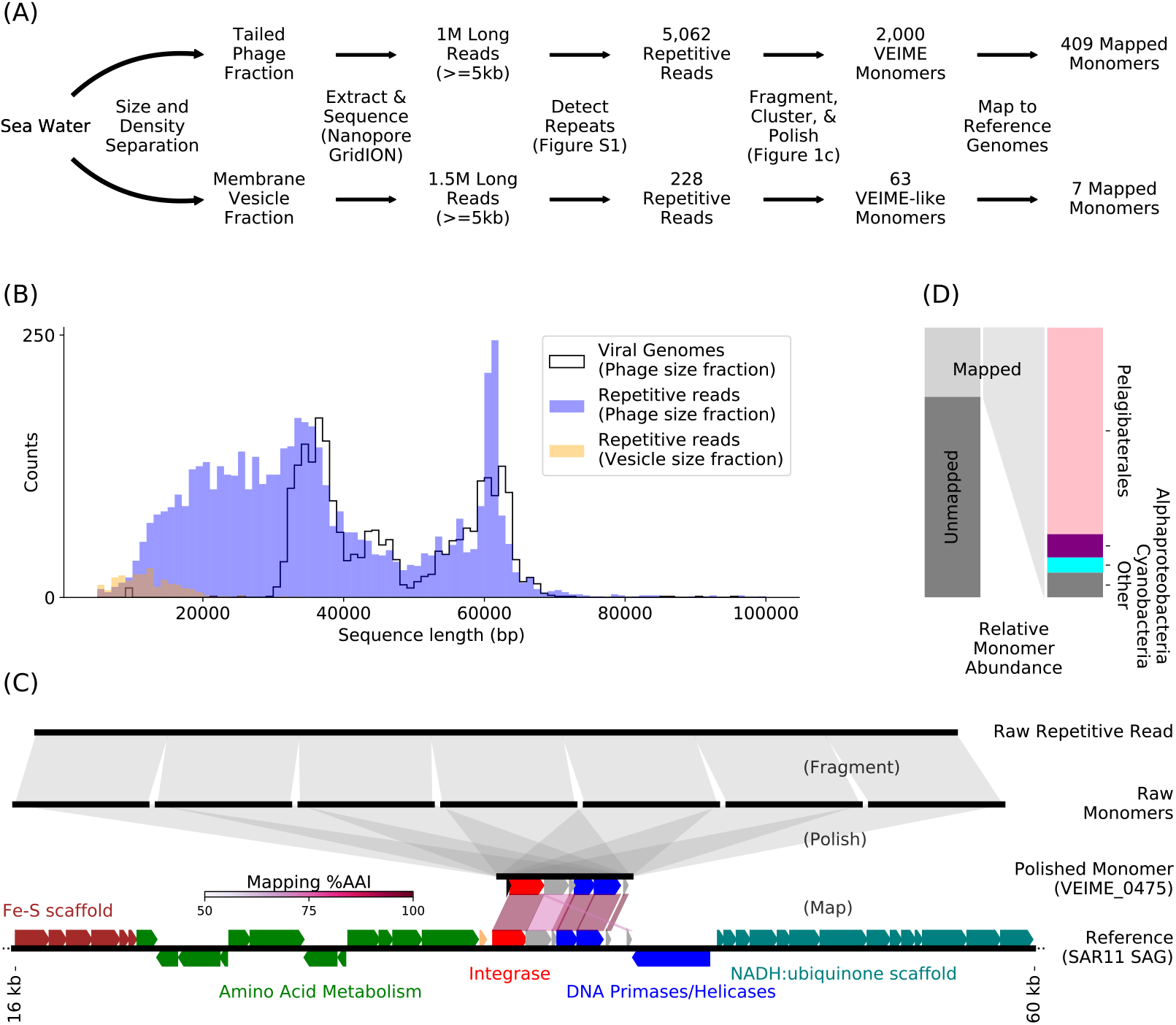
Concatemeric DNAs within phage particles facilitate discovery of new phage-parasitizing mobile genetic elements. The top panel **(A)** shows the workflow for preliminary identification of Virion Encapsidated Integrative Mobile Elements (VEIMEs), and Membrane Encapsulated Integrative Mobile Elements (MEIMEs) in fractionated samples. The histogram of concatemeric read lengths **(B)** shows peaks that correspond to common phage genome lengths, represented here by a histogram of the lengths of reads with direct terminal repeats (DTRs). The bottom panel **(C)** illustrates the fragmenting and self-polish steps of the workflow and shows a polished VEIME monomer aligned with a reference genome. The left side of panel **(D)** shows the relative abundance of VEIMEs and MEIMEs that mapped to genomes compared to those that did not. The right side of **(D)** displays the relative abundance of mapped VEIMEs, and their frequency of occurrence in different bacterial host taxa.

Polishing and de-replication of the repeated sequences in concatemers revealed a diverse set of novel phage-mobilized genetic elements referred to here as <u>V</u>irion <u>E</u>ncapsidated <u>I</u>ntegrative <u>M</u>obile <u>E</u>lements (VEIMEs). Within the single seawater sample analyzed here, we identified 2000 unique concatemer-derived monomeric VEIMEs in the tailed-phage-enriched fraction and 63 in the membrane-vesicle-enriched fraction (Fig. 1A; Dataset S2 and S3). About 20% of VEIMEs were also found integrated into the genomes of a variety of common bacterioplankton groups including *Pelagibacter* and other Alphaproteobacteria, *Prochlorococcus, Verrucomicrobia, Flavobacteria*, and additional bacterial taxa, demonstrating that these elements were widespread among diverse marine planktonic bacteria (Fig. 1, C and D; *SI Appendix*, Fig. S3; Dataset S3).

### Marine phage satellites have diverse, unique, and distinctive gene suites that differentiate them from previously described phage satellites

We next asked whether the marine VEIMEs might share some features with previously characterized human microbiome-associated phage satellites (i.e. P4-like satellite phage, PICIs, and PLEs). Using a bipartite graph of genomes and gene annotations of VEIMEs along with known phage satellites, we found that marine phage satellites partitioned into twelve different modules (Fig. 2). The majority of VEIMEs (98%) were in modules composed solely of VEIMEs, all of which showed significant intra-module gene content variation (e.g. Modules 1-4, 6, 8, 12; Fig. 2; *SI Appendix*, Fig. S2; Dataset S3, S5, and S6). The remaining modules included both human microbiome-associated reference phage satellites as well as VEIMEs (Fig. 2; Dataset S3, S4 and S5). Marine phage satellites in these modules, however, encoded very different gene suites than those of previously characterized phage satellites (*SI Appendix*, Fig. S2; Dataset S6). Different VEIME modules were also often associated with specific bacterial host taxa. For example, VEIMEs in three modules (Modules 2, 6, and 8) were found mostly in *Pelagibacter* and other Alphaproteobacteria host genomes, while Module 3 VEIMEs were associated with Cyanobacteria and Verrucomicrobia (*SI Appendix*, Fig. S2 and S3; Dataset S3 and S6).

**Fig 2.**
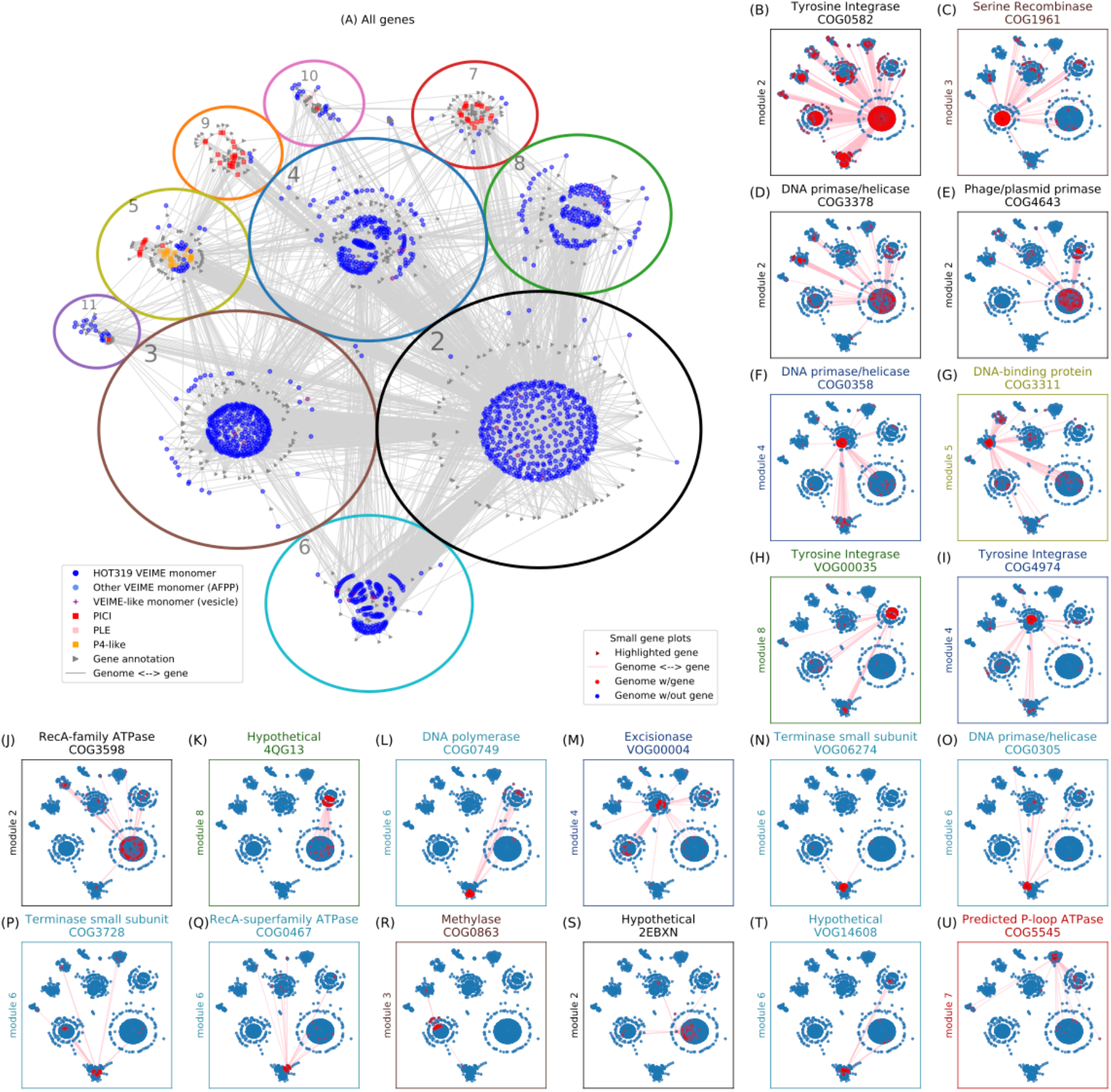
Bipartite graphing reveals interrelationships among VEIMEs and reference phage satellites. We constructed a bipartite graph by connecting EggNOG and VOG gene annotations (triangles) to the satellite genomes (other shapes) in which they were found. The displayed graph layout, used in all subplots, exaggerates module separation for clarity. **Large plot (A):** Module boundaries are proportional in area to the number of MGEs in each module. The shape and color of nodes indicate their type. All gene annotations are gray triangles. Published PICIs, P4-like phage, and PLEs are red, orange, and pink squares, respectively. VEIMEs are dark blue (from HOT319 75m) or light blue (AFPPs from ref. 21) circles. **Small plots:** Each small plot highlights one of the top 20 most common gene annotations, arranged in order of abundance (left to right, and top to bottom) starting from the COG0582 Integrase at the left of the top row, to the COG5545 ATPase at the right end of the bottom row. The specific gene referenced is shown as a dark red triangle. MGEs are shown as circles, and colored red if they contain the gene, and blue if not.

Analysis of VEIME gene repertoires indicated that most VEIMEs shared some common features reflecting their postulated integrative, replicative and parasitic life cycles (Fig. 2; Dataset S5 and S6). Most conspicuously, >80% of all VEIMEs encoded integrases, either a tyrosine integrase, a serine recombinase, or both (Figs. 2 and 3; figs. S2, S4, S5 and S6; Dataset S3, S5, and S6). Notably, the only reference phage satellites in our analyses that encoded a serine recombinase were the three reference PLEs associated with Module 10 (Fig. 2; (7, 10)). Other mobile element-like genes shared across many VEIME modules included some involved in phage satellite replication, capsid assembly, and DNA packaging, including DNA primase/helicases, the DNA-binding prophage transcriptional regulator AlpA, terminase small subunit homologues, motor-like ATPases, and excisionases (Fig. 2; figs. S2, S4, and S7; Dataset S5 and S6).

**Fig. 3.**
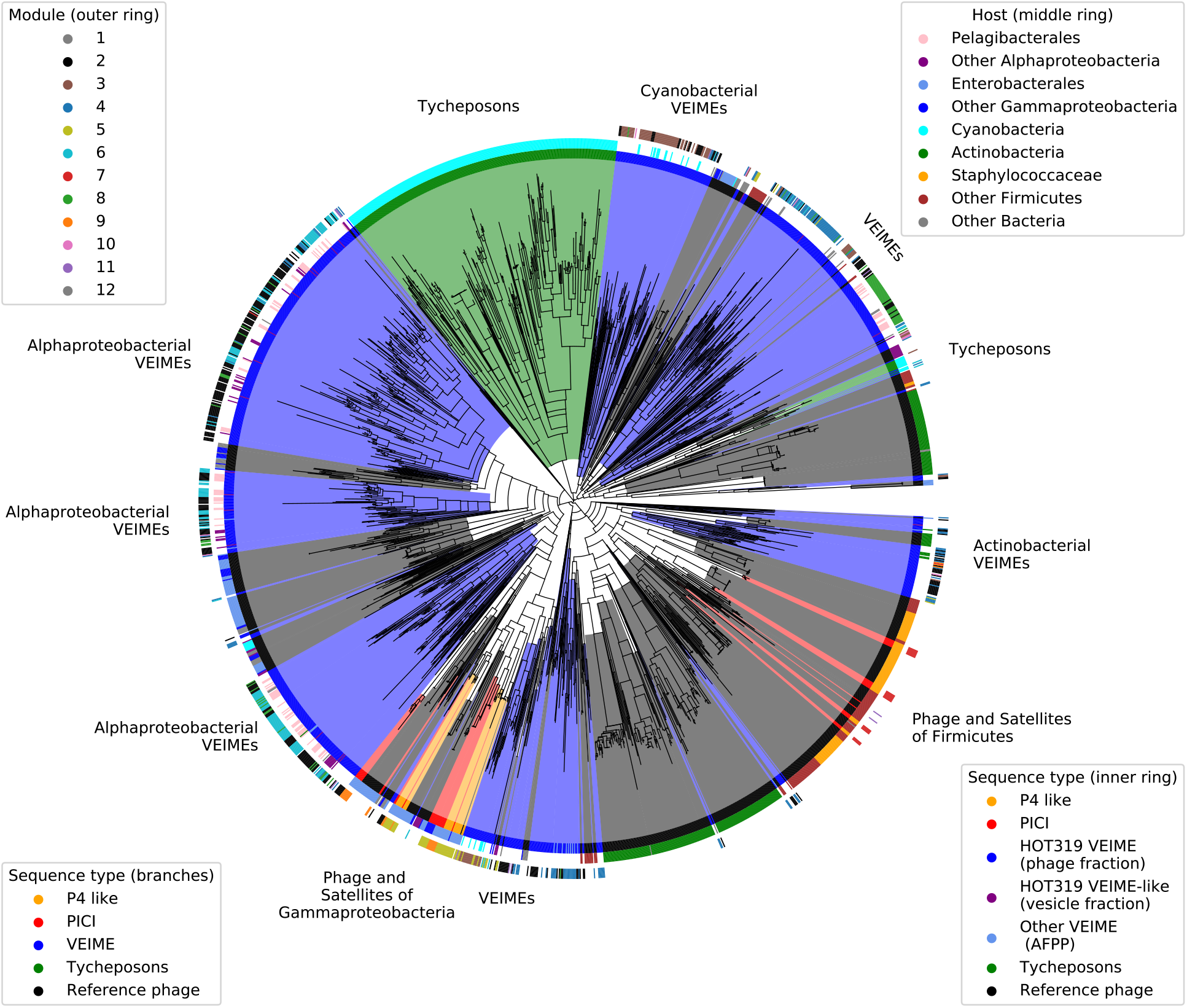
Relationships of tyrosine integrases among VEIMEs, reference phage satellites, and bacteriophages. A phylogenetic tree of tyrosine recombinases found in environmentally derived and reference phage satellites, bacteriophages, and tycheposons. Branches are colored by sequence type, with environmental satellites broken down further by sample in the first colored ring. The second ring indicates the taxonomy of known hosts with gaps for any unknown host associations. The outer ring indicates the module assignment from Figure 2.

Their extensive functional gene diversity and contents suggests that the marine phage satellites may use a range of mechanisms to reproduce and hijack helper phage replication cycles. All the marine VEIMEs were initially detected as concatemeric repeat sequences in virus particles, which is consistent with a plasmid-like rolling circle DNA replication mechanism. Although marine VEIME concatemeric DNAs were encapsidated in virion particles, the vast majority did not encode any capsid-like structural genes (Dataset S2 and S3), which many known phage satellites use to parasitize helper phage. Together with the correspondence of viral genome and VEIME concatemer size ranges (Fig. 1B), these data suggest that capsid remodeling (and concomitant capsid size reduction) is not as common a strategy in marine VEIMEs, as it is for previously described Gram-positive and Gram-negative phage satellite systems (6, 10, 18, 21).

While 91% of the Module 3 VEIMEs encoded a serine recombinase, tyrosine integrases predominated in the other modules (Figs. 2 and 3; figs. S2, S5, and S6; Dataset S5 and S6). The VEIME-only, tyrosine integrase-dominated modules could further be differentiated by their module-specific gene repertoires (Fig. 2; Dataset S5 and S6; figs. S2, S4, and S5). For example, although VEIMEs in the *Pelagibacter/*Alphaproteobacteria-associated groups (Modules 2, 6, and 8) shared some core replicative genes, each of these modules also contained unique genes that were rare or absent among the others (figs. S2, S4 and S5; Dataset S5 and S6). Module 2 VEIMEs encoded HNH endonucleases (22) and periplasmic serine proteases (23), whereas Module 6 VEIMEs contained small subunit terminases (*SI Appendix*, Fig. S7) and DNA polymerase genes (*SI Appendix*, Fig. S2, Dataset S6). In contrast, Module 8 VEIMEs encoded RecD-like helicases, hydrolases, and phage protein D14 genes (Fig. 2; Dataset S5 and S6).

Mechanistically, small subunit terminases have been found among previously described phage satellites that redirect helper phage replication towards the preferential packaging of phage satellite DNA (10), HNH endonucleases are also common components of phage DNA packaging machinery (22), and some serine proteases are known to be involved in procapsid maturation (23). Notably, the VEIME-encoded terminase small subunit sequences were most similar to homologs found within bona fide bacteriophage genomes recovered from the same habitat (*SI Appendix*, Fig. S7). Overall, the wide variety of functional gene suites observed among VEIMEs (Data SX and SY) suggests that considerable variability exists in the parasitic molecular interactions between marine phage parasites and their helper phages.

Although gene content analyses suggest that some similarities may exist between VEIME life cycle strategies and those of known phage satellites, some VEIMEs may also use different replication strategies. Unique genes in Module 3 included homologs of plasmid replication initiation and partitioning genes, hinting that some VEIMEs may employ a plasmid-like replication strategy. Module 4 VEIMEs contained the highest proportion of excisionases, methyltransferases, YwqK-like antitoxin and RecA-like recombinase genes (Fig. 2; figs. S2 and S4; Dataset S5 and S6) further expanding the list of potential mechanisms for phage interference employed by marine VEIMEs. The fact that some VEIMEs did not encode any genes previously shown to be involved in phage satellite replication, further implies that yet-to-be characterized helper phage interference and satellite DNA packaging remain to be discovered. Notably, integrase genes were not identified in ~20% of the VEIMEs, indicating that some VEIMEs might exist predominantly as plasmids, or take advantage of host or helper phage encoded integrases, or use as-yet unidentified replication and phage interference strategies. We also found VEIME-like concatemeric elements in the lower density membrane-vesicle-enriched fraction of our seawater sample (Fig 1A), suggesting that some of the mobile element concatemers might also be mobilized via extracellular vesicles (i.e. in the absence of any capsid), or potentially mobilize by hijacking tailless helper phages.

### Integrase gene phylogenies reveal that diverse marine phage satellites are different from previously characterized phage parasitizing mobile elements

VEIME gene phylogenies provided additional insight into the relationships amongst VEIMEs and other phage satellites and MGEs (Fig. 3; figs. S4, S5, S6, and S7). The tyrosine integrases found in the majority of VEIMEs formed several distinct clusters different from those of known phage satellites and bacteriophages (Fig. 3, figs. S4 and S5). The serine recombinases found in Module 3 also exhibited considerable diversity, with one VEIME integrase clade specifically associated with known PLE serine recombinases (*SI Appendix*, Fig. S6). Other VEIME-encoded serine recombinases appeared most closely associated with a new group of transposon-like MGEs (Tycheposons) recently reported in the cyanobacterium *Prochlorococcus* (24). Thus, VEIME gene content and integrase phylogenies both support the view that marine phage satellites are part of a large and diverse group, comprised of many subtypes most of which are distinct from previously characterized MGEs.

### Marine phage satellites are ubiquitous and abundant in the ocean

To assess the spatiotemporal distribution of VEIMEs, we surveyed their relative abundances in metagenomic data collected at our NPSG sampling site, and other global oceanic sampling stations (Fig. 4; *SI Appendix*, Fig. S8 (25)). The VEIMEs reported here, originally derived from a single 25m NPSG seawater sample, were present primarily in shallower water virioplankton size fractions, with some more common in surface waters, others more common at greater depths, and still others with more uniform depth distributions (Fig. 4A). Temporally, the VEIMEs appeared present year-round in the NPSG, with no clear seasonal pattern (Fig. 4B). VEIME homologs were also evident in virus enriched samples in other oceanic regions (25); Fig. 4C; *SI Appendix*, Fig. S8), primarily in shallow surface waters, or around the deep chlorophyll maximum layer (Fig. 4C). Additionally, these marine phage satellites were found within the genomes of bacteria originating from diverse locales around the globe (*SI Appendix*, Fig. S3), further reflecting their ubiquity and abundance in marine plankton. Taken together, these results suggest that VEIMEs are widespread and persistent features of global ocean planktonic microbiomes worldwide.

**Fig. 4.**
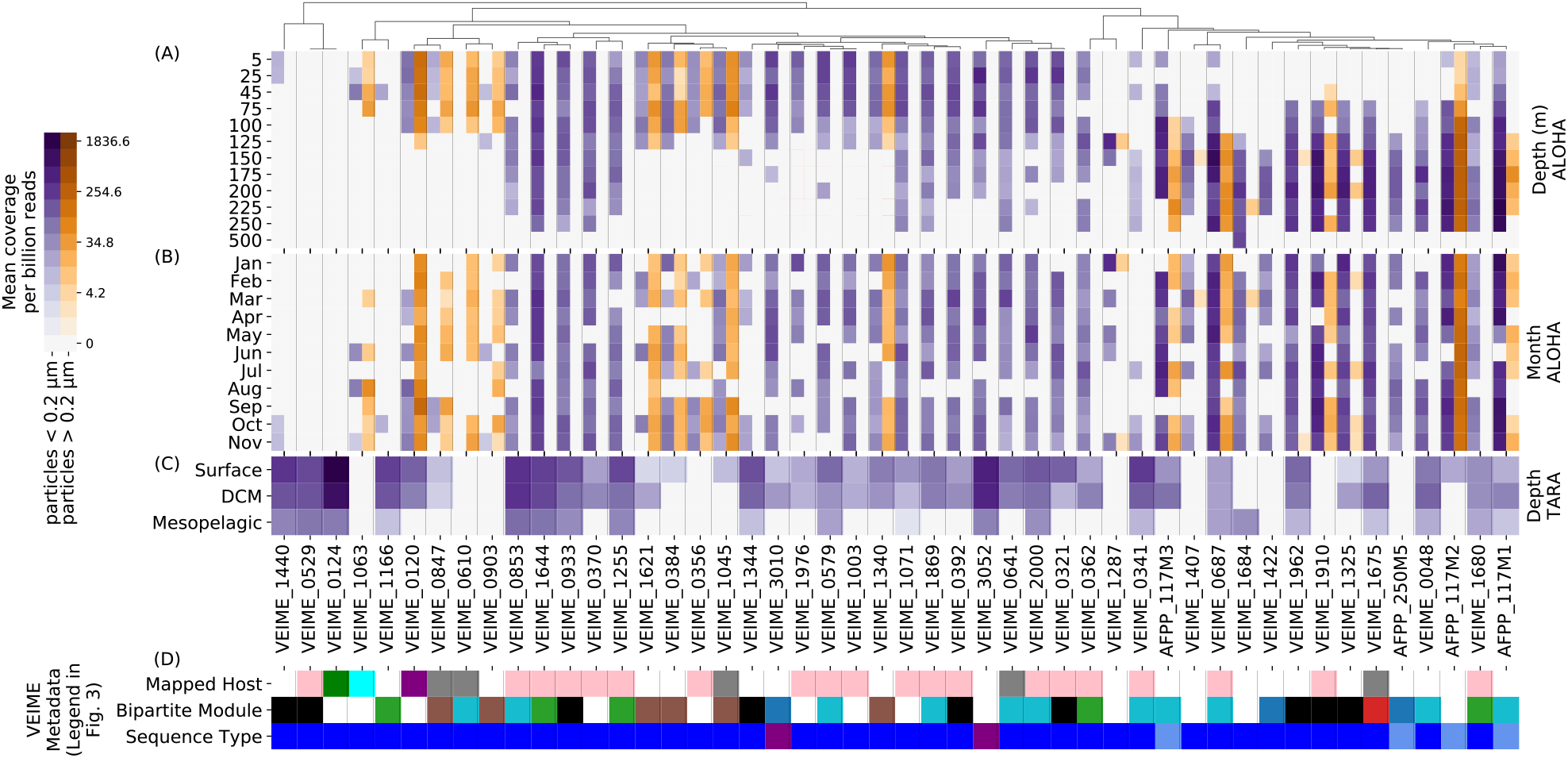
VEIME DNA abundances in marine picoplankton and virioplankton size fractions. We compared VEIMEs to assembled contigs from published short read metagenomes and used the normalized local metagenomic short read coverage of the contigs in the aligned regions as a proxy for environmental abundance of aligned VEIMEs. In **(A)** abundances in bacterioplankton (orange) and picoplankton (purple) metagenomes from Station ALOHA are averaged by depth and displayed side by side for each monomer. In **(B)**, the same data is aggregated by calendar month. In **(C),** abundance in TARA ocean viral metagenomes is shown. VEIME placement along the x-axis follows a hierarchical clustering (shown at the top) by correlation of abundances across all samples. In **D)**, metadata on hosts, modules and sequence types are shown, using the same annotation color scheme as in Figure 3.

The search for predominant cellular hosts of marine phage satellites provided new information on their biology and ecology. Unlike previously reported phage satellites found exclusively in copiotrophic bacteria, the marine phage satellites we report here occurred in some of the most oligotrophic bacterial species known. Specifically, the most prevalent cellular hosts of surface water marine VEIMEs were *Pelagibacter* and the cyanobacterium *Prochlorococcus*, among the most abundant bacterial groups present in open ocean surface waters (26, 27). Recently, the presence of temperate phage in oligotrophic *Pelagibacter* species has been validated (28, 29), consistent with our observations of phage satellites in the genomes of this ubiquitous bacterioplankton group. Similarly, cyanophages of *Prochlorococcus* often encode integrases, which allows them to integrate into host genomes (30). Among VEIME-containing bacterial genomes however (*SI Appendix*, Fig. S3), only a small percentage provided evidence of co-existing prophage in the same genome. Additionally, oligotrophic shallow waters in general contain a greater proportion of lytic to temperate phage, compared to deeper waters *(31*). It seems possible therefore that some marine VEIMEs, analogous to the life cycle of known PLEs (*12*), may parasitize invading lytic phage, in addition to temperate phages.

## Conclusion

Our new data and analyses expand considerably currently known phage satellite distributions, diversity, host ranges, and gene contents considerably. These new data also provide insight into marine satellite parasitic life cycles, helper phage interactions, and reproductive strategies. Phage satellites are known to have profound biological and ecological impacts, including provision of phage defense and immunity (5, 9, 11, 12) and diminishment of viral productivity that may also decrease virus-induced mortality. Phage satellites, along with other phage-like entities that package random genomic DNA in capsid-like structures like gene transfer agents (31, 32), can mobilize bacterial pathogenicity and other phenotypic traits (4, 8, 15) via lateral gene transfer (4, 8, 14, 16). Their abundance, ubiquity, and varieties in the ocean suggest their probable impacts on genome evolution and lateral gene transfer in host-cells and helper bacteriophage. While most phage satellites reported to date have been found in human-associated bacteria, many of which are pathogenic, our results suggest that phage-parasitizing MGEs are probably common components of other microbial ecosystems as well.

The abundance and association of marine phage satellites with diverse host cell taxa – including some of the most abundant microbes on the planet – highlights their potential for impacting virus and bacterial primary and secondary production in the sea. Their planktonic virus enablers are also abundant, with typical virion concentrations reaching ten times that of surrounding cells, around 10^4^ and 10^8^ ml^-1^ (33, 34). As such, viruses significantly impact microbial plankton mortality, and its downstream effects on primary and secondary productivity and carbon export to deep waters (33–35). Estimates have suggested that aquatic viruses can lyse as many as 20-40% of available prey cells per day (9,10), thereby contributing an estimated 145 gigatons to annual global carbon flux in tropical and subtropical seas (33). Both lytic and temperate bacteriophage are active in marine plankton, with lysogeny typically being thought to prevail under low nutrient concentrations and high virus-to-host cell ratios. More recent studies however have suggested that lysogeny may instead be more prevalent in habitats with greater bacterial production rates (36, 37). In either case, the discovery reported here that parasitic phage satellites are abundant and dynamic components of marine virioplankton, reinforces the notion that these MGEs may significantly impact viral productivity and virus-induced cell mortality in the ocean.

Accumulating evidence now suggests that mobile genetic elements frequently drive host-cell resistance to bacteriophage (38). In *Vibrio* species for example, a variety of phage defense MGE mechanisms appear to be in play. Along with phage-receptor mutations, these can include integrative and conjugative elements (ICEs), clustered regularly interspaced short palindromic repeat-Cas systems (CRISPR-Cas), abortive-infective systems (Abi), restriction modification systems (R-M), and other phage defense gene suites, that occupy large percentages of the flexible genome (39–41). Many antiphage systems are frequently co-located with mobile genetic elements reinforcing the idea that phage predation (and defenses against it) accelerate lateral gene transfer, and help to drive bacterial evolution (40–43). Known phage satellites, like PICIs and PLEs, have recently been recognized as key elements of antiphage defense systems (12, 13, 44, 45). Some of the marine phage satellites we report here encoded known phage defense and interference genes (including CRISPR-Cas, R-M, phage transcriptional regulators, and Abi-like genes), suggesting they too may confer diverse phage defense functionalities to their planktonic cellular hosts. As is the case with other mobile genetic elements (13, 39, 41, 43), these previously unrecognized marine phage satellites likely also impact the evolutionary rates and genetic diversity of host cells and the bacteriophage partners that they parasitize.

## Materials and methods

### Concentration of virus and vesicle enriched samples from seawater

Seawater was collected from a depth of 25 meters, and prefiltered to minimize cellular contamination. The resulting virus and membrane vesicle enriched filtrate was concentrated via tangential flow filtration (TFF) before DNA extraction, as outlined below. The 25 m seawater sample was collected on ton January 31^st^ and February 1^st^ (HOT-319 cruise), 2020 at Station ALOHA (22°45’ N, 158° W); http://hahana.soest.hawaii.edu/hot/).

A total of 440 L of seawater was collected using a Niskin bottle rosette attached to a conductivity-temperature-depth (CTD) package. The seawater was pre-filtered by peristatic pumping through a 0.1 uM Supor cartridge filter (Acropak 500, Pall, USA). The resulting virus-enriched filtrate was concentrated by tangential flow filtration (TFF) over a 30 kDa filter (Biomax 30 kDa membrane, catalogue #: P3B030D01, Millipore). Subsequently, the retentate was reduced to a volume of ~100 mL and stored at 4°C. Next, ~10L of < 30 Kd permeate reserved from the tangential flow filtration was added to the recirculation vessel and the system was run for an additional 30 minutes to release virus particles trapped in the filter. After 30 minutes the retentate volume was reduced to ~100 mL and this flushing retentate was added to the ~100 mL of the initial retentate, resulting in a final retentate volume of ~200 mL. The virus-containing retentates were stored at 4°C until further processing.

### Small particle fractionation and purification

Particles were isolated and purified from the TFF concentrate by ultracentrifugation at 32,000 rpm (~126,000 x*g*) for 2 hours at 4°C in a Beckman-Coulter SW32Ti rotor. The pelleted material was resuspended in residual seawater and separated across an iodixanol density gradient (Optiprep; Sigma-Aldrich). The gradient was formed as follows: successive 0.5 mL layers of iodixanol (45%, 40%, 35%, 30%, 25%, 20%, 15%, and 10%; all in a 3.5% (w/v) NaCl, 3.75 mM TAPS pH 8, 5 mM CaCl_2_ buffer background) were placed in a 4 mL UltraClear ultracentrifuge tube (Beckman-Coulter), with the particle sample as the top layer. The gradient was spun at 45,000 rpm (~200,000 x *g)* for 6 hours at 4 °C in a SW60Ti rotor (Beckman Coulter). Successive 0.4 mL fractions were collected by pipetting and their densities measured. Fractions between 1.14 – 1.19 g/mL were pooled to form the “vesicle enriched” sample (containing both extracellular vesicles and potentially some non-tailed viruses), and fractions >1.2 g/mL were combined to form the “tailed phage” sample (20, 46, 47). Particles in the tailed phage fraction were washed twice by diluting with filter-sterilized buffer (3.5% (w/v) NaCl, 3.75 mM TAPS pH 8, 5 mM CaCl_2_) followed by ultracentrifugal pelleting (32,000 rpm, 2 hrs, 4°C, SW60Ti rotor). Particles from the vesicle-enriched sample were washed as above, but in 1x PBS buffer. To remove any potential free DNA associated with the outside of the vesicle-enriched fraction, the sample was incubated with 2U of TURBO DNase (Invitrogen) at 37°C for 30 minutes. After a second round of TURBO DNase treatment as above, the enzyme was heat inactivated at 75°C for 15 minutes.

### DNA purification from tailed phage and vesicle enriched fractions

Lysis and DNA purification were performed in a 2 mL screwcap vials using the Qiagen Genomic-tip 20/G protocol, and buffers from the Qiagen Genomic DNA Buffer kit following manufacturer’s recommendations (Qiagen, Hildern, Germany). First, an RNase A solution (200 ug/mL) was prepared by adding 20 ul of 10 mg/ml RNase A to 1 ml of Buffer B1 (50mM Tris•HCl, pH 8; 50mM EDTA, pH 8; 0.5% Tween-20; 0.5% Triton X-100). Next, 1 mL of the Buffer B1 lysis buffer containing RNase A was added to 35 uL of viral concentrate. Then, 45 uL of a Proteinase K solution (20 mg/mL) prepared in sterile water was added, followed by the addition of 350 uL of Buffer B2 lysis buffer (3M guanidine HCl; 20% Tween 20). The samples were then incubated at 50°C for 120 minutes.

The resulting lysate was then loaded onto a Qiagen Genomic-tip 20/G column and purification was performed following manufacturers recommendations (Qiagen, Hildern, Germany). The Genomic-tip 20/G column was equilibrated with 1 mL of Buffer QBT equilibration buffer (750 mM NaCl; 50 mM MOPS pH 7.0; 15% isopropanol; 0.15% Triton X-100) by gravity flow. Sample lysate was then mixed by inverting several times and carefully pipetted sequentially onto an equilibrated Genomic-tip 20/G column, allowing the sample to enter the resin by gravity flow. Next, 1 mL Buffer B1 was combined with 350 uL Buffer B2, and the sample lysate tube was gently rinsed with 1 mL of this solution, and the rinse solution applied to the Genomic-tip 20/G column. The Genomic-tip 20/G column was washed by gravity flow by applying 1 mL of Buffer QC wash buffer (1.0 M NaCl; 50 mM MOPS, pH 7; 15% isopropanol) three times, in succession. Finally, the high molecular weight genomic DNA was eluted from the column by two successive applications of 1mL Buffer QF elution buffer (1.25 M NaCl; 50 mM Tris•HCl, pH 8.5; 15% isopropanol) pre-warmed to 50°C, resulting in purified DNA preparations in a 2 mL final volume.

The column purified DNA was concentrated by isopropanol precipitation as follows: The DNA eluant was split into two 2 mL conical screw cap tubes, 1mL per tube. The DNA was precipitated by adding 0.7 mL of room-temperature isopropanol per each 1 mL of DNA solution, followed by mixing by gentle inversion. After 2 hours at room temperature, the DNA was pelleted by centrifugation at 10,000 x g for 30 min at 4°C. The supernatant was removed, and the DNA precipitate washed by gentle addition of 1 mL of cold 70% ethanol and incubation for 60 seconds, followed by centrifugation at 10,000 x g for 15 min at 4°C. The supernatant was removed and the DNA pellet air dried for 5 min. The purified DNA pellets were resuspended in a final volume of 26 uL of 1X TE buffer (10 mM Tris•HCl, pH 8.0; I mM EDTA pH 8.0), and allowed to dissolve for a minimum of 10 min at room temperature, before final storage at 4°C. Final DNA quantity and quality was assessed initially by spectrophotometry and quantified via Quant-iT Picogreen dsDNA fluorometric assay (catalogue #P7589, Invitrogen). The virus-enriched sample collected from 440 L of 0.22um pre-filtered seawater collected at a depth of 25 m yielded a total of 5.1 μg of purified, high molecular weight DNA. The total yield for the corresponding vesicle fraction was 8.2 μg of purified, high molecular weight DNA.

### Oxford Nanopore sequencing methodology

Virus and vesicle enriched DNAs were processed using the Nanopore Ligation Sequencing Kit (LSK-109, Oxford Nanopore Technologies, Ltd.) following manufacturer’s instructions for the processing of high molecular weight DNA. A total of 2 virus enriched libraries were prepared using 2 and 1.5 ug of DNA each for the sequencing runs. All libraries were sequenced on a GridION X5 using FLO-MIN106 (R 9.4.1) flowcells (Oxford Nanopore Technologies, Ltd.). Read basecalls were generated from the signal traces using Guppy v3.0. The sequencing yield for the virus enriched fraction totaled 31 Gbp, generating reads with an N50 of 37.67 kb. The sequencing yield for the corresponding membrane vesicle-enriched fraction totaled 46 Gbp, with an N50 of 4.6 kbp.

### Concatemer detection

Repetitive long reads were identified as follows. In the first phase, each read over 5 kb was compared to itself using minimap2 (48) and lastal (49). Only forward strand self-hits of at least 500 bp (Fig. S1A) were retained from either method. In the second phase, if a read had three or more self-hits (a full-length central hit and at least one hit offset in each direction), a repeat size was calculated from the hit positions using two different methods.

The “clust” method first calculates the offset of each hit as the average of the difference between that hit’s start positions and the difference between its end positions (Fig. S1B). Next the “clust” method groups offsets using agglomerative clustering to allow for fragmented hits. Finally the median distance between neighboring offset groups is taken as the repeat size. The “fft” method starts with hit offsets, smooths their locations with kernel density estimation (Fig. S1C), uses fast fourier transform to find the dominant frequency, and takes the corresponding wavelength as the repeat size.

For all four combinations of the two methods and two search tools, the calculated repeat size was compared to the actual hits. If the hits covered at least 50% of the expected single offset self-hit (red line in Fig. S1.A), and the repeat size was less than 60% of the total read length, then the read was marked as a concatemer by that method. If a read was marked as concatemeric by any method, then it was considered a concatemer using the median repeat size of all methods that flagged it (Dataset S2). Code for finding concatemers and calculating repeat size is available at: https://github.com/jmeppley/concatemer_finding.

### VEIME generation

Polished VEIME sequences (Dataset S3) were generated from concatemeric reads as follows. Each read was broken up into non-overlapping repeat-sized fragments. All fragments were compared to each other with lastal (49), retaining hits of at least 80% of the fragment length. Fragments from different reads were merged into one pool for polishing, if the majority of the fragments from each read were highly similar (> 80% sequence similarity over > 80% of their lengths). The consensus sequence of each pool of fragments was determined using 3 passes of racon (50) with a final pass of medaka (https://github.com/nanoporetech/medaka), following the assembly free viral genome (AFVG) polishing methods as previously described (19). The consensus sequences were further polished using two passes of racon with Illumina short reads derived from the same sample, as previously described (19). Briefly, genomic DNA from each sample was sheared to an average size of 350 bp using a Covaris M220 Focused-ultrasonicator (Covaris, Woburn, MA) with Micro AFA fiber tubes (Covaris, #520166, Woburn, MA). Libraries were sequenced using a 150 bp paired-end NextSeq High Output V2 reagent kit (Illumina, FC-404-2004, San Diego, CA). Finally, to improve gene predictions, erroneous frameshifts were corrected where possible using proovframe (51) and genes retrieved from GTDB version 95 (52).

### Bipartite Network Modules

Genomic sequences from the VEIMEs – including the VEIME-like monomers derived from the membrane vesicle fraction - were pooled with previously published phage satellites (see “Reference Sequences” below), and assembled into a bipartite network and partitioned into modules (Dataset S3, S5, and S6) as follows. Genes predicted using prodigal (53) were annotated independently with both EggNOG (54) using eggnog_mapper (55) and VOGdb using hmmsearch (56). Details of tools used, including versions and parameters are provided in Dataset S1.

A bipartite network (Fig. 2) was defined with satellites as one set of nodes and gene families as the other, with an edge between a satellite and gene family if that satellite includes a gene annotated with that gene family. The bipartite network comprised two connected components. The smaller component, containing 3 VEIMEs, was dubbed module 1, and the larger was partitioned into modules 2 through 12 using the constant Potts model (57) as implemented by the leidenalg Python package (Dataset S1). For most analyses, modules 1 and 12, each containing fewer than 5 sequences, were ignored.

### Reference Sequences

Polished VEIMEs were searched for in three databases of marine microbial genomes, GTDB (58), GORG (59), and Mar. Micro. DB (60) using minimap2 (Dataset S1). Only hits covering at least 80% of the VEIME sequence were kept (Fig. S3). The taxonomy of the best hit was assumed to be the host taxonomy for the VEIME (Dataset S3).

To provide context for the concatemer-derived mobile element sequences, previously published phage satellite sequences and viral genomes were downloaded from NCBI. Phage satellite references include PICIs listed in (2, 6, 44, 61). Reference phage genomes were chosen for tailed phage with complete genomes and known bacterial hosts. In all 90 satellites and 1700 viral genomes were downloaded (Dataset S4).

Additionally, 16 monomers, taken from previously published concatemeric <u>a</u>ssembly-<u>f</u>ree <u>p</u>utative <u>P</u>ICIs (AFPPs; (*22*)) were included in the analyses (Dataset S3). Similar to the VEIMEs, these were generated from concatemeric Nanopore reads sequenced from 0.2 um filtered, TFF concentrated seawater collected at Station ALOHA.

Finally, long reads with direct terminal repeats (DTRs) were gathered from the tailed phage fraction and self polished into assembly-free viral genomes (AFVGs) using DTR-phage-pipeline (22). These were used to provide a distribution of phage genome sizes in this environment (Fig 1B), and a set of environmental terminase small-subunit genes for comparison with VEIMEs (Fig. S7).

### Environmental Metagenome Abundances

The historical presence of VEIME sequences in station ALOHA waters was assessed as follows. Raw metagenomic reads were collected from 456 metagenomic samples collected from station ALOHA between 2014 to 2018 at depths ranging between 5m and 500m deep (Dataset S7) and assembled into contigs as described previously (27). VEIME sequences were mapped against contigs using minimap2, and hits covering at least 80% of the VEIME were kept. Raw reads were mapped against contigs using BWA mem (62) and used to calculate base-by-base coverage of contigs at VEIME hit locations. The mean contig coverage across the hit location was normalized to the number of sequenced reads and used as a proxy for environmental VEIME abundance.

Additionally, the global distribution of VEIMEs was assessed using TARA GOV 2.0 reads and assemblies (25). Assembled GOV contigs were downloaded from iVirus and matched by sampling station and depth to raw reads obtained from NCBI (Dataset S7). VEIME abundance in these samples was calculated as for the Station ALOHA metagenomes.

### Phylogenetic trees

Phylogenetic trees were inferred for tyrosine integrases using the VOG00035 hmm (Fig. 3; Fig. S4), for serine recombinases using VOG00893 (Fig. S6), and for terminase small subunits using VOG06274 (Fig. S7). For each selected gene annotation, genes from VEIMEs and reference sequences were aligned individually using hmmalign (56) and combined into a single multiple sequence alignment (MSA) for each gene. MSAs were trimmed of columns present in fewer than half of the genomes, and genomes containing fewer than 5% of the amino acids were removed from the alignments. Trees were inferred from MSAs using IQ-TREE (63) with the partitioned LG+GAMMA model (64).

## Acknowledgments

We thank the captains and crews of R/V Kilo Moana the HOT program for cruise support and oceanographic data acquisition. We thank Thomas Hackl for supplying tycheposon integrase gene alignments. This work is a contribution of the Simons Collaboration on Ocean Processes and Ecology, and the Center for Microbial Oceanography: Research and Education (C-MORE). This work was supported by: Simons Foundation grant 329108 (EFD); Simons Foundation grant 721223, (EFD); Gordon and Betty Moore grant 3777, (EFD); National Science Foundation OCE-2049004 (SJB); Simons Foundation Early Career Investigator grant (SJB)

