## Supplementary material for "Marine viral particles reveal an expansive repertoire of phage-parasitizing mobile elements": Supplment_Figures

**This PDF file includes:**

Figures S1 to S8
Legends for Datasets S1 to S7

**Other supporting materials for this manuscript include the following:**

Datasets S1 to S7

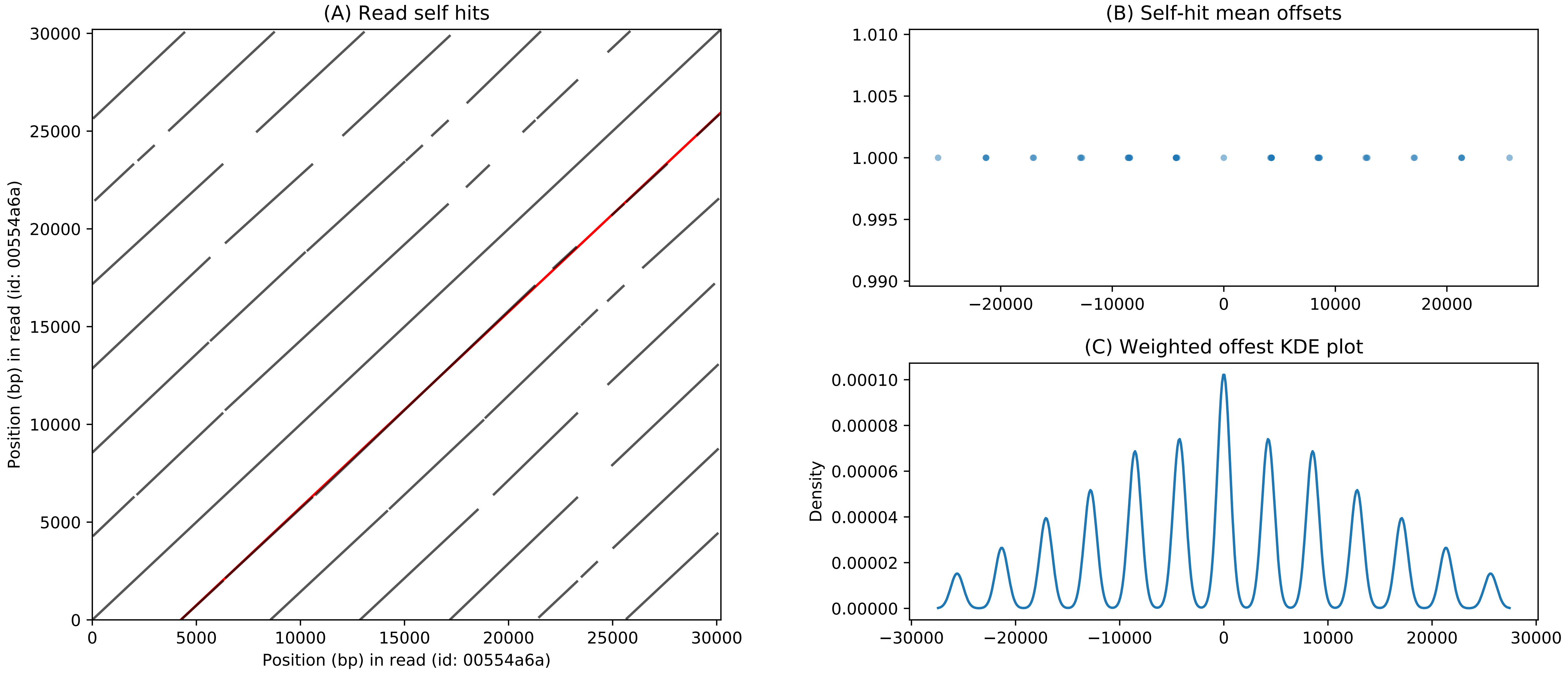


Figure S1: Concatemeric Read Detection
Self hits, shown as a dotplot in (A), are grouped by offset (B) and used to infer the repeat size by clustering or by fast-fourier-transform of their KDE (C). The calculated repeat size is evaluated by comparing the theoretical alrgest self alignment (red line in (A)) to the actual hits.

### Figure S2: VEIME Genome Maps and Annotations Organized by Modules

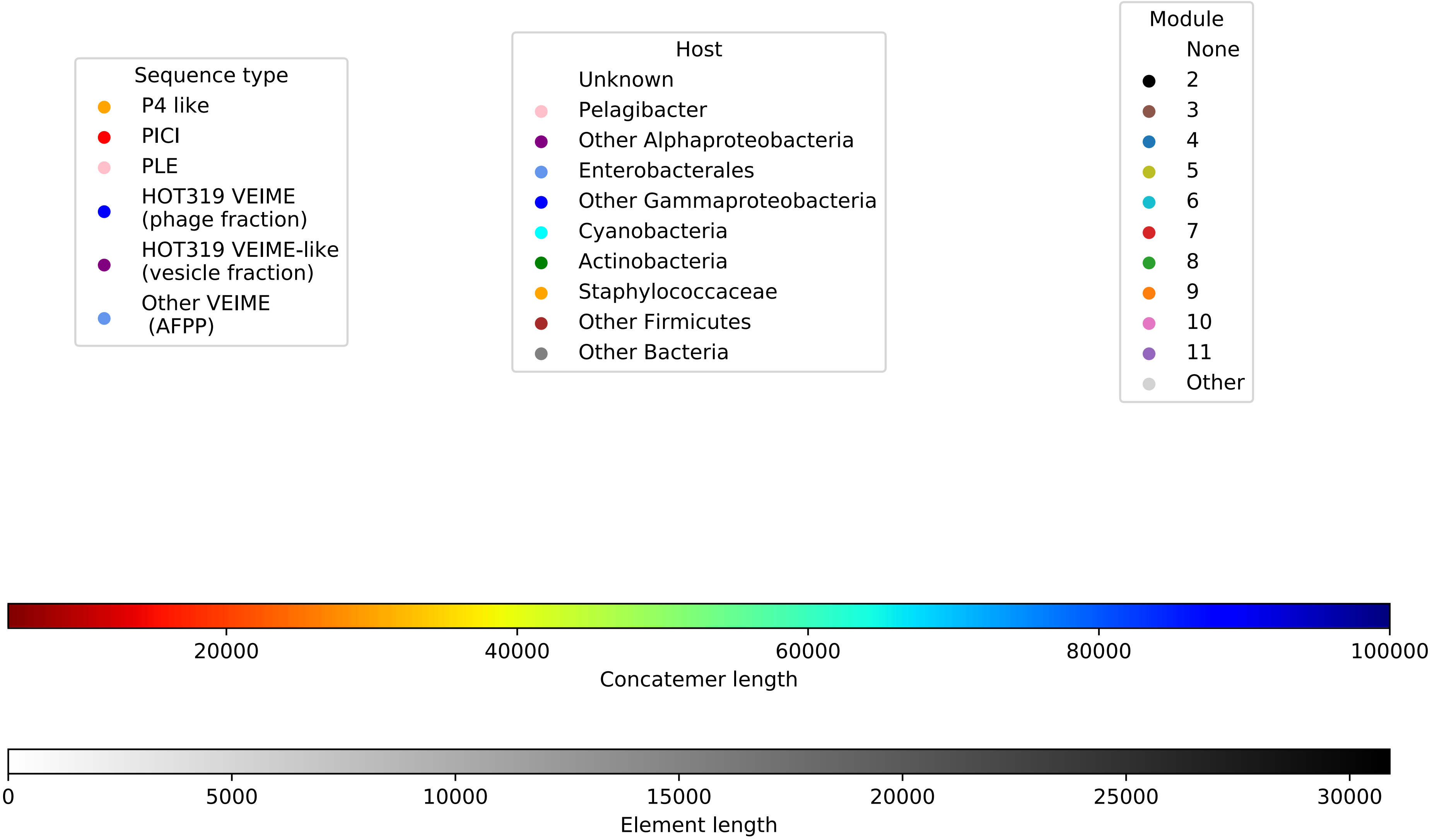


Figure S2: Module Genome Maps
With one page per module, each page shows the gene annotations for each phage satellite in that module (lower left), arranged by hierarchical clustering based on shared gene conteng (lower right panel). The lower center panel displays genome metadata (legend above). The top panel summarizes the gene positions across all satellite genomes.

Figure S2: VEIME Genome Maps & Annotations Organized by Modules continued

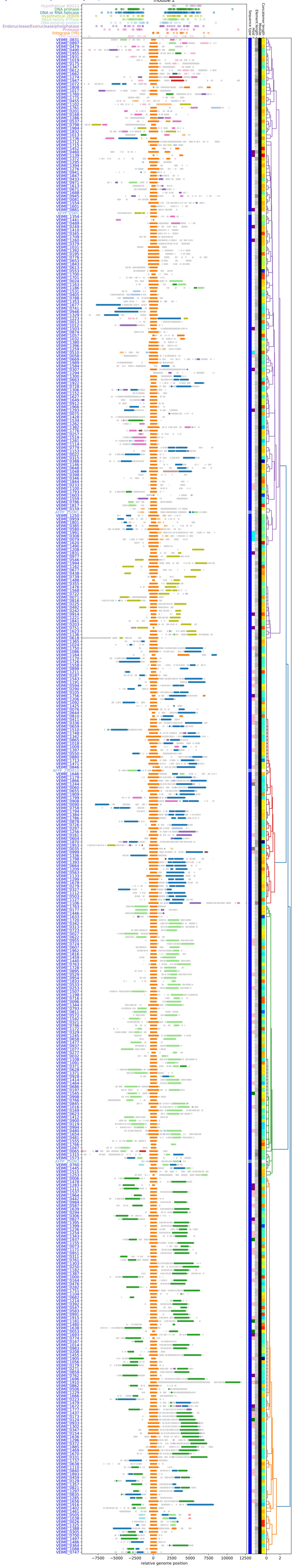

**Figure S2: VEIME Genome Maps & Annotations Organized by Modules continued**

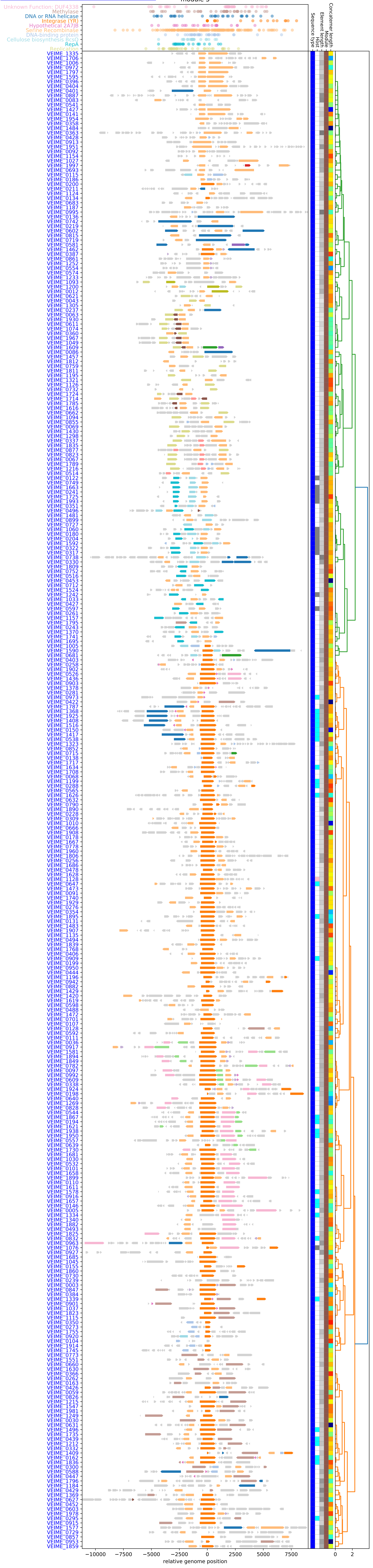

**Figure S2: VEIME Genome Maps & Annotations Organized by Modules continued**

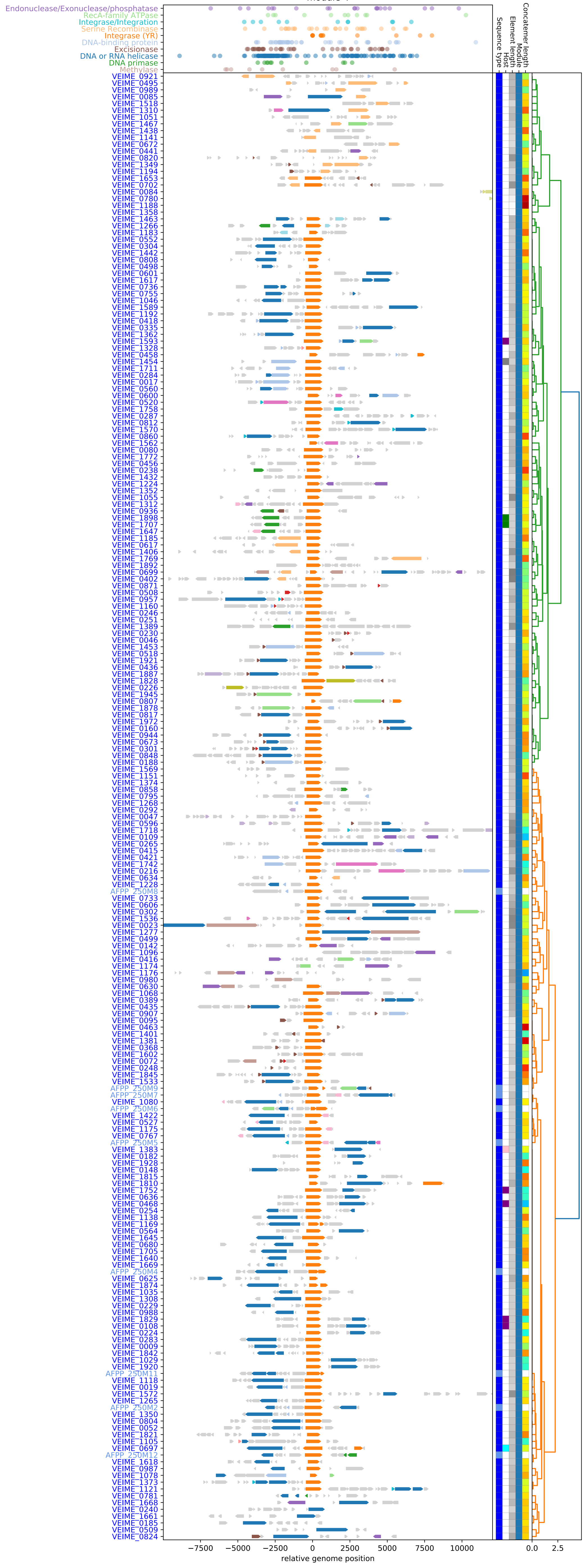

**Figure S2: VEIME Genome Maps & Annotations Organized by Modules, cont'd**

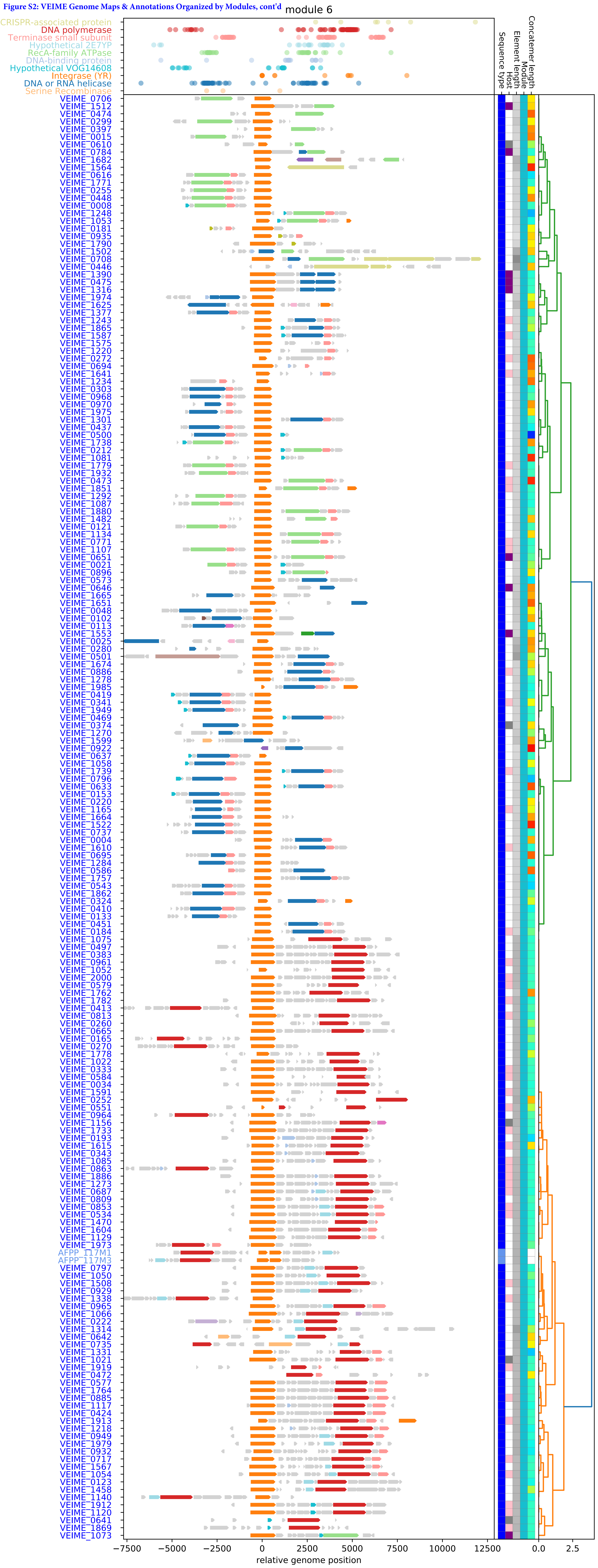

**Figure S2: VEIME Genome Maps & Annotations Organized by Modules continued**

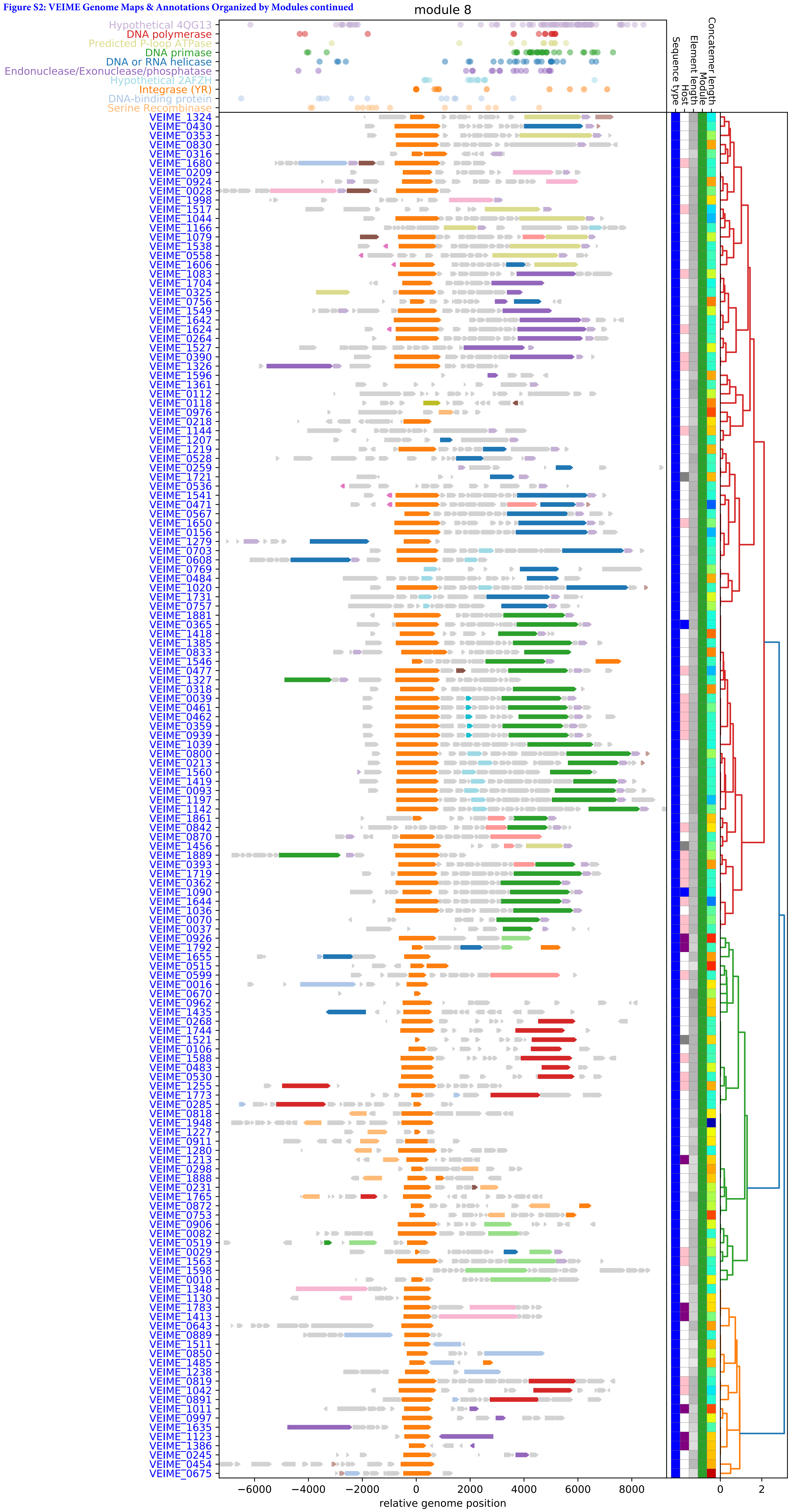

**Figure S2: VEIME Genome Maps & Annotations Organized by Modules continued**

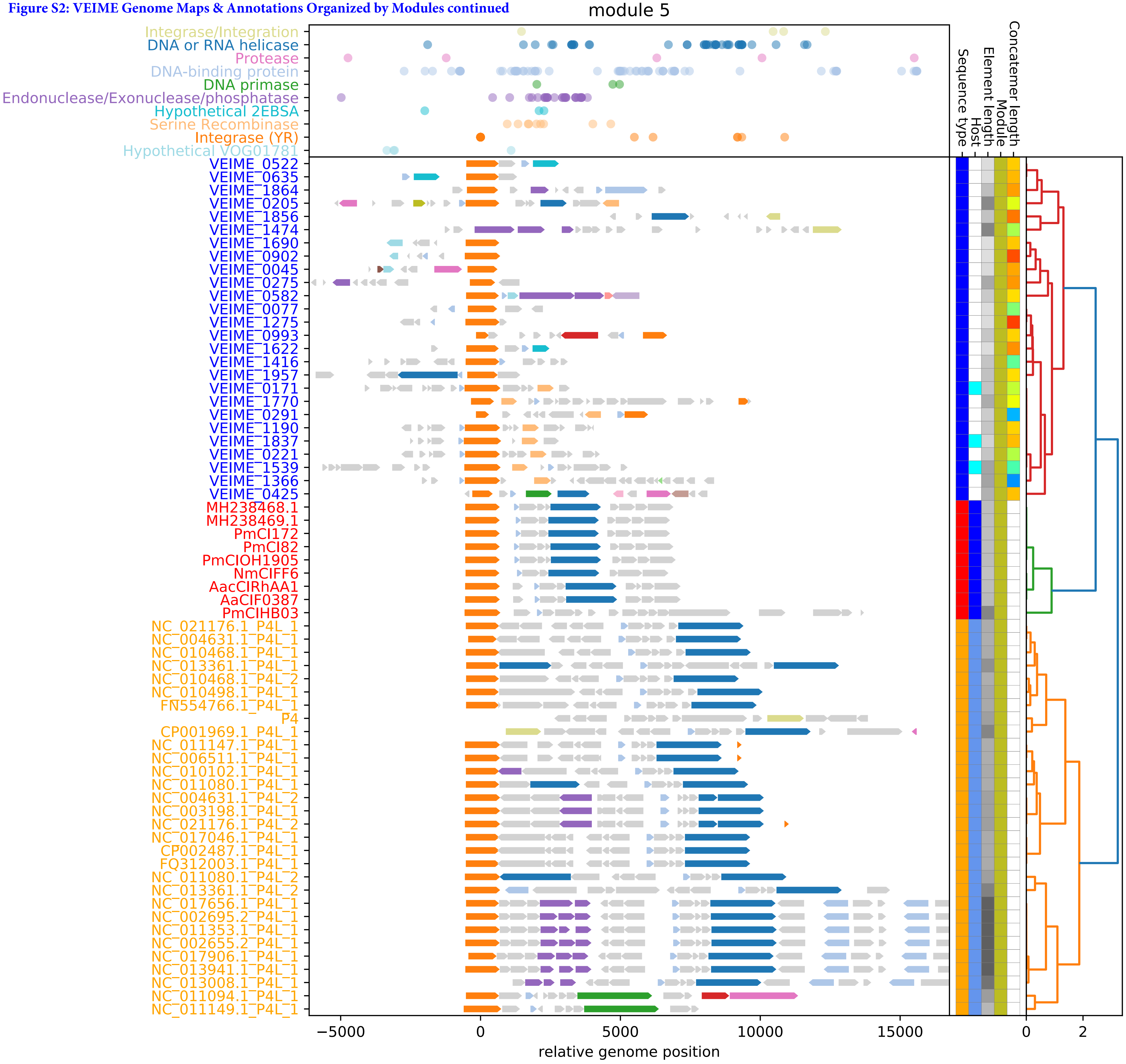

Figure S2: VEIME Genome Maps & Annotations Organized by Modules continued

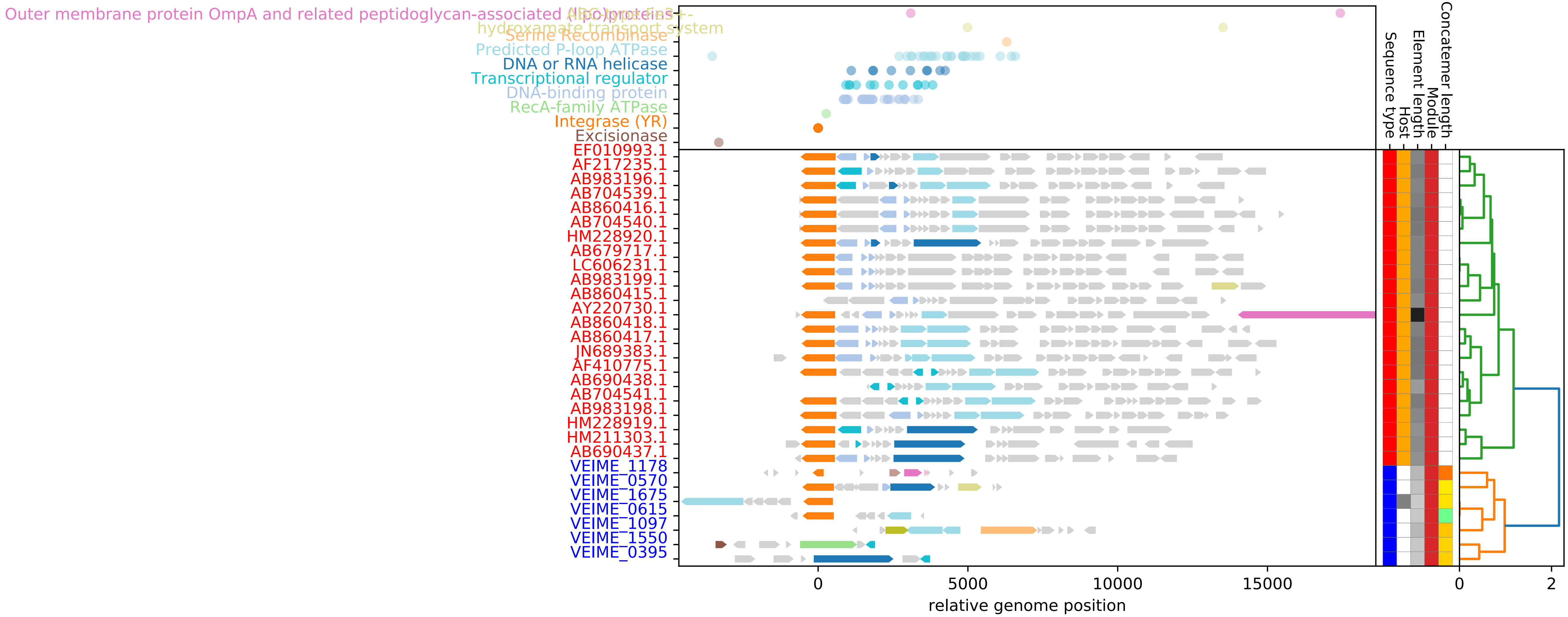

**Figure S2: VEIME Genome Maps & Annotations Organized by Modules continued**

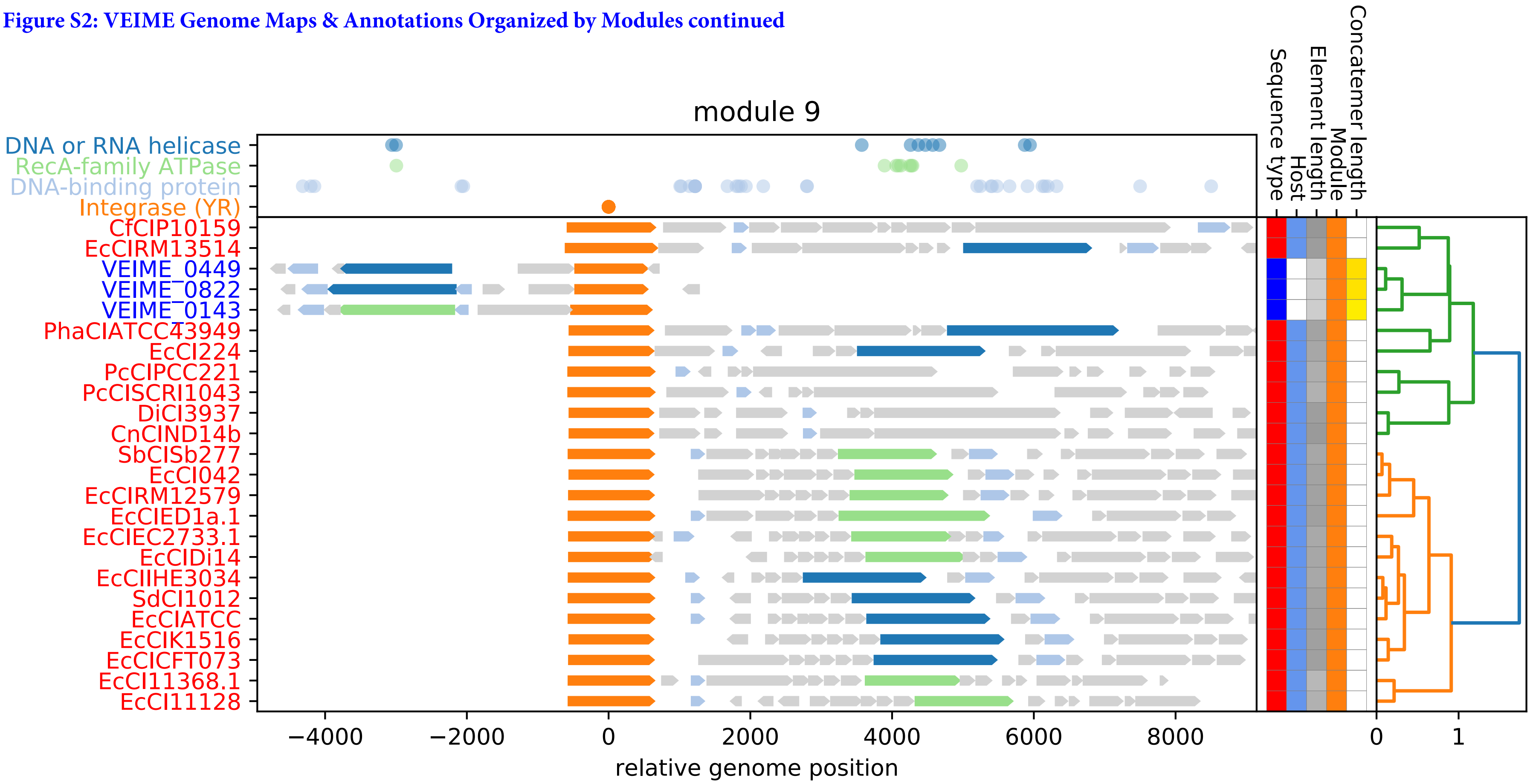

**Figure S2: VEIME Genome Maps & Annotations Organized by Modules continued**

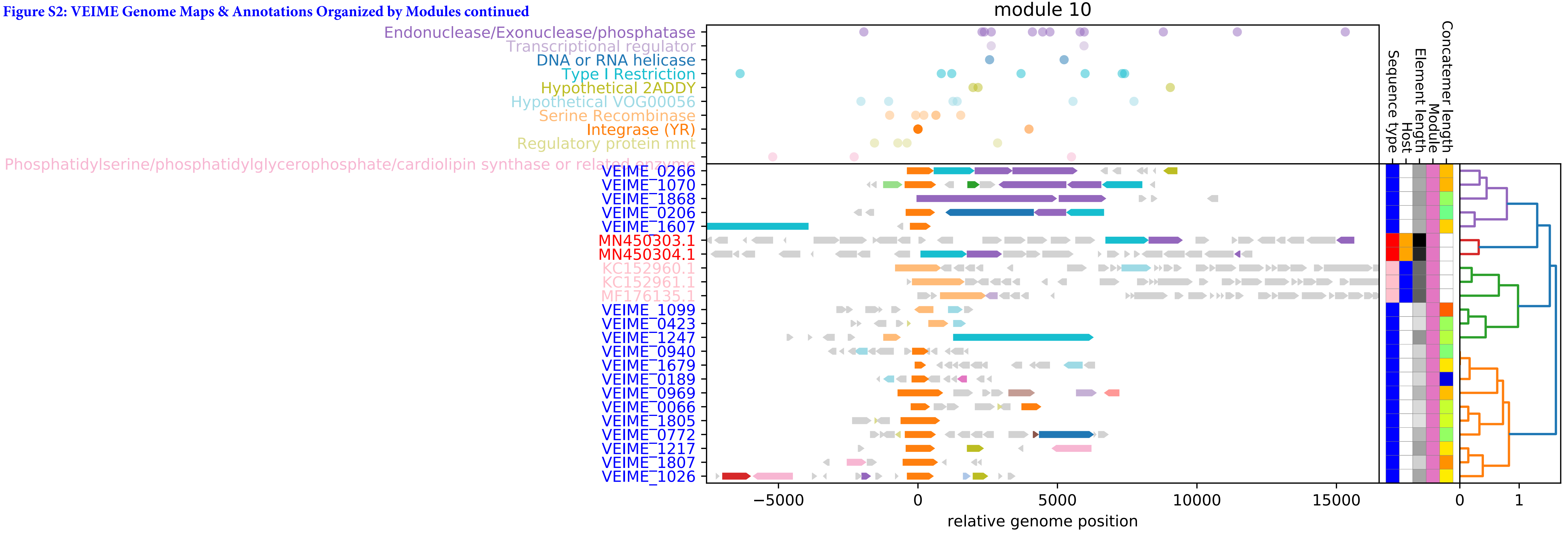

**Figure S2: VEIME Genome Maps and Annotations by Module continued**

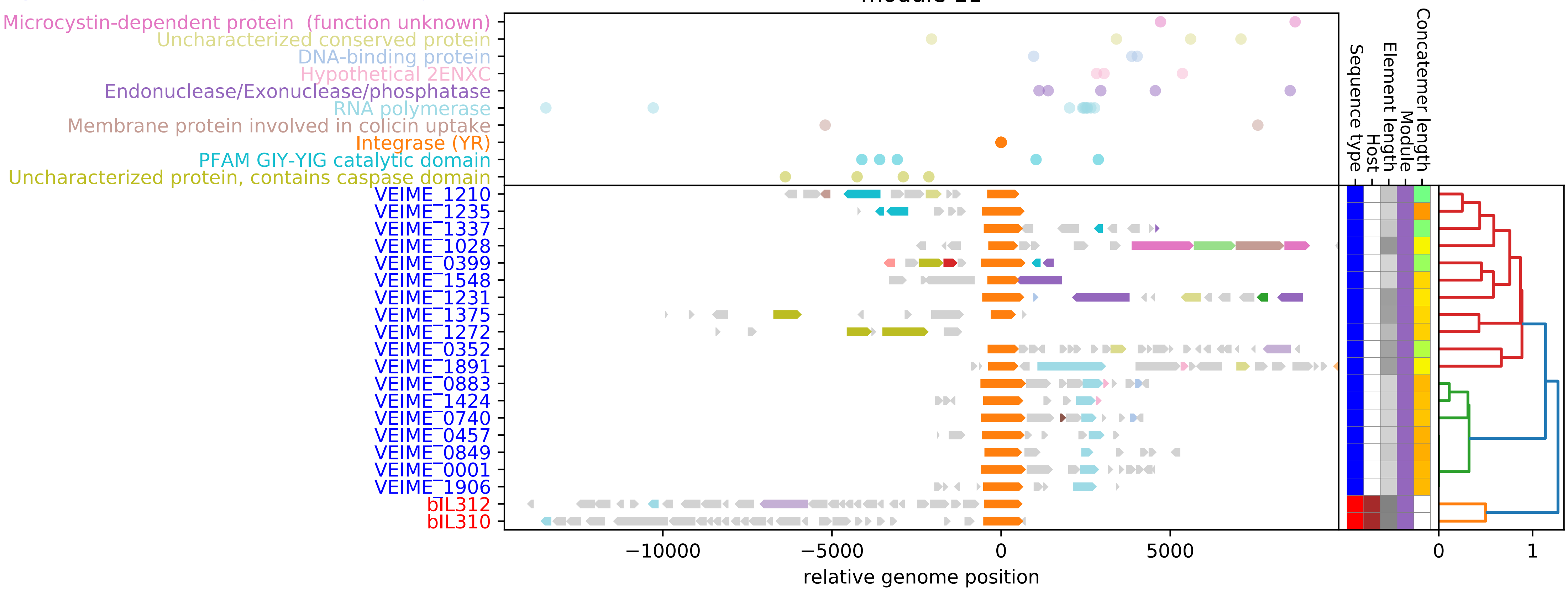

Figure S2: VEIME Genome Maps & Annotations Organized by Modules continued

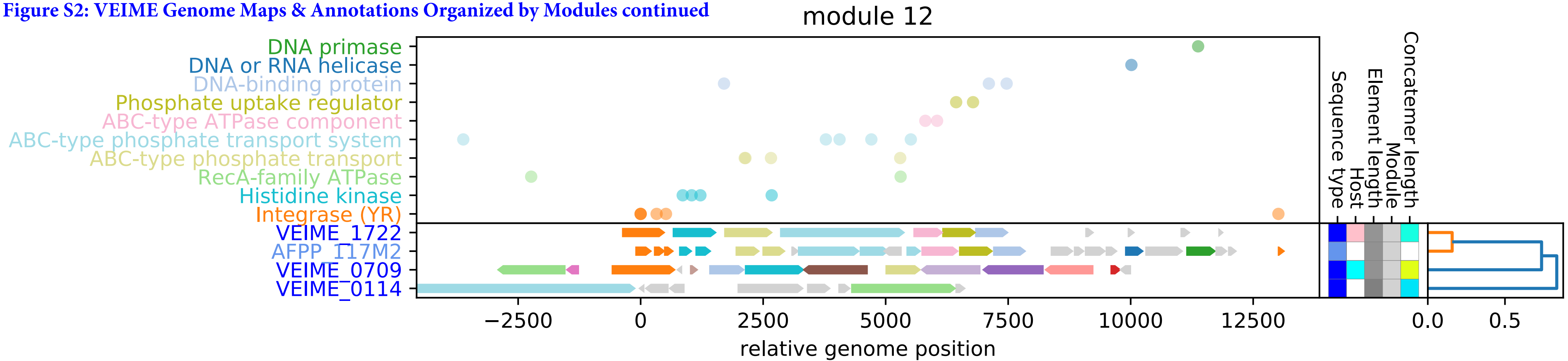

**Figure S2: VEIME Genome Maps & Annotations Organized by Modules continued**

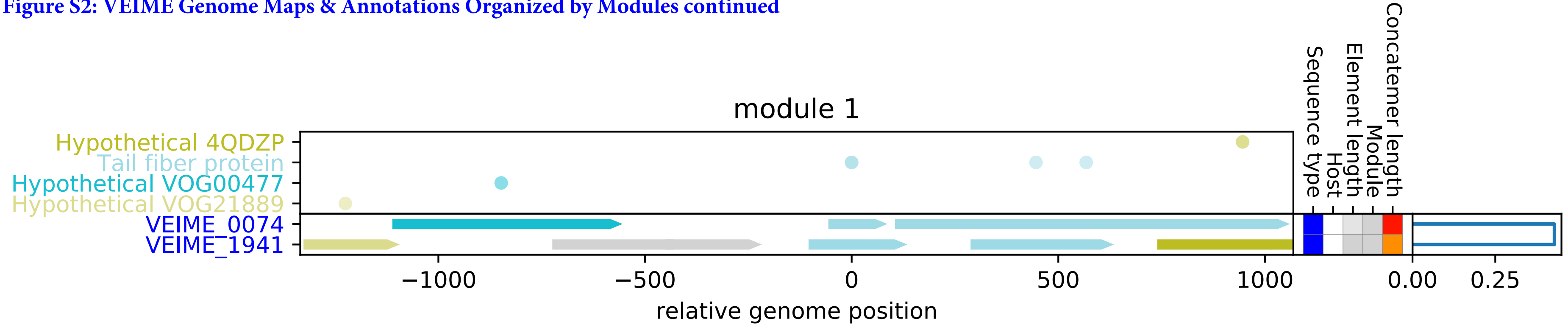

Figure S3: Genome maps of VEIMEs to microbial genomic contigs. VEIMEs matching contigs at least 50kb in length over at least 90% of the VEIME length are plotted against the genomic context of their match. Homology between VEIME and genomic genes is indicated by connecting parallelograms shaded by AA %ID (scale below). Short regions of genomic self similarity are indicated with arcs colored by nucleotide identity using the same %ID color scale.

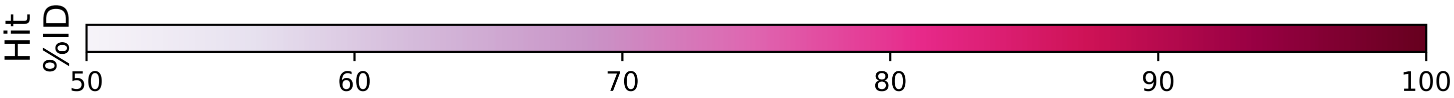

**Fig. S3.VEIME_genome_mappings cont'd.**

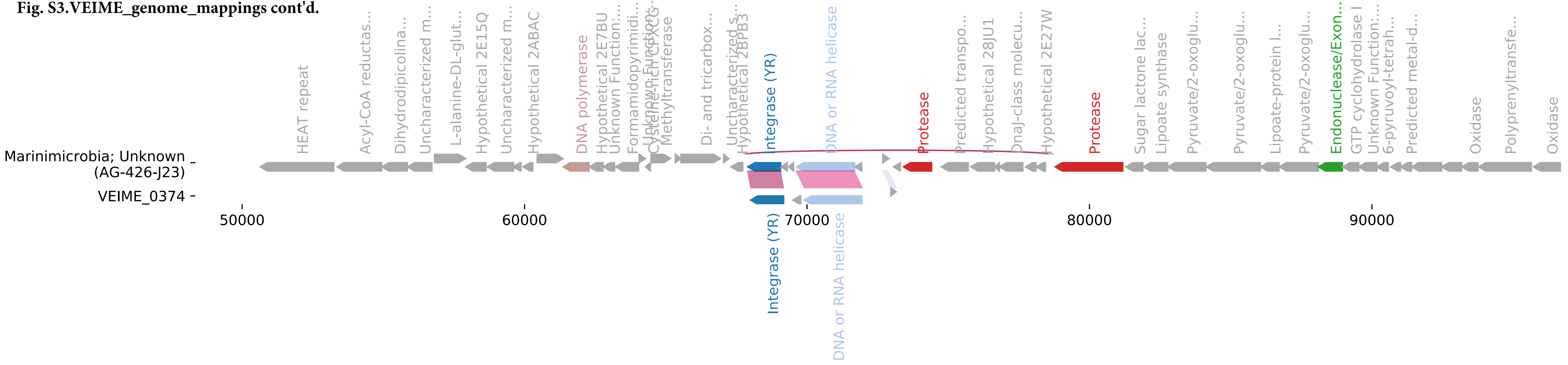

**Fig. S3.VEIME_genome_mappings cont'd.**

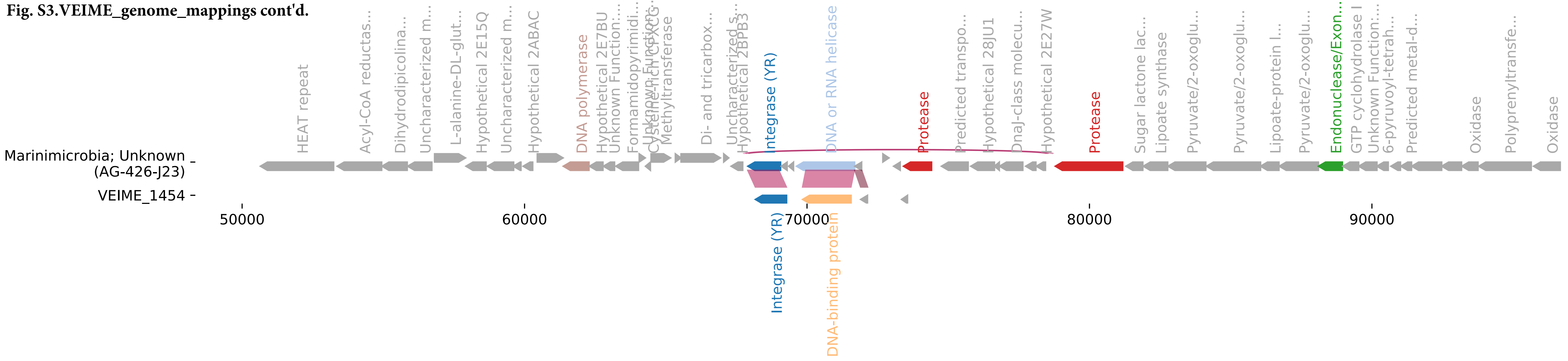

**Fig. S3.VEIME_genome_mappings cont'd.**

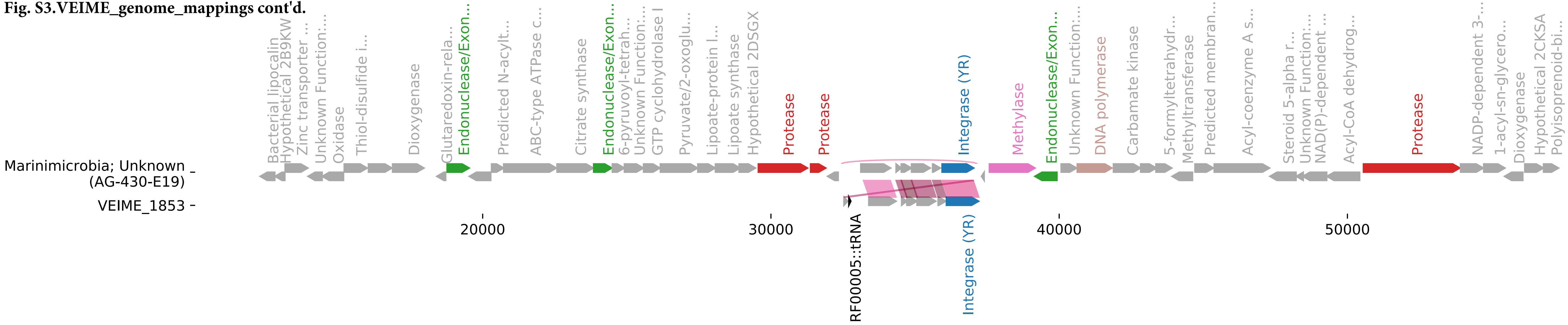

**Fig. S3.VEIME_genome_mappings cont'd.**

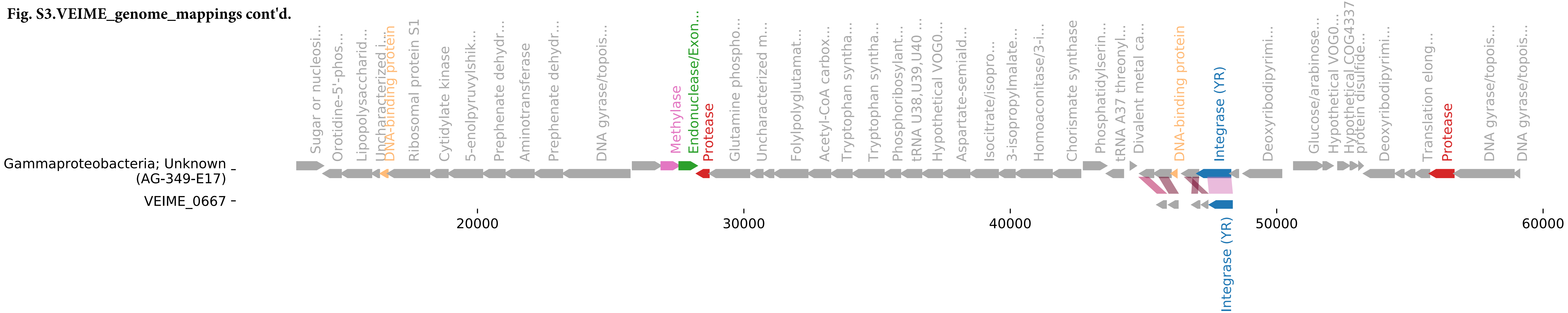

Fig. S3.VEIME_genome_mappings cont'd.

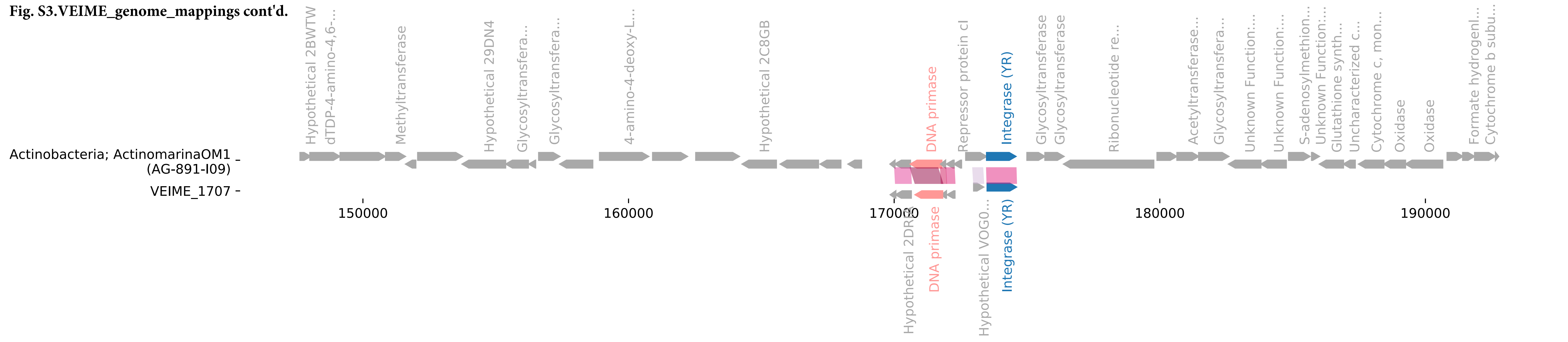

Fig. S3.VEIME_genome_mappings cont'd.

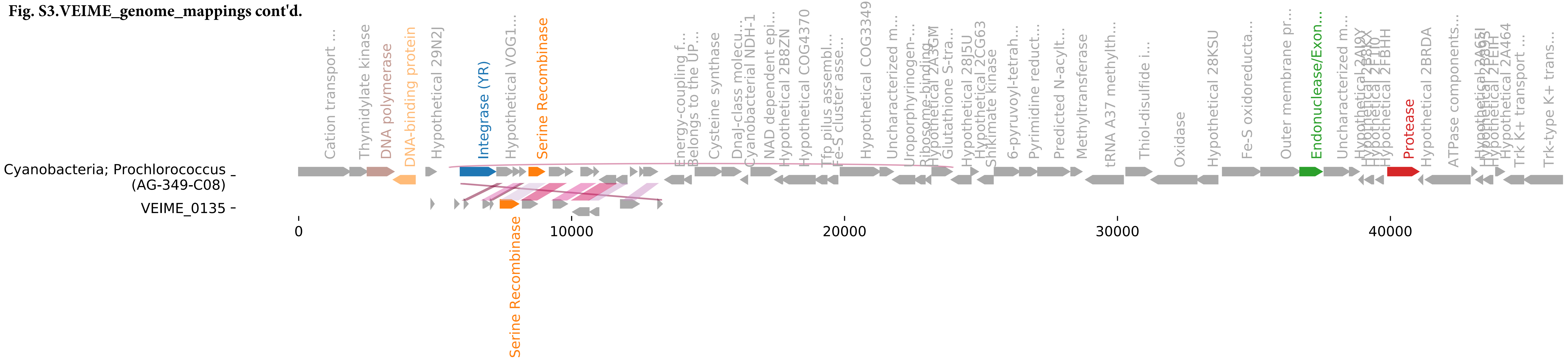

**Fig. S3.VEIME_genome_mappings cont'd.**

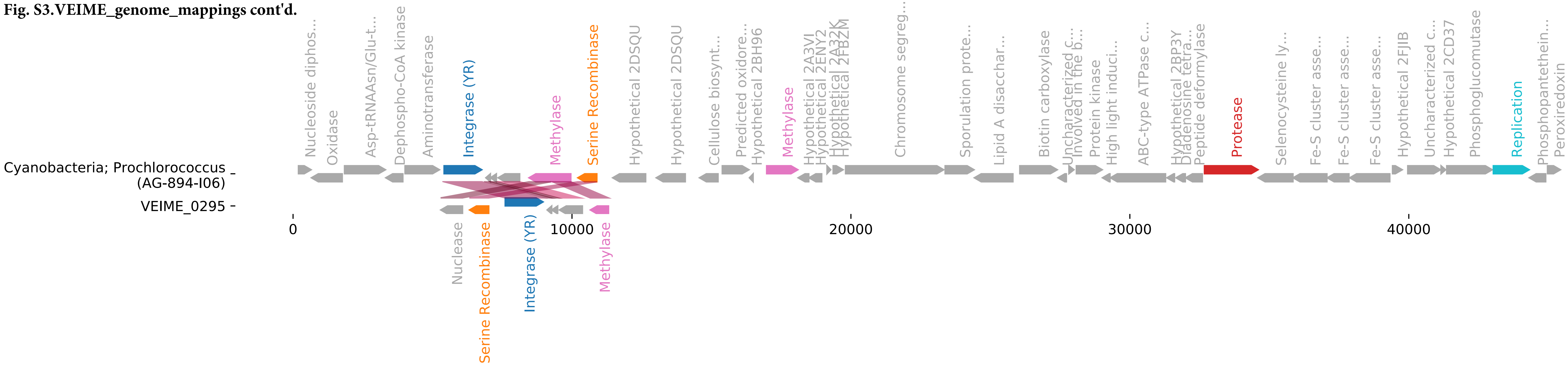

**Fig. S3.VEIME_genome_mappings cont'd.**

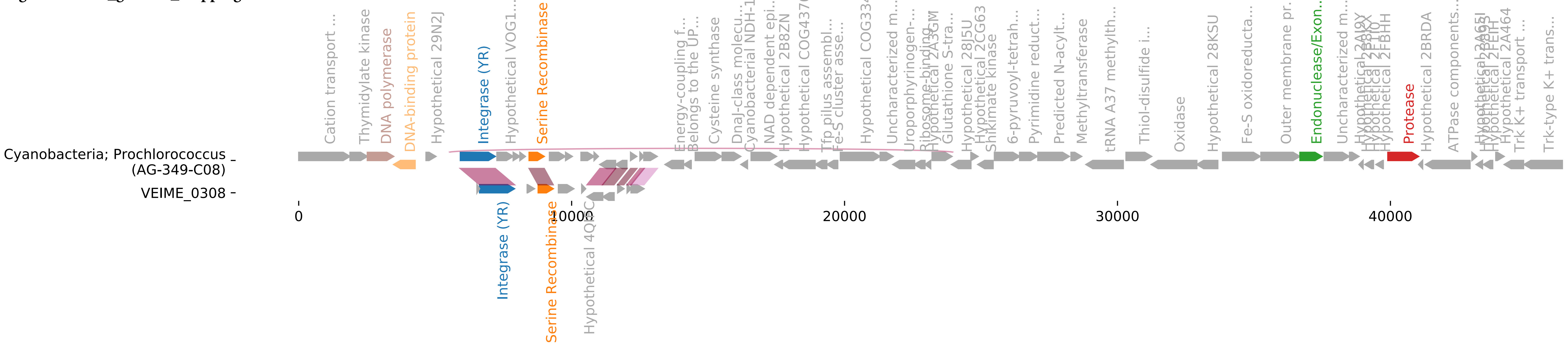

**Fig. S3.VEIME_genome_mappings cont'd.**

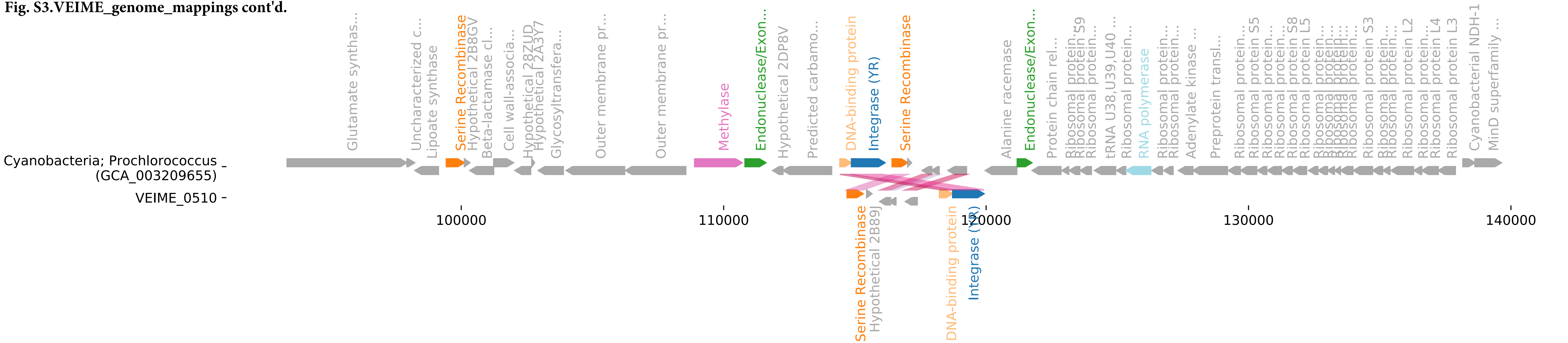

**Fig. S3.VEIME_genome_mappings cont'd.**

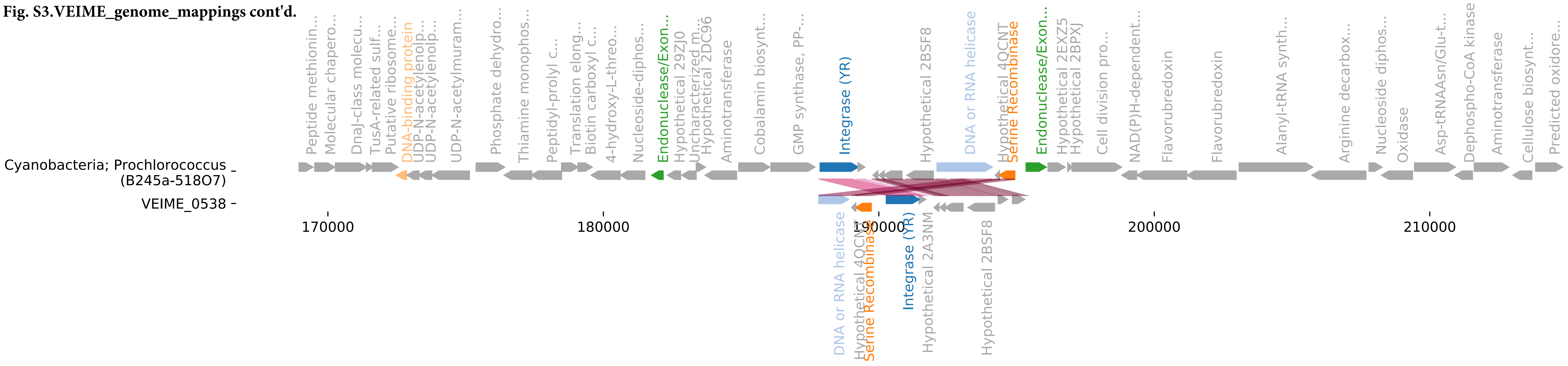

**Fig. S3.VEIME_genome_mappings cont'd.**

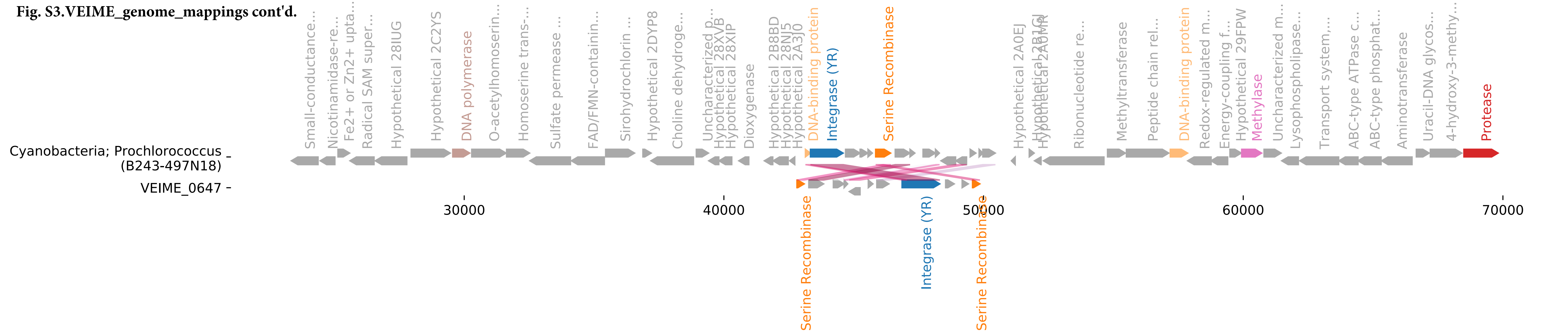

**Fig. S3.VEIME_genome_mappings cont'd.**

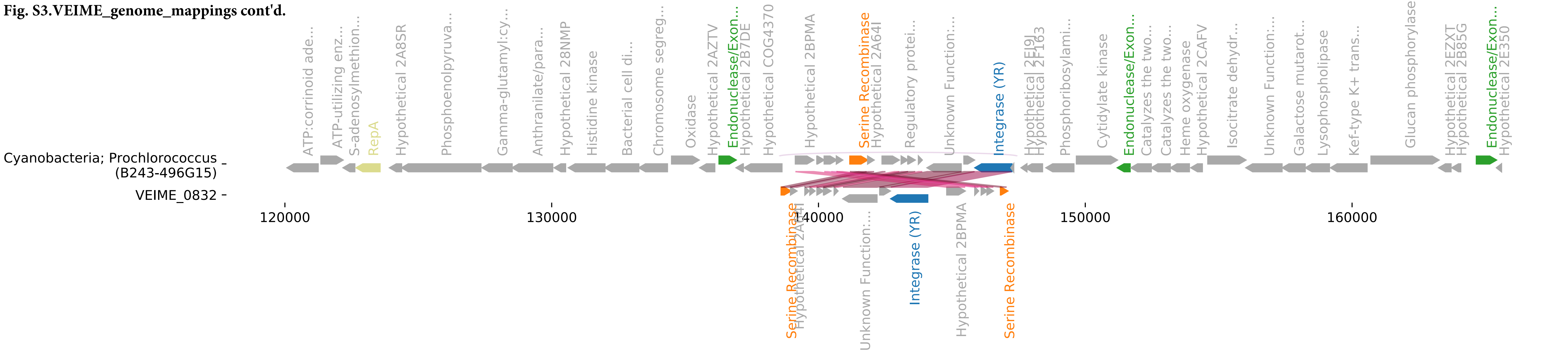

**Fig. S3.VEIME_genome_mappings cont'd.**

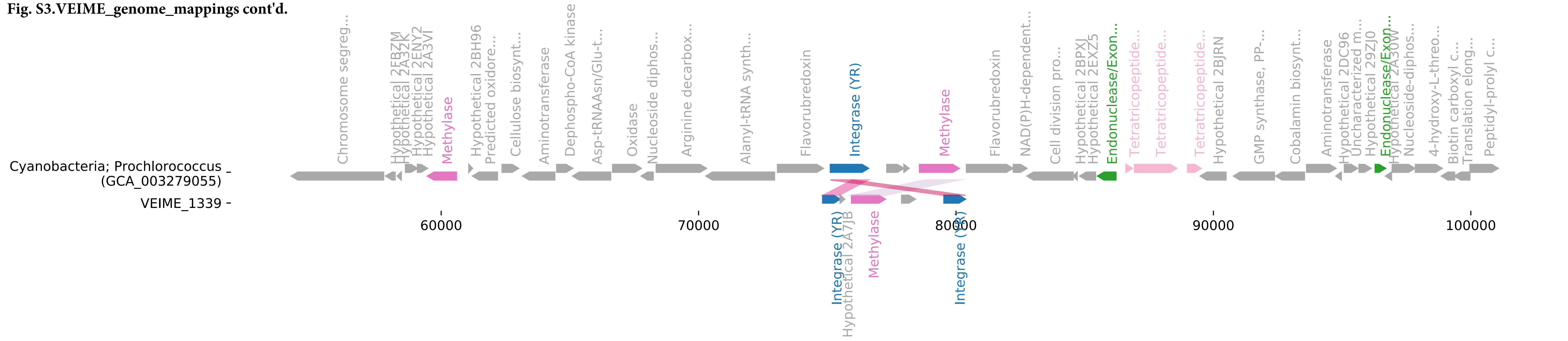

Fig. S3.VEIME_genome_mappings cont'd.

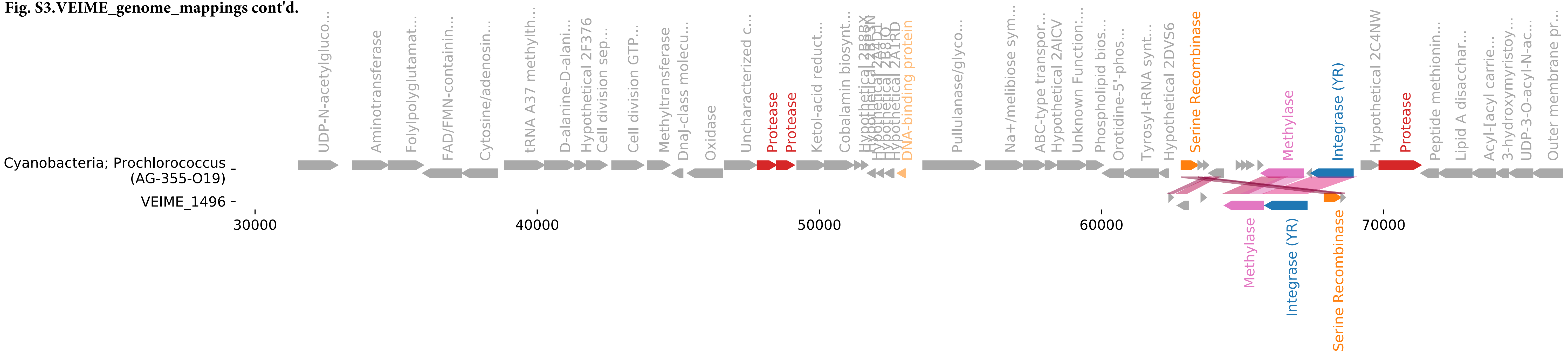

**Fig. S3.VEIME_genome_mappings cont'd.**

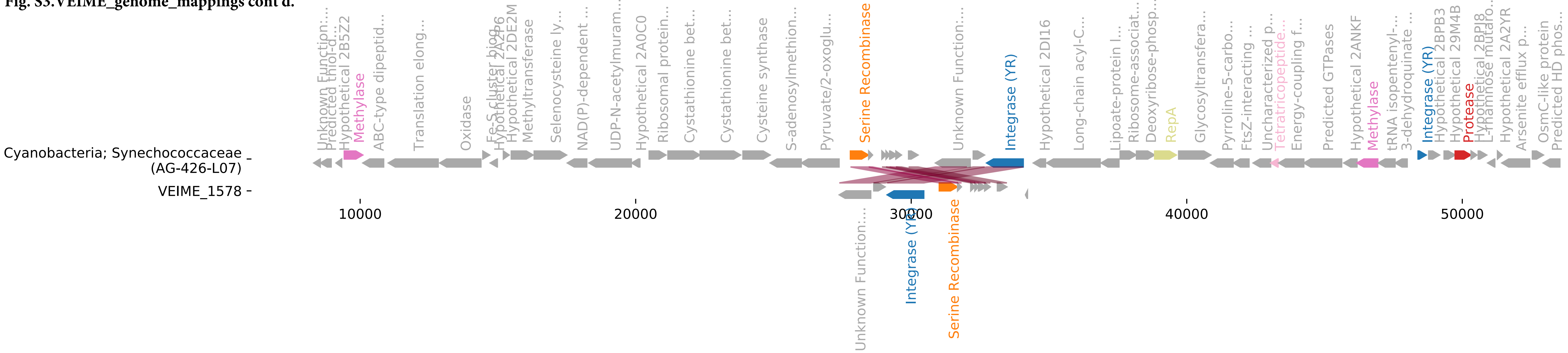

Fig. S3.VEIME_genome_mappings cont'd.

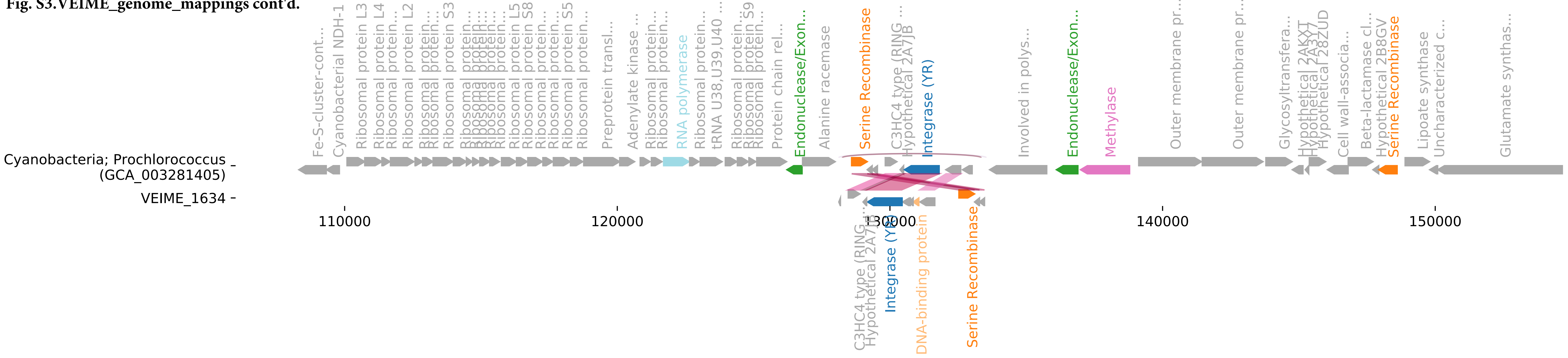

**Fig. S3.VEIME_genome_mappings cont'd.**

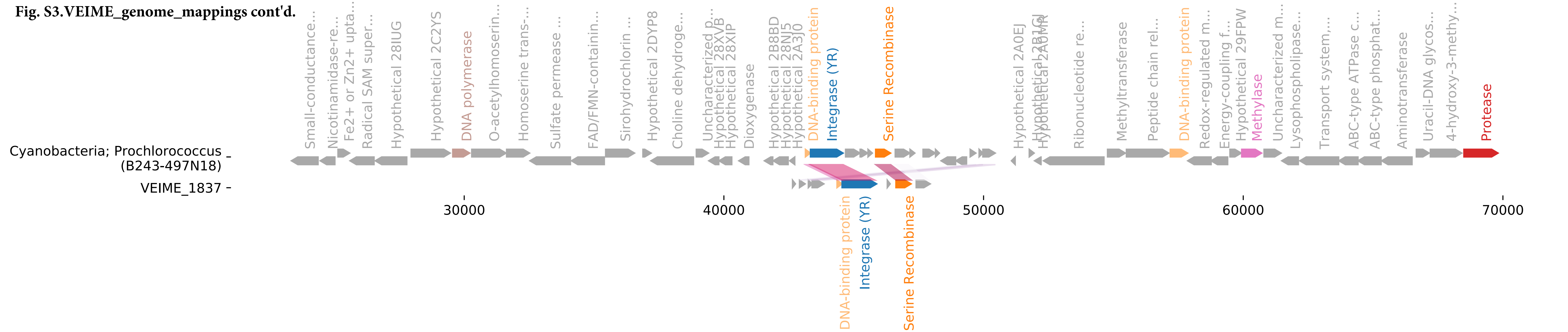

**Fig. S3.VEIME_genome_mappings cont'd.**

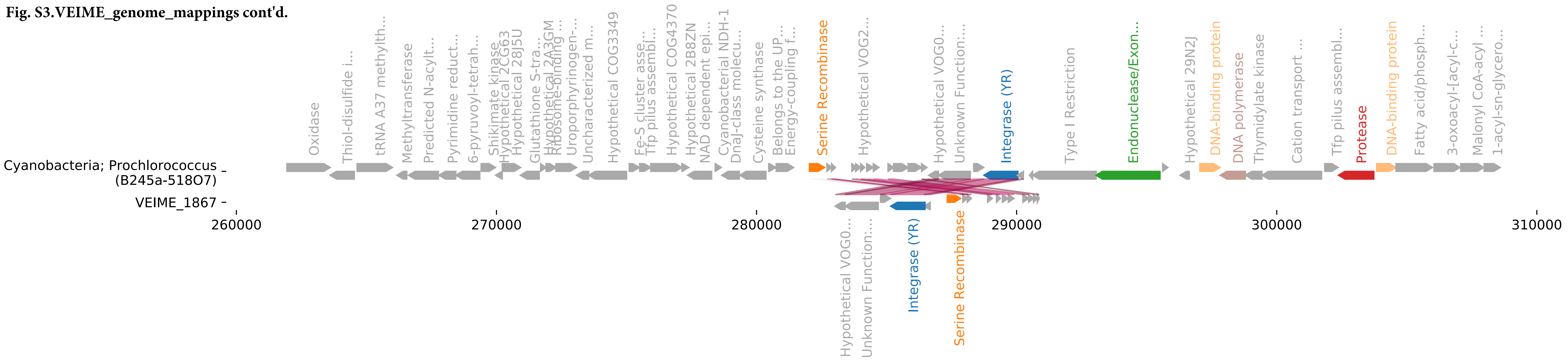

**Fig. S3.VEIME_genome_mappings cont'd.**

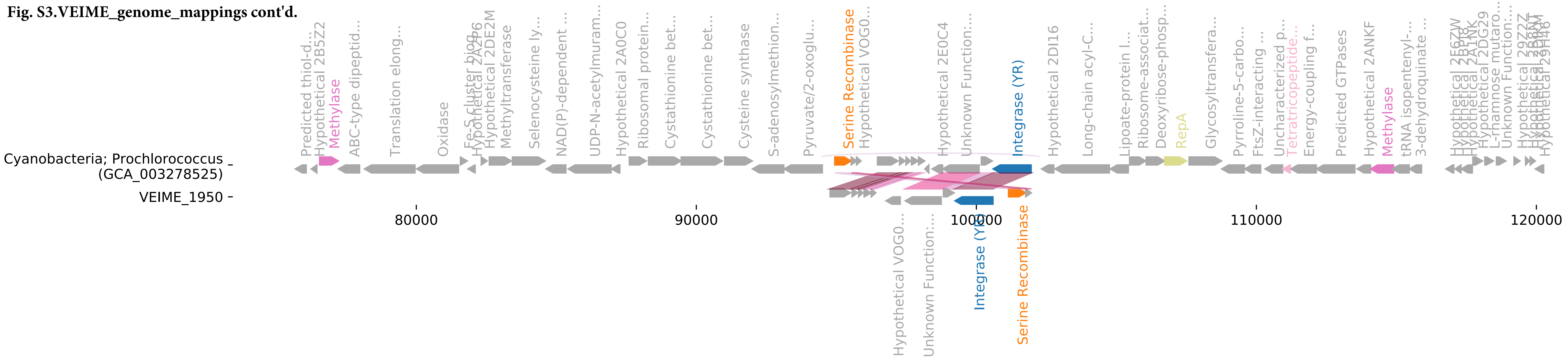

**Fig. S3.VEIME_genome_mappings cont'd.**

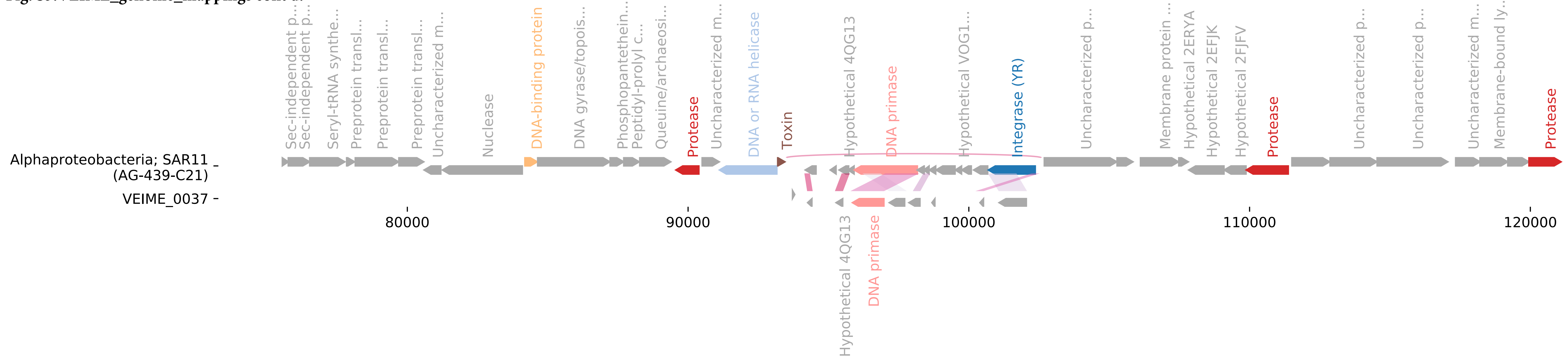

**Fig. S3.VEIME_genome_mappings cont'd.**

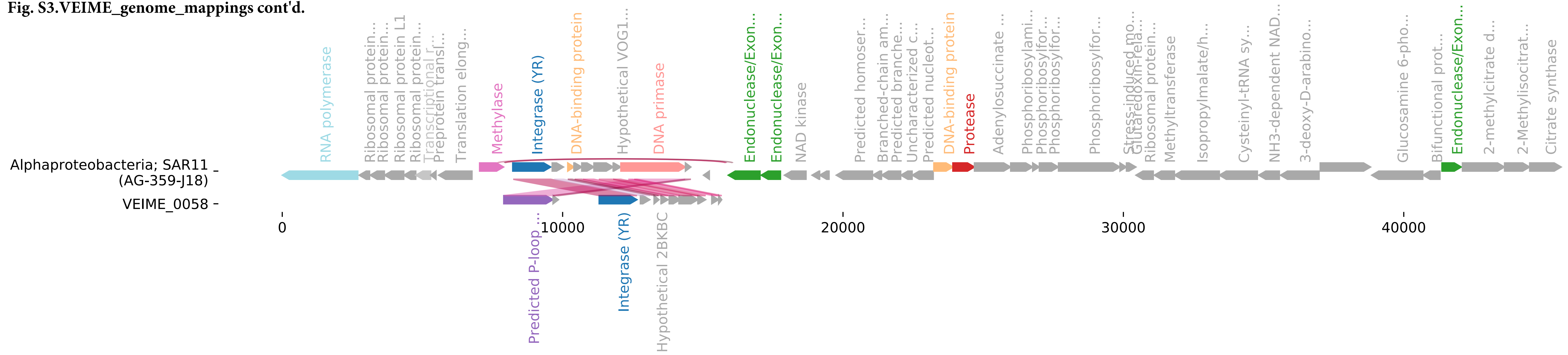

**Fig. S3.VEIME_genome_mappings cont'd.**

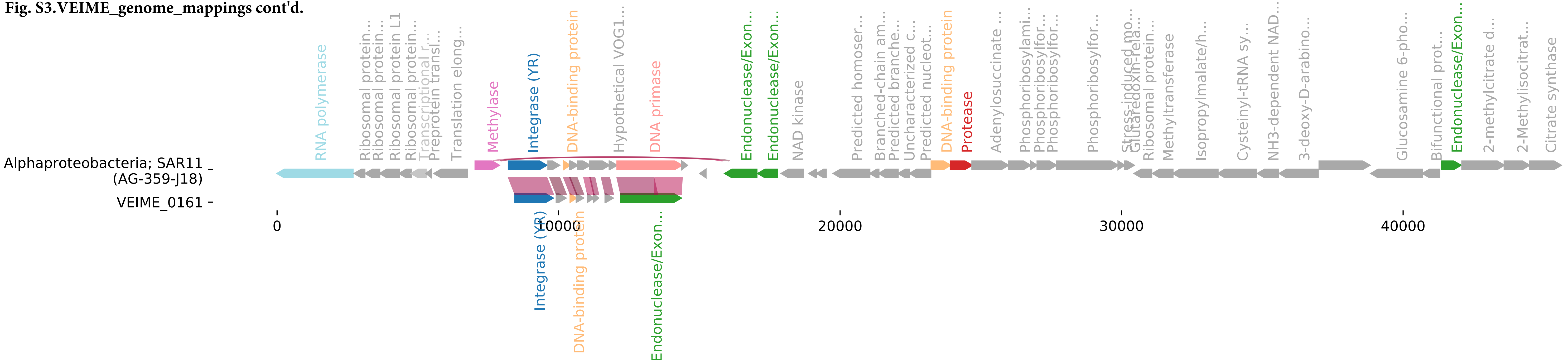

Fig. S3.VEIME_genome_mappings cont'd.

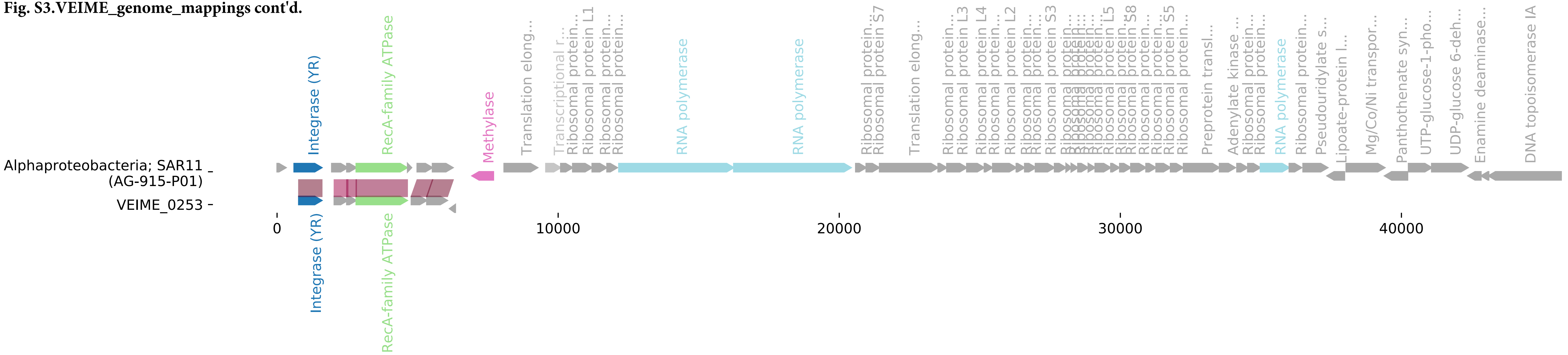

Fig. S3.VEIME_genome_mappings cont'd.

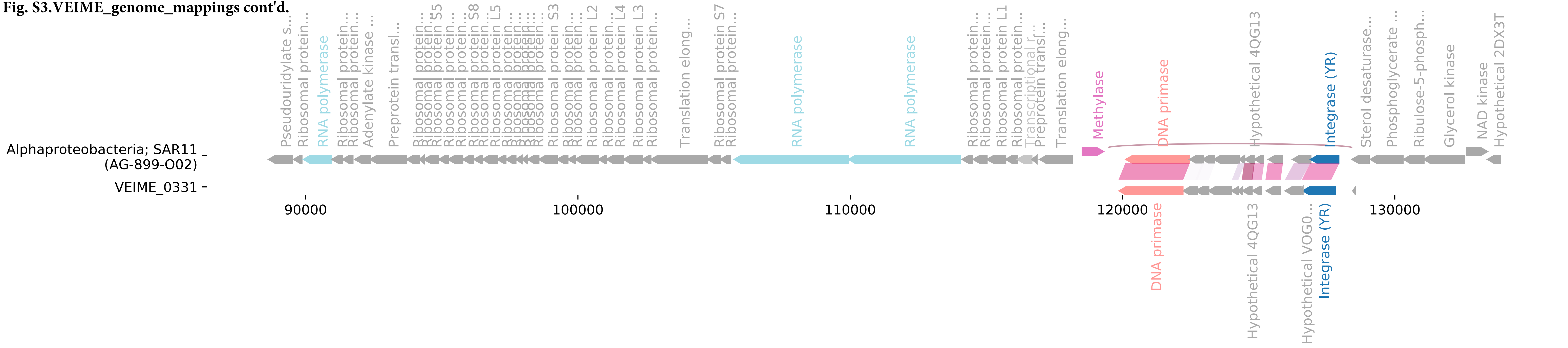

**Fig. S3.VEIME_genome_mappings cont'd.**

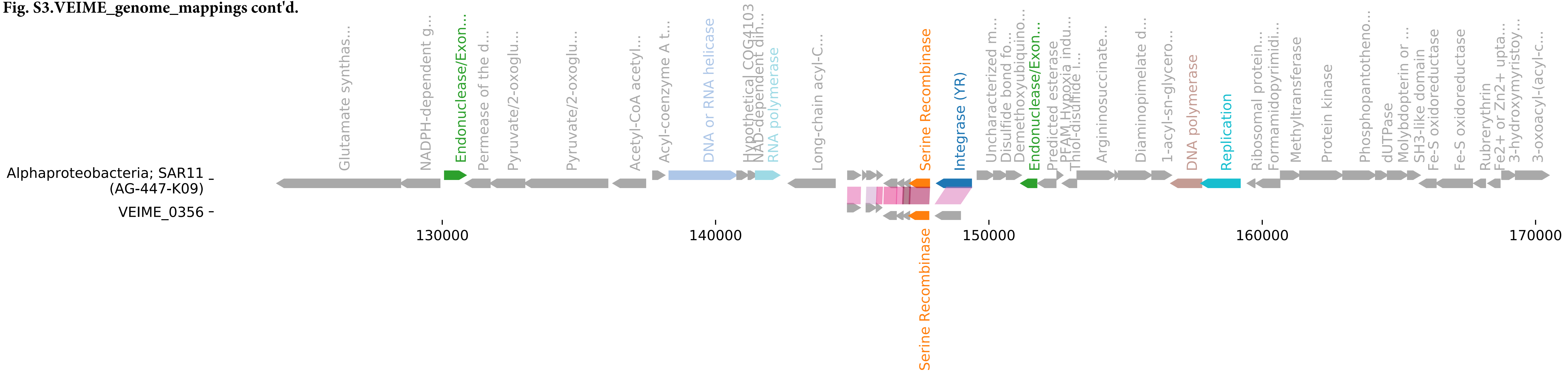

**Fig. S3.VEIME_genome_mappings cont'd.**

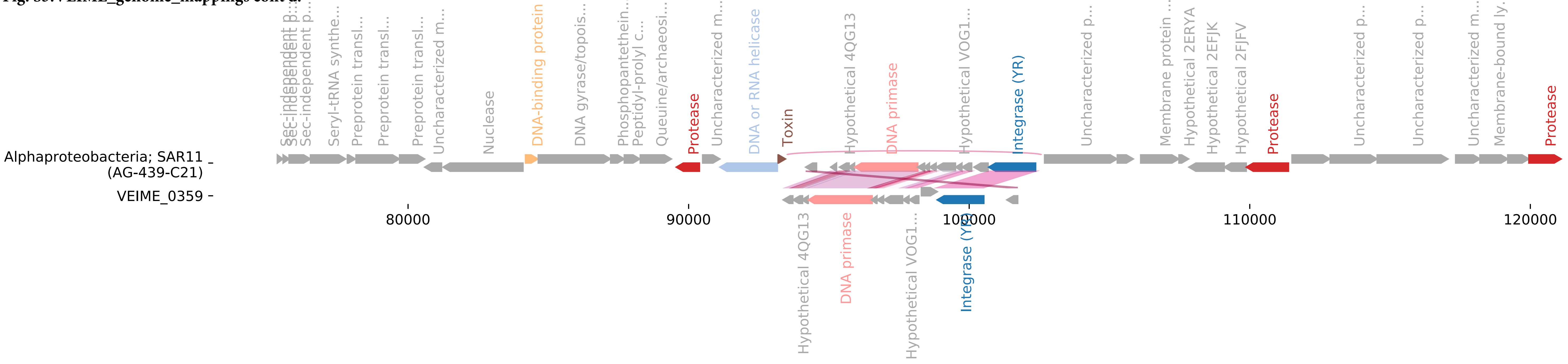

**Fig. S3.VEIME_genome_mappings cont'd.**

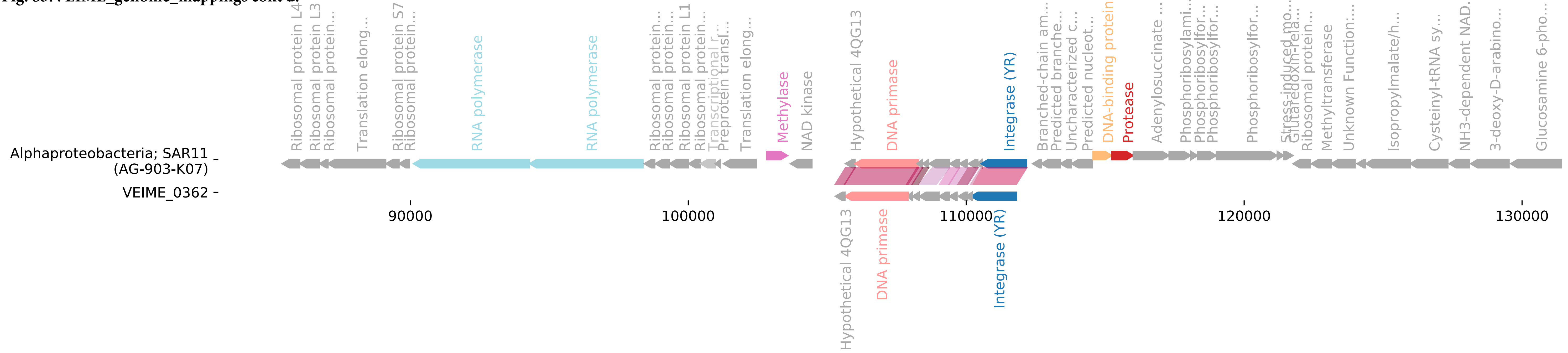

**Fig. S3.VEIME_genome_mappings cont'd.**

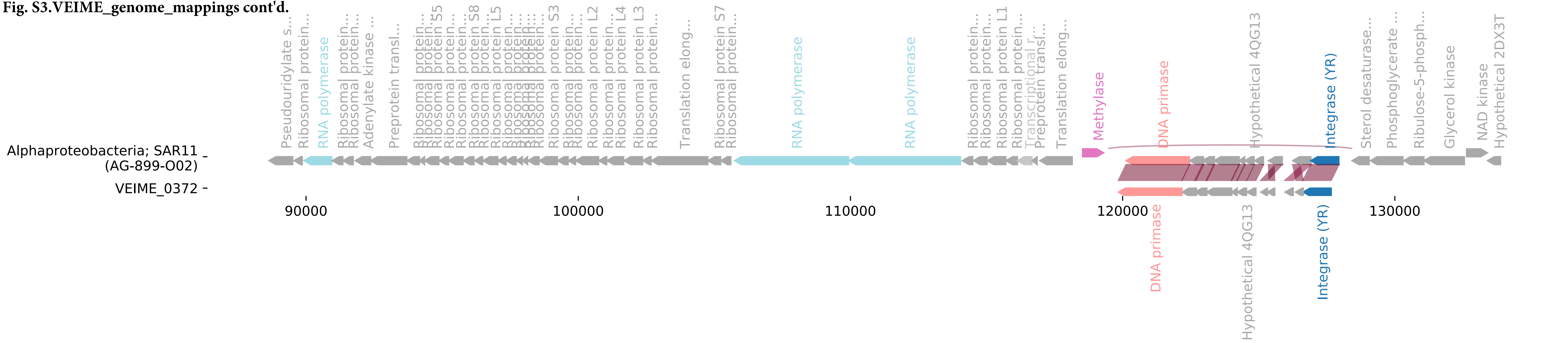

**Fig. S3.VEIME_genome_mappings cont'd.**

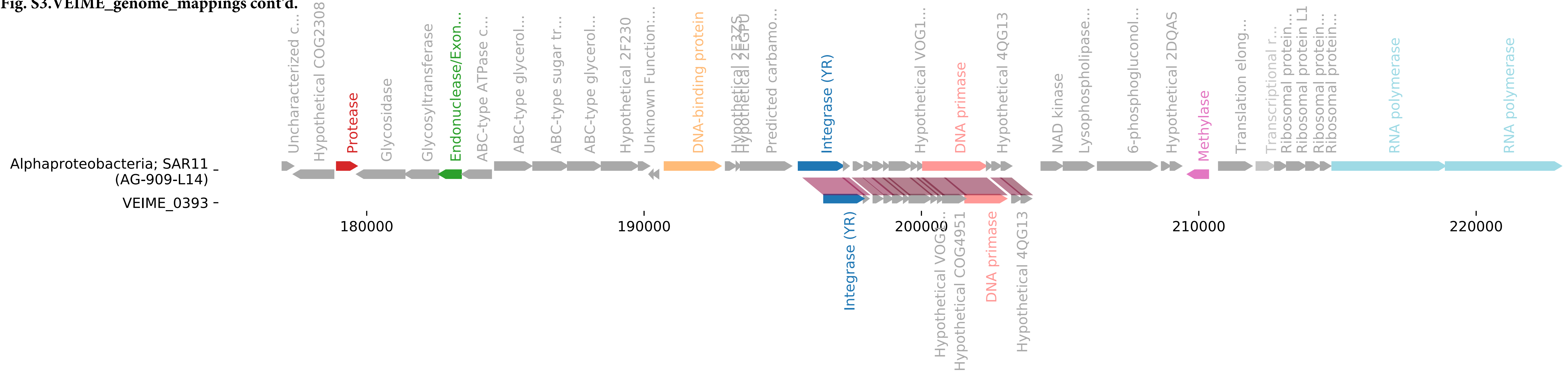

**Fig. S3.VEIME_genome_mappings cont'd.**

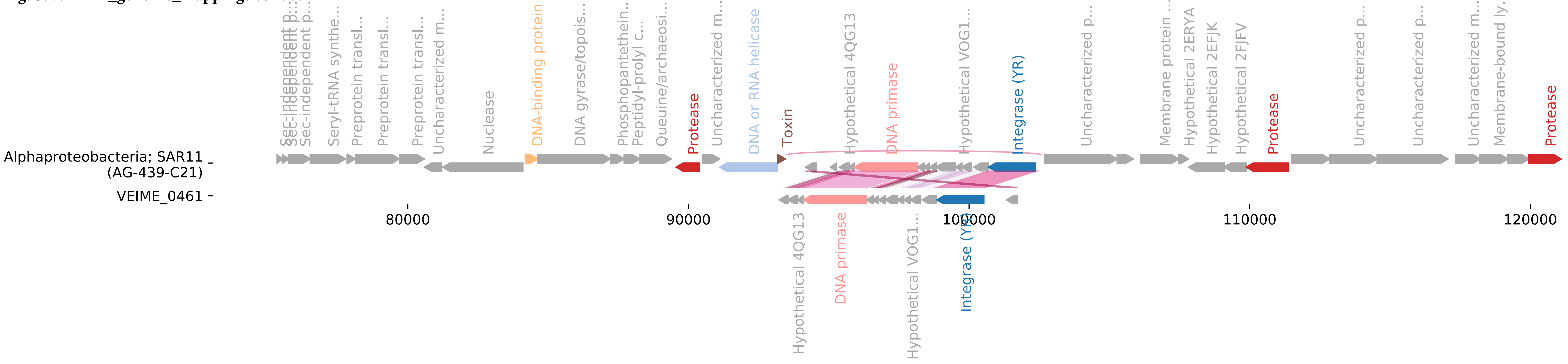

**Fig. S3.VEIME_genome_mappings cont'd.**

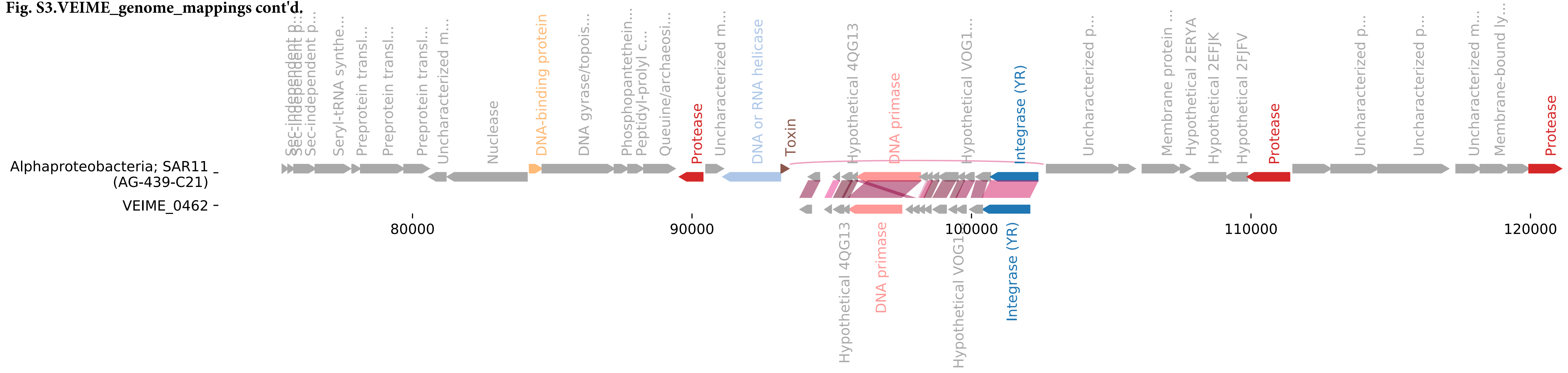

**Fig. S3.VEIME_genome_mappings cont'd.**

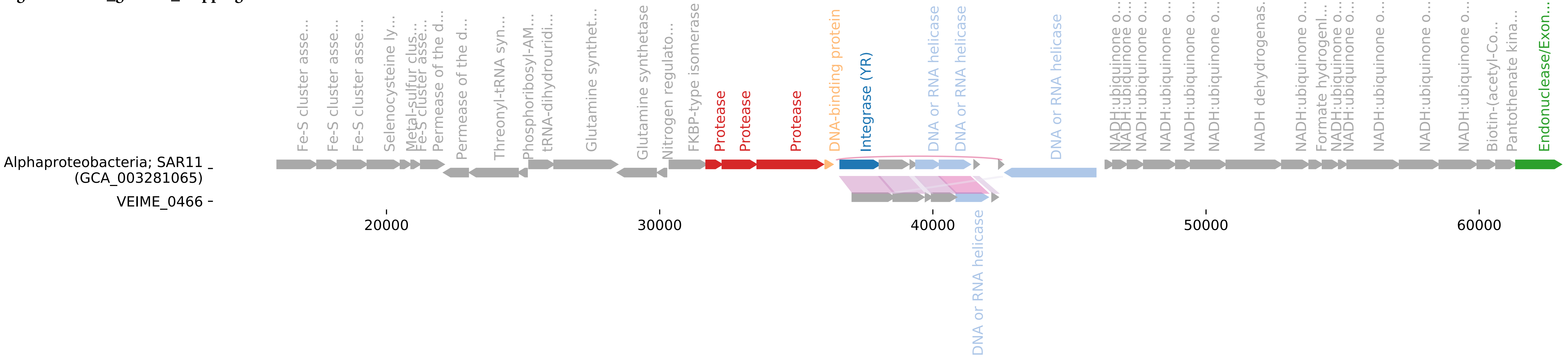

**Fig. S3.VEIME_genome_mappings cont'd.**

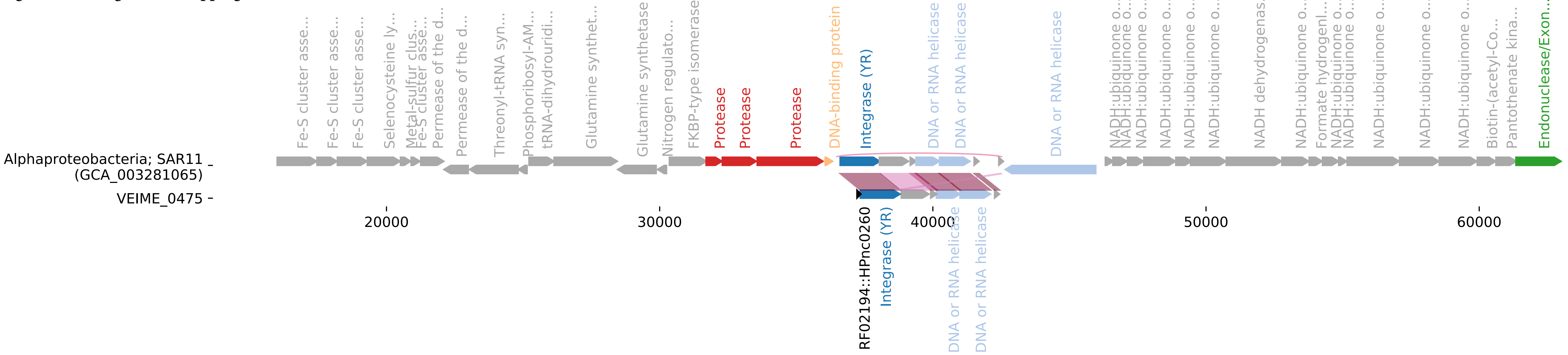

**Fig. S3.VEIME_genome_mappings cont'd.**

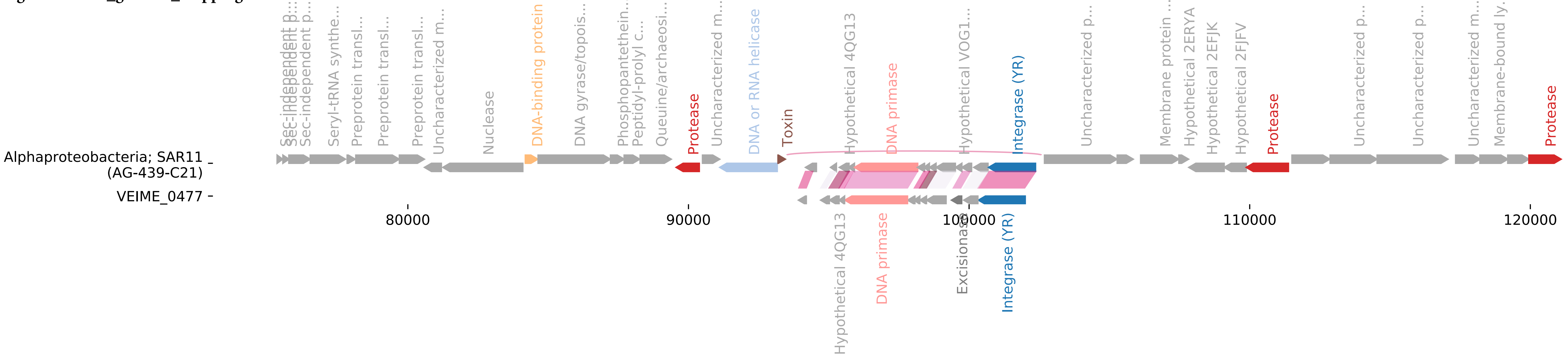

**Fig. S3.VEIME_genome_mappings cont'd.**

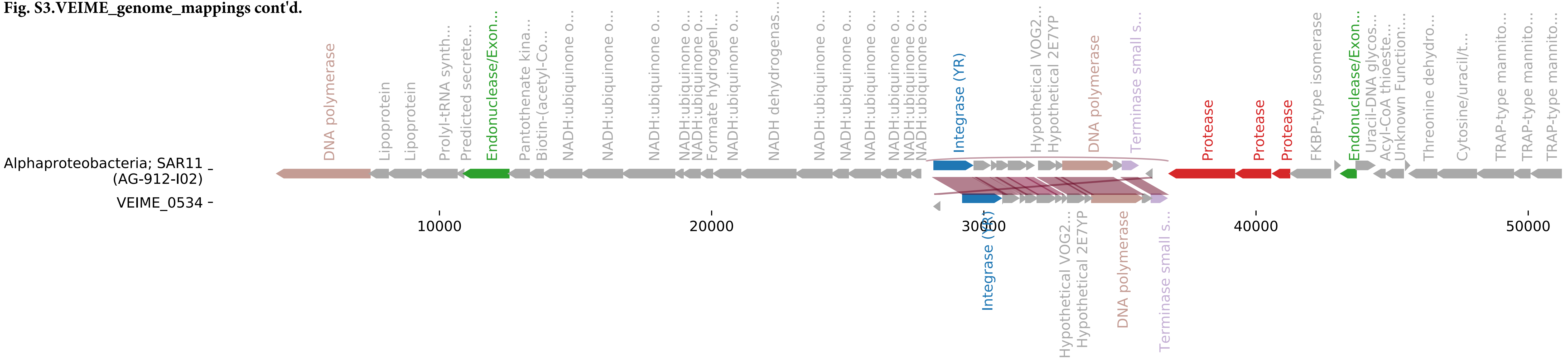

**Fig. S3.VEIME_genome_mappings cont'd.**

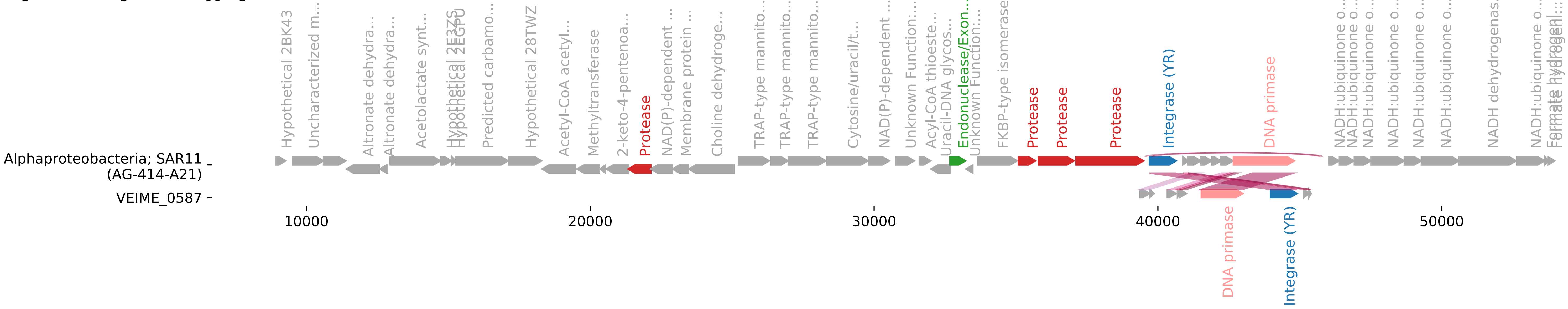

**Fig. S3.VEIME_genome_mappings cont'd.**

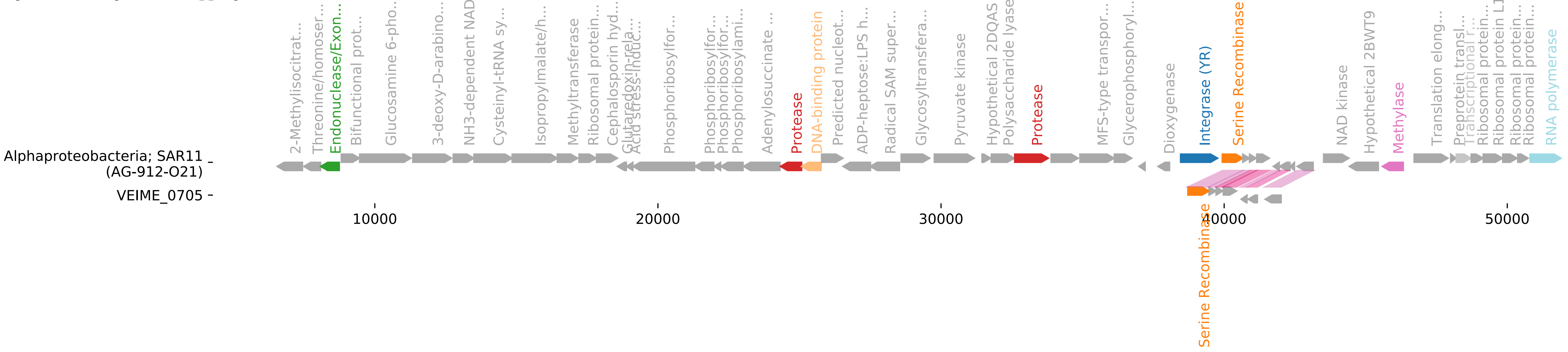

**Fig. S3.VEIME_genome_mappings cont'd.**

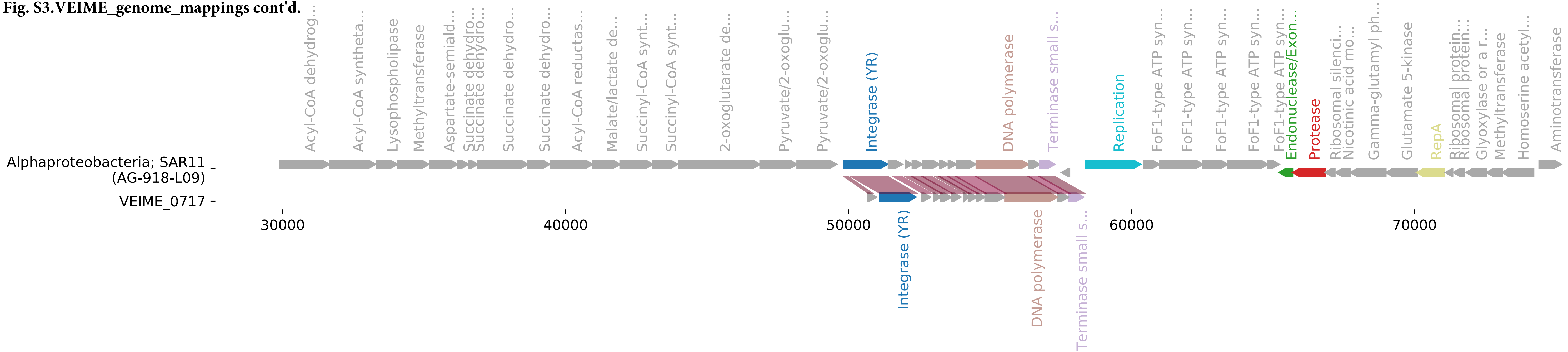

**Fig. S3.VEIME_genome_mappings cont'd.**

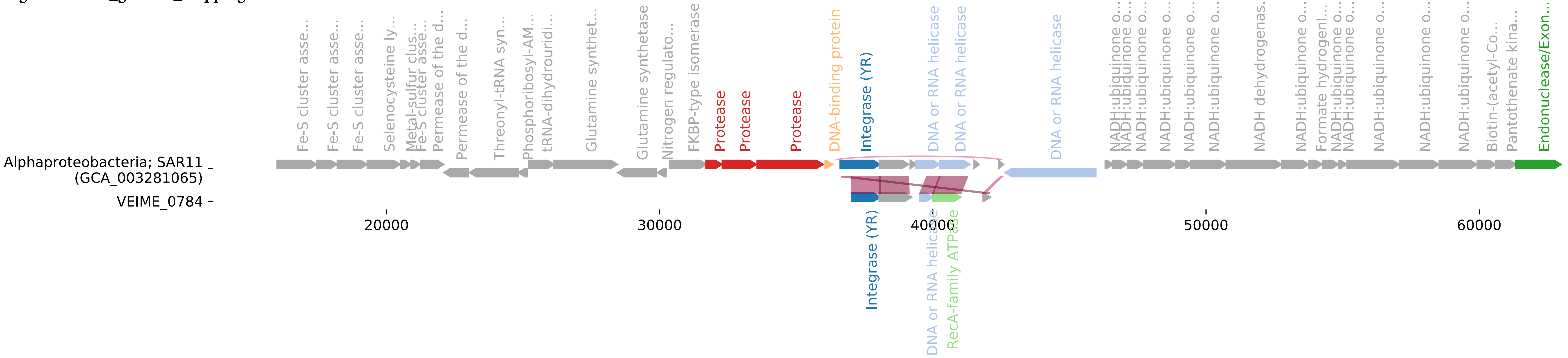

**Fig. S3.VEIME_genome_mappings cont'd.**

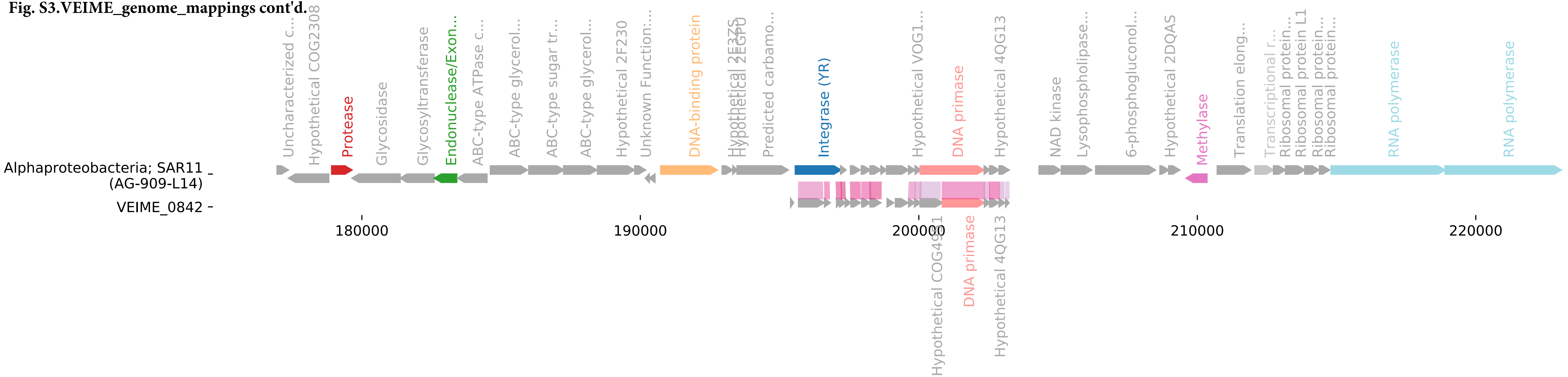

**Fig. S3.VEIME_genome_mappings cont'd.**

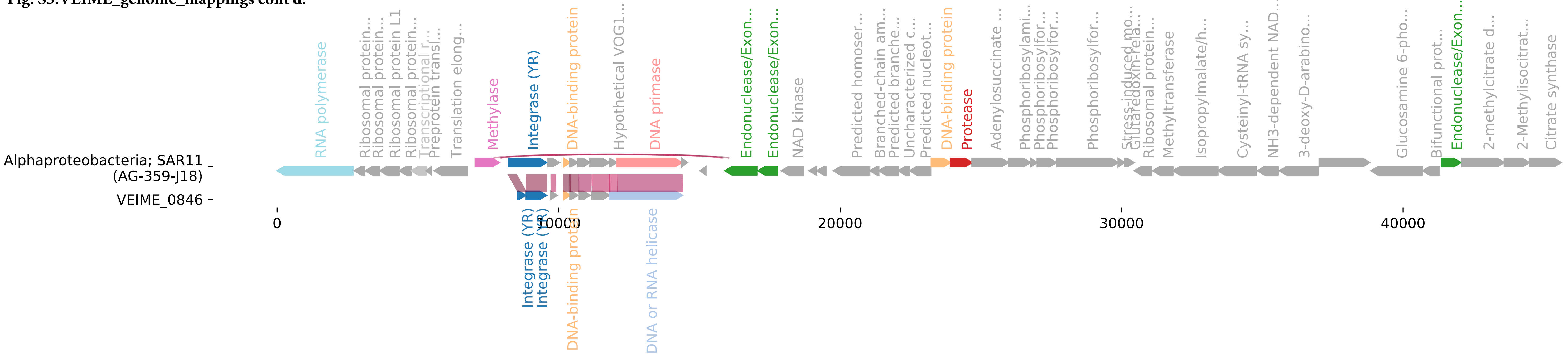

**Fig. S3.VEIME_genome_mappings cont'd.**

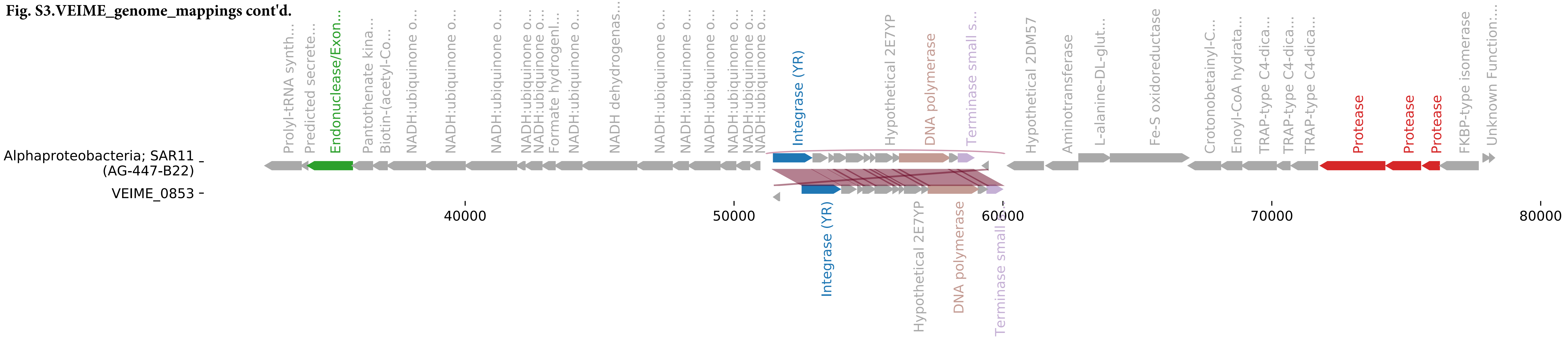

**Fig. S3.VEIME_genome_mappings cont'd.**

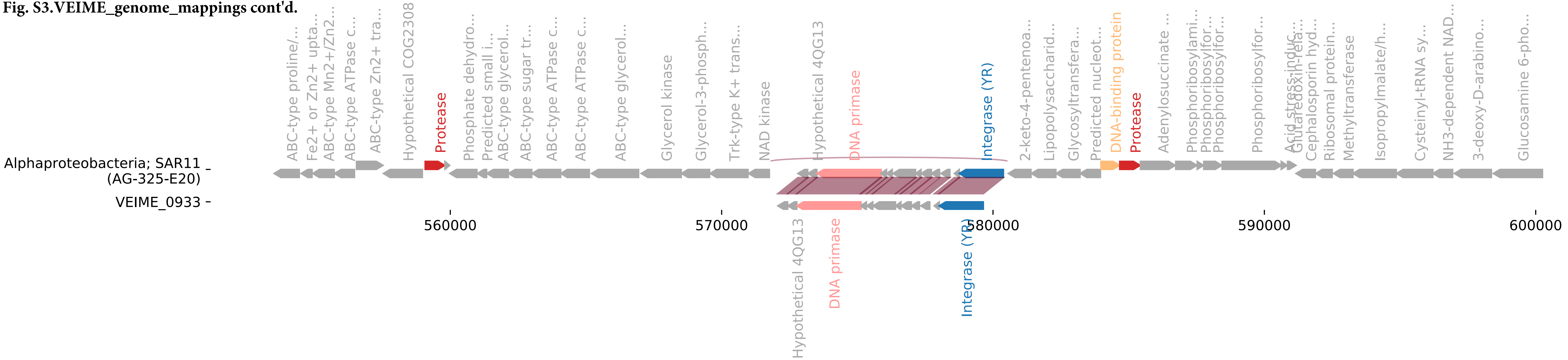

**Fig. S3.VEIME_genome_mappings cont'd.**

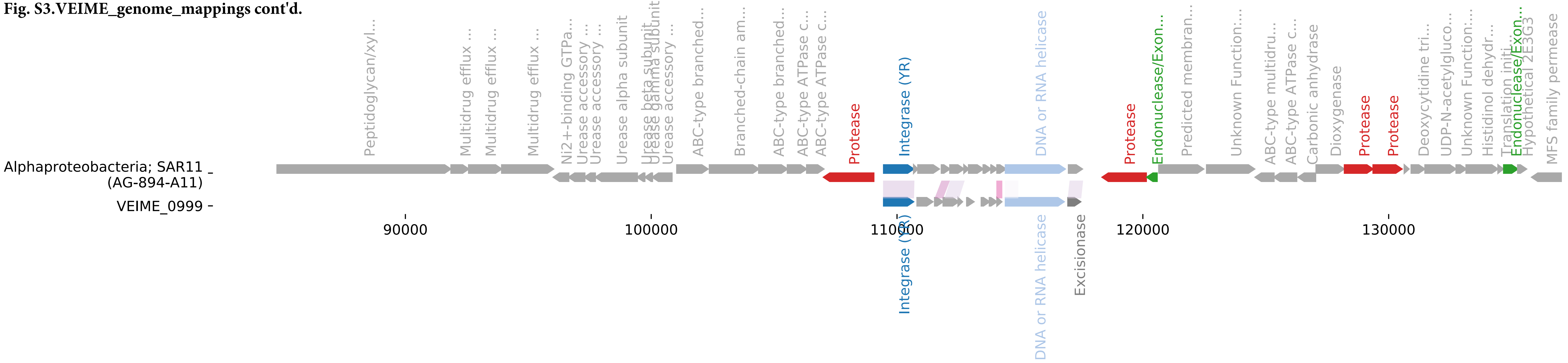

**Fig. S3.VEIME_genome_mappings cont'd.**

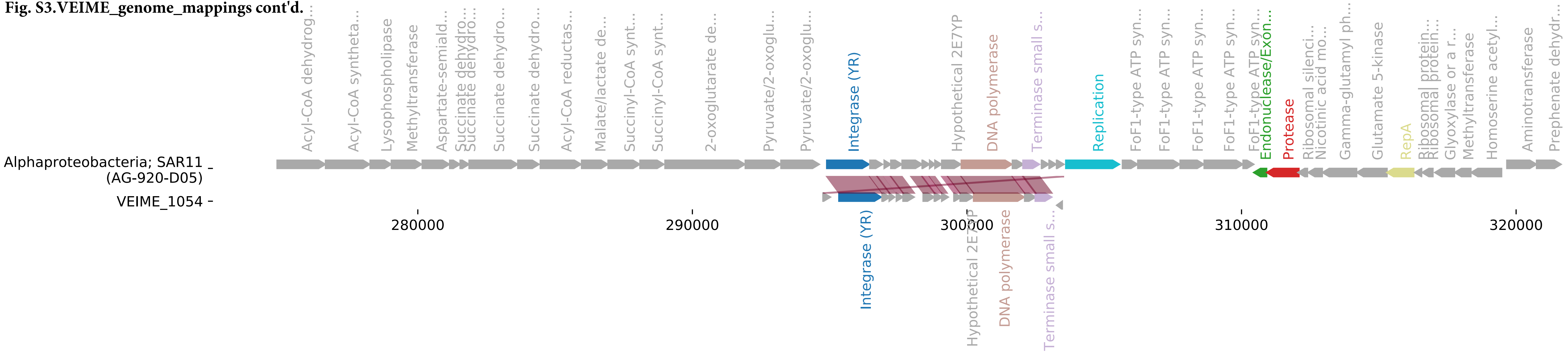

**Fig. S3.VEIME_genome_mappings cont'd.**

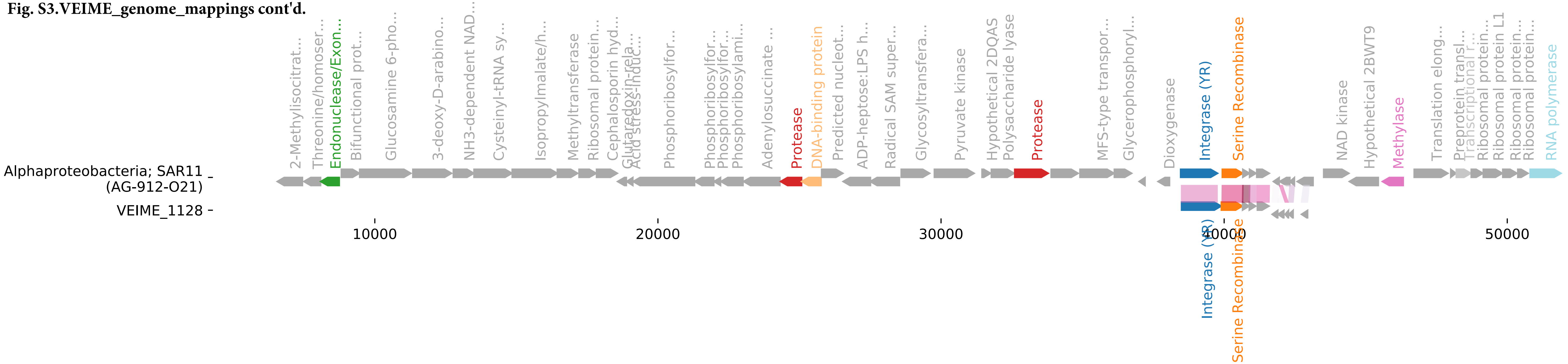

**Fig. S3.VEIME_genome_mappings cont'd.**

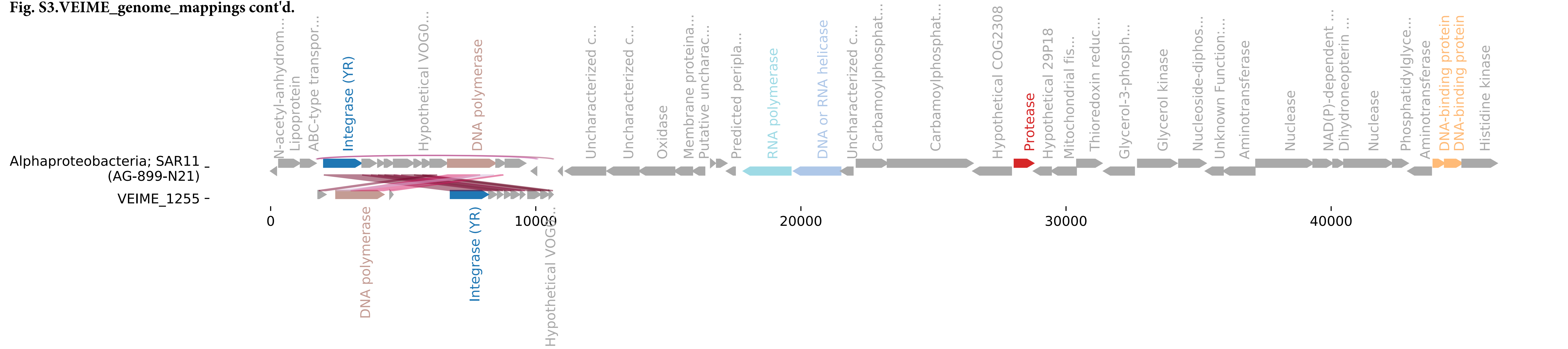

**Fig. S3.VEIME_genome_mappings cont'd.**

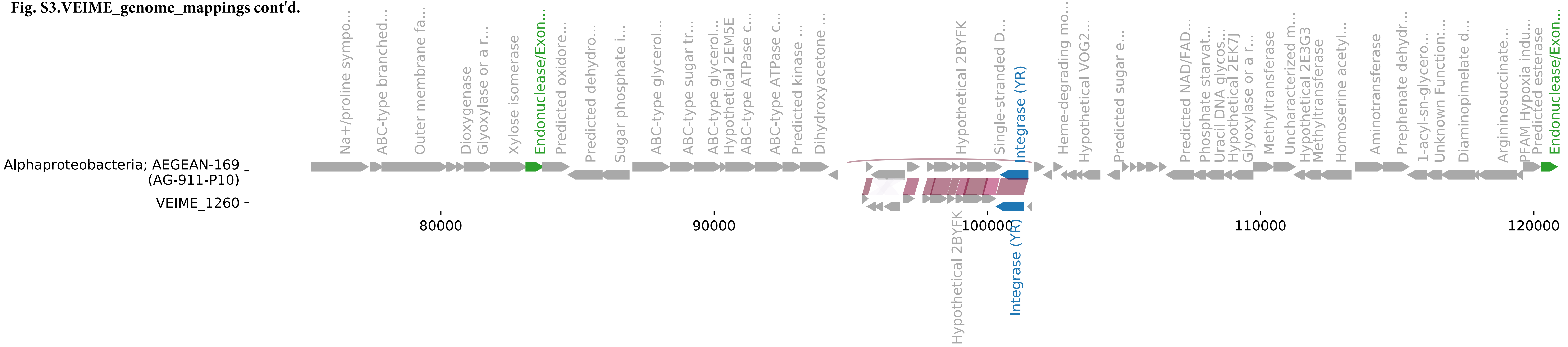

**Fig. S3.VEIME_genome_mappings cont'd.**

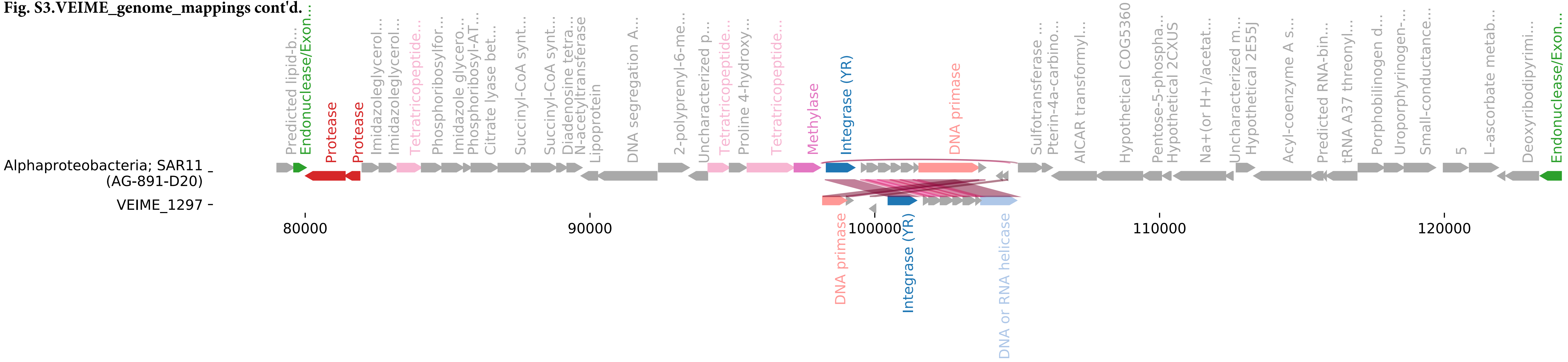

**Fig. S3.VEIME_genome_mappings cont'd.**

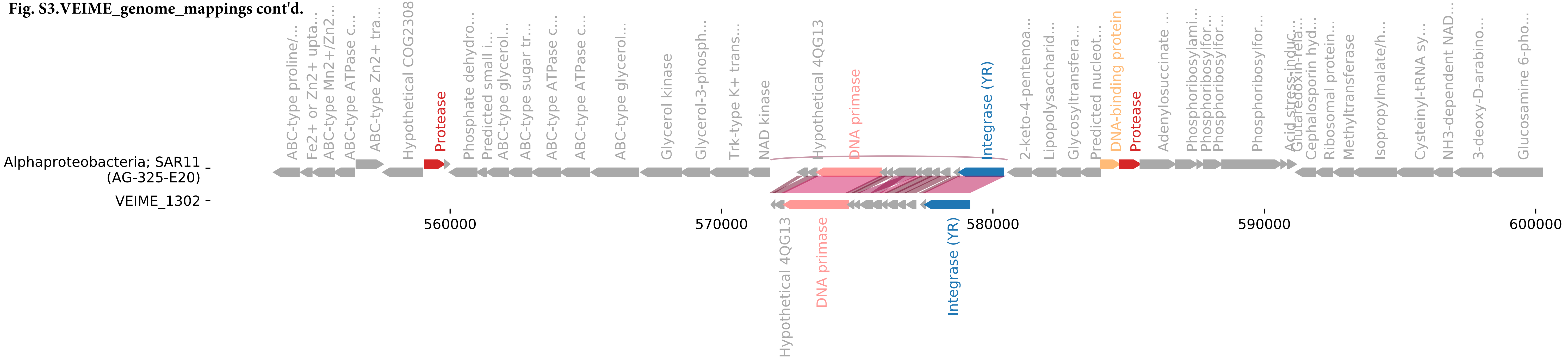

**Fig. S3.VEIME_genome_mappings cont'd.**

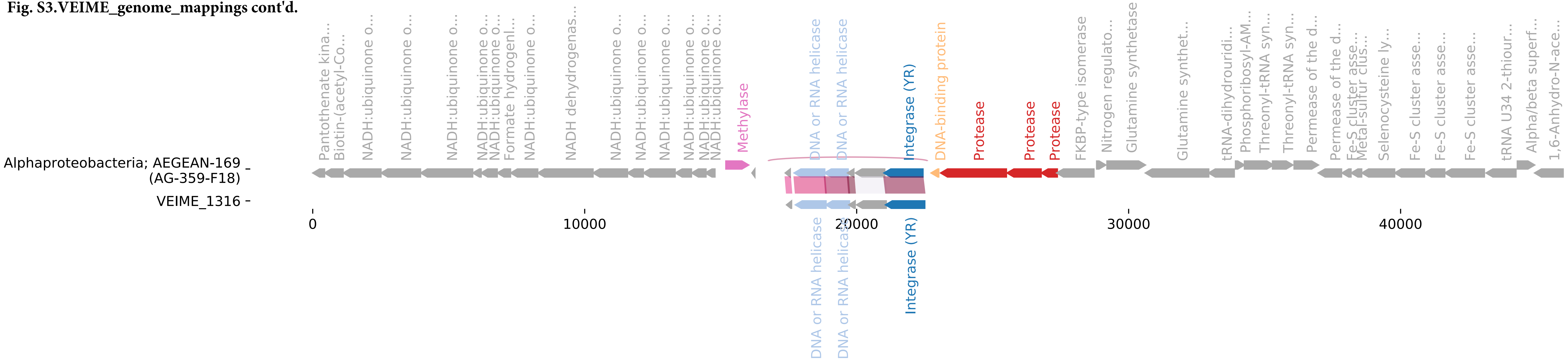

**Fig. S3.VEIME_genome_mappings cont'd.**

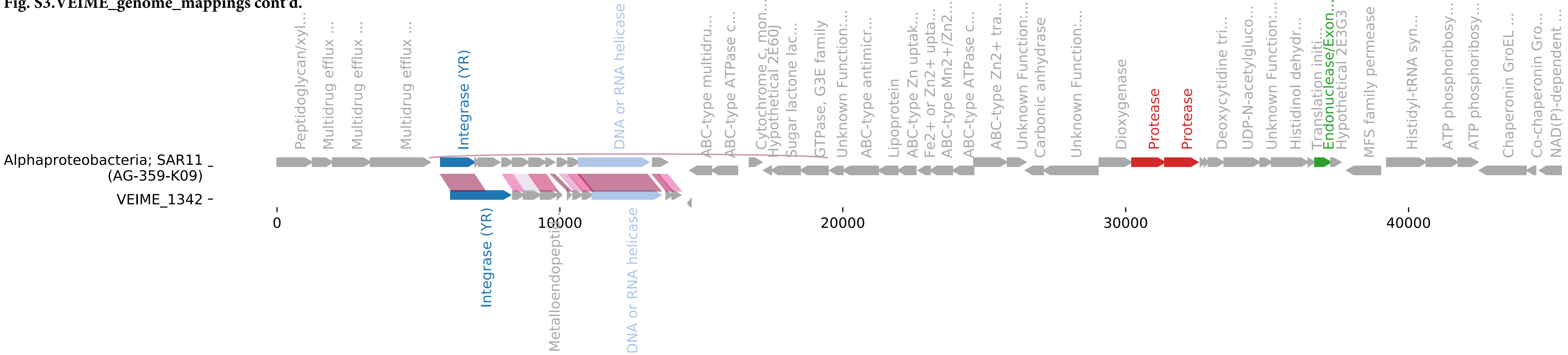

**Fig. S3.VEIME_genome_mappings cont'd.**

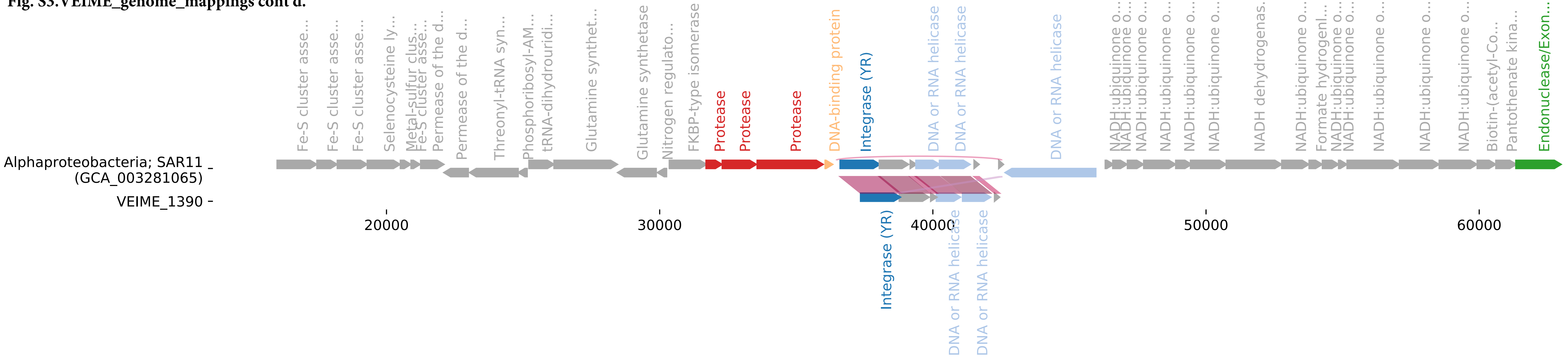

Fig. S3.VEIME_genome_mappings cont'd.

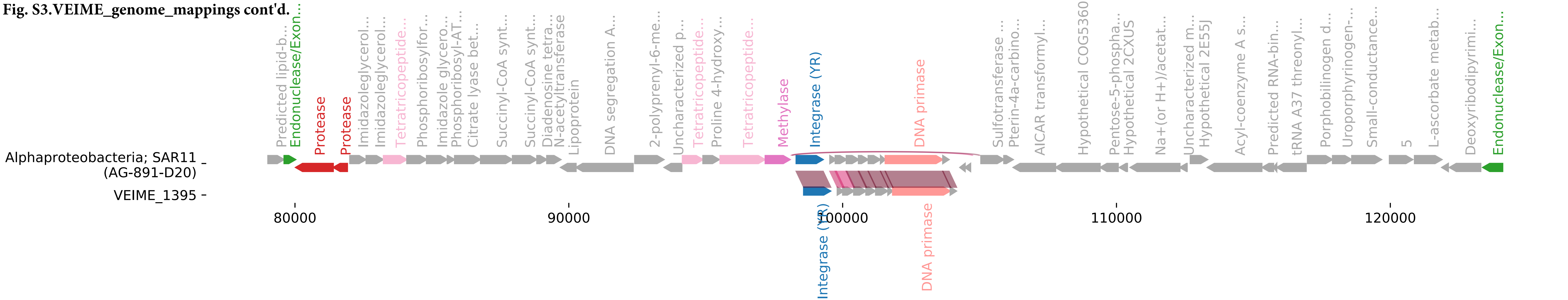

**Fig. S3.VEIME_genome_mappings cont'd.**

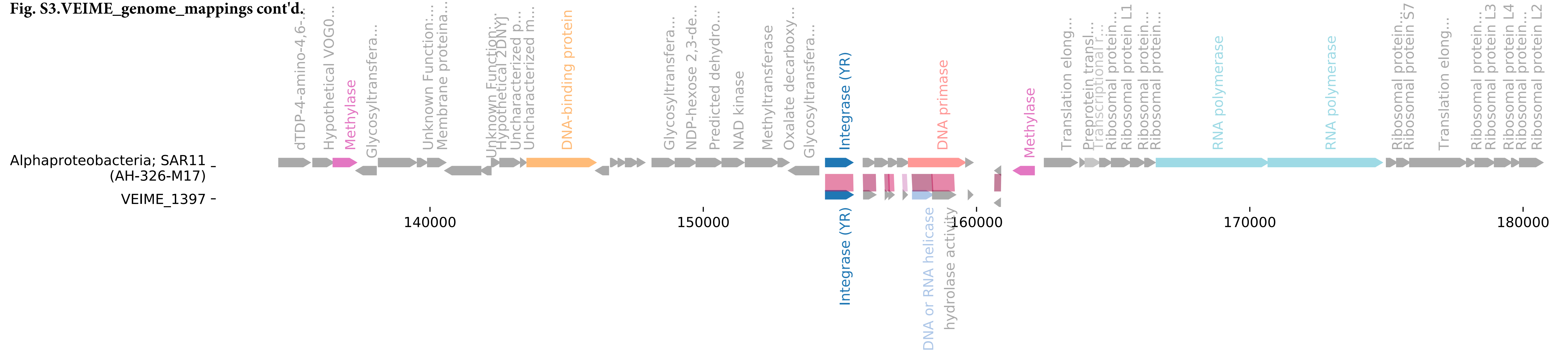

**Fig. S3.VEIME_genome_mappings cont'd.**

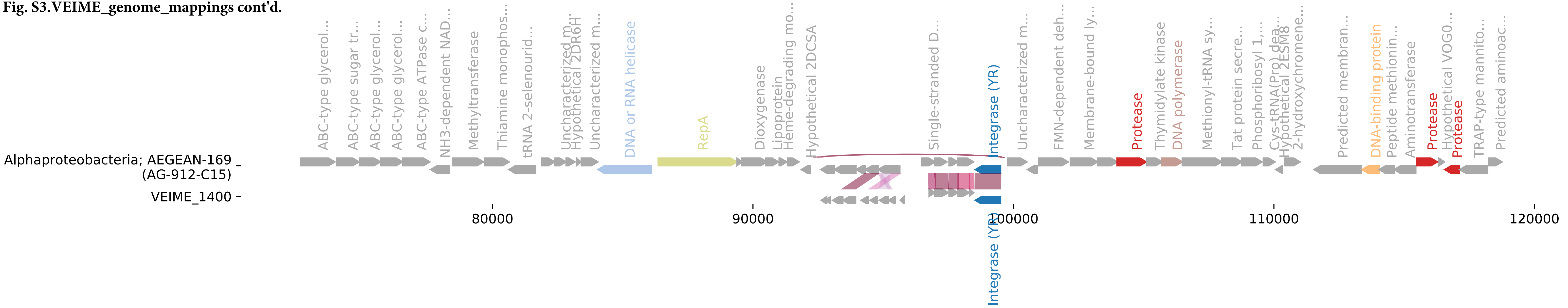

**Fig. S3.VEIME_genome_mappings cont'd.**

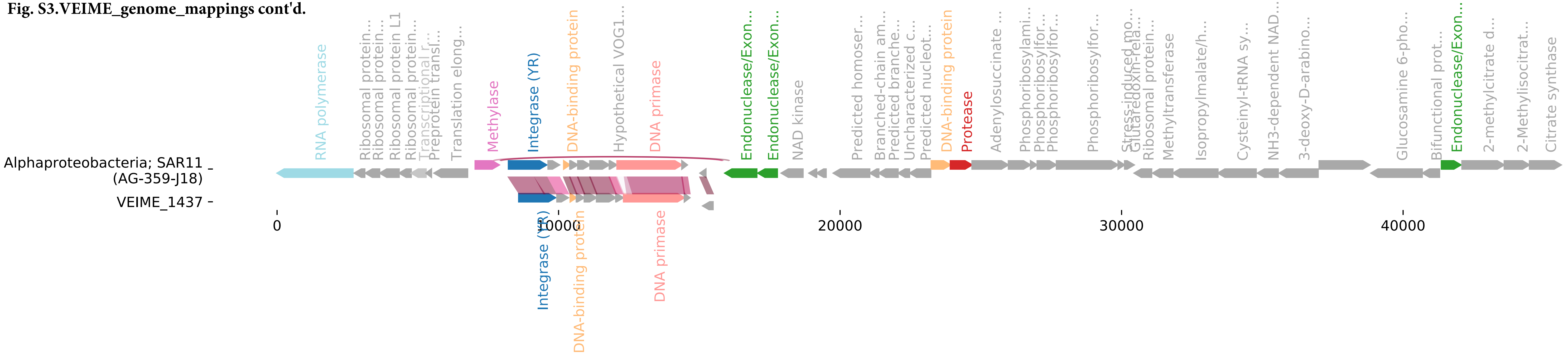

**Fig. S3.VEIME_genome_mappings cont'd.**

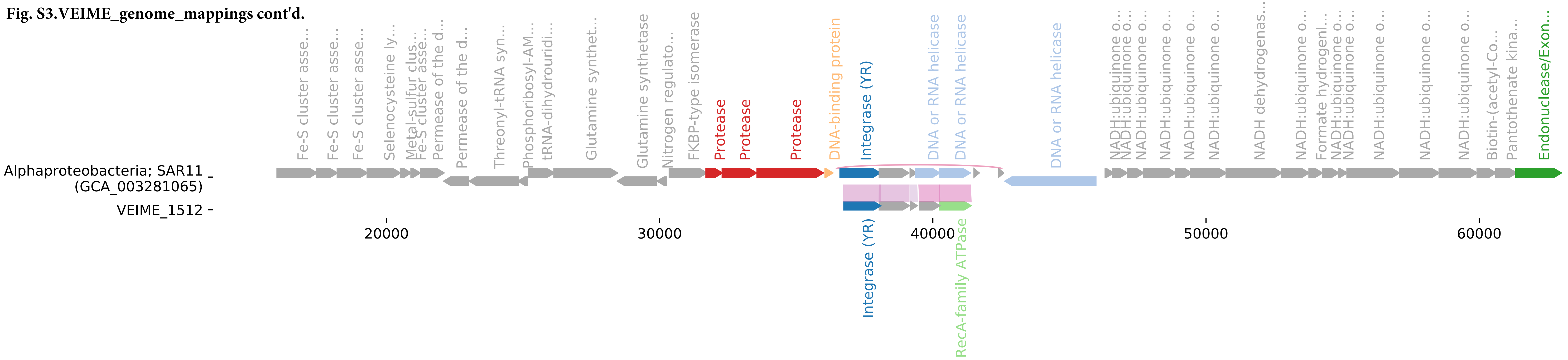

**Fig. S3.VEIME_genome_mappings cont'd.**

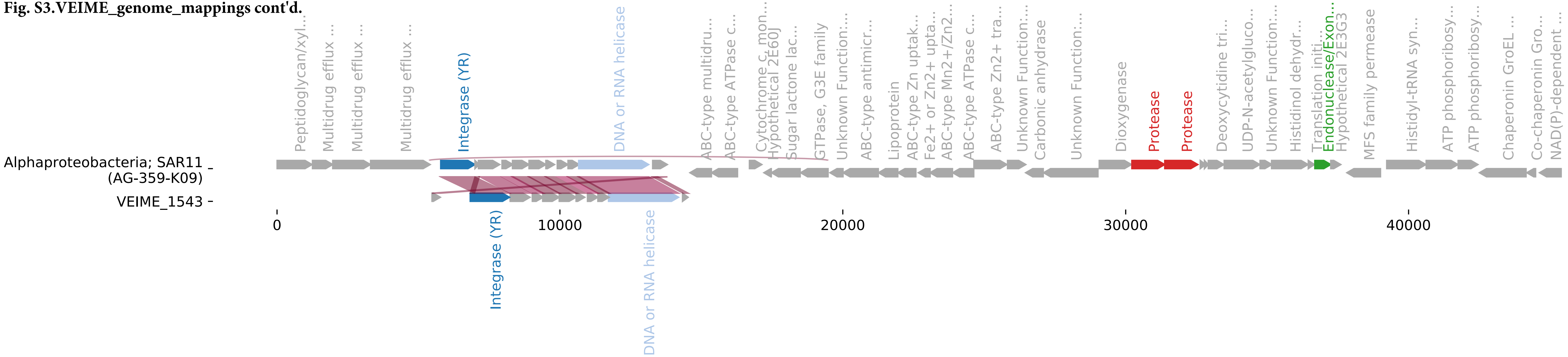

**Fig. S3.VEIME_genome_mappings cont'd.**

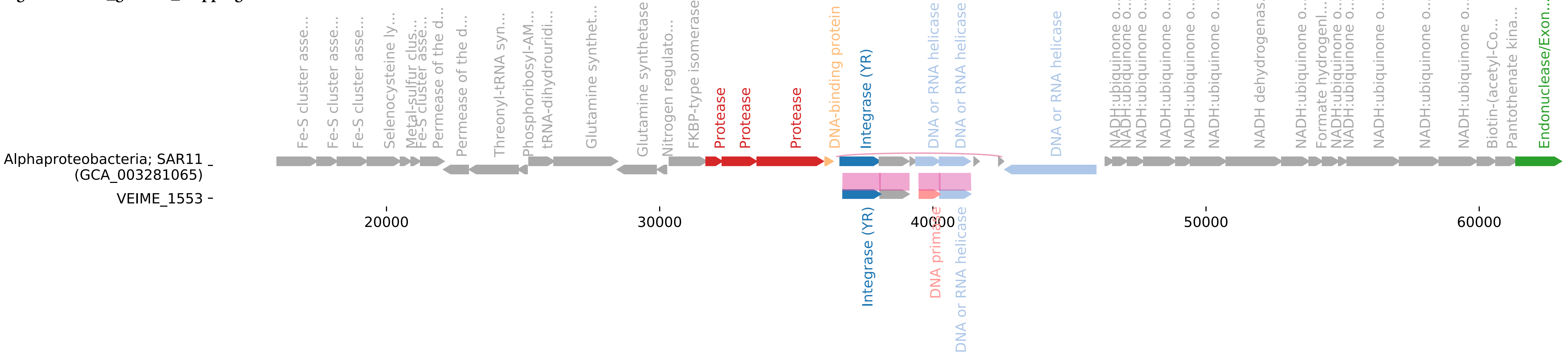

Fig. S3.VEIME_genome_mappings cont'd.

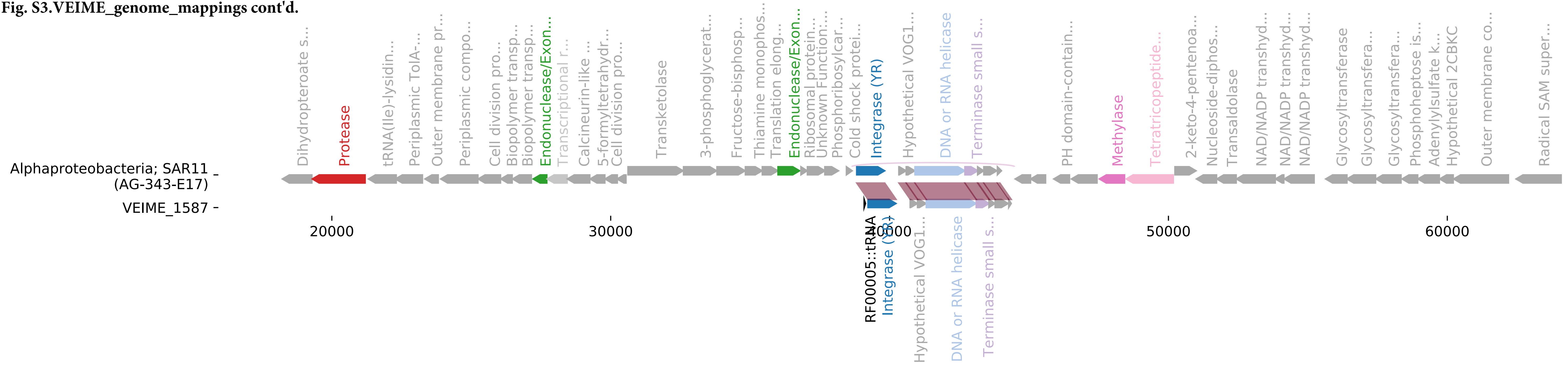

**Fig. S3.VEIME_genome_mappings cont'd.**

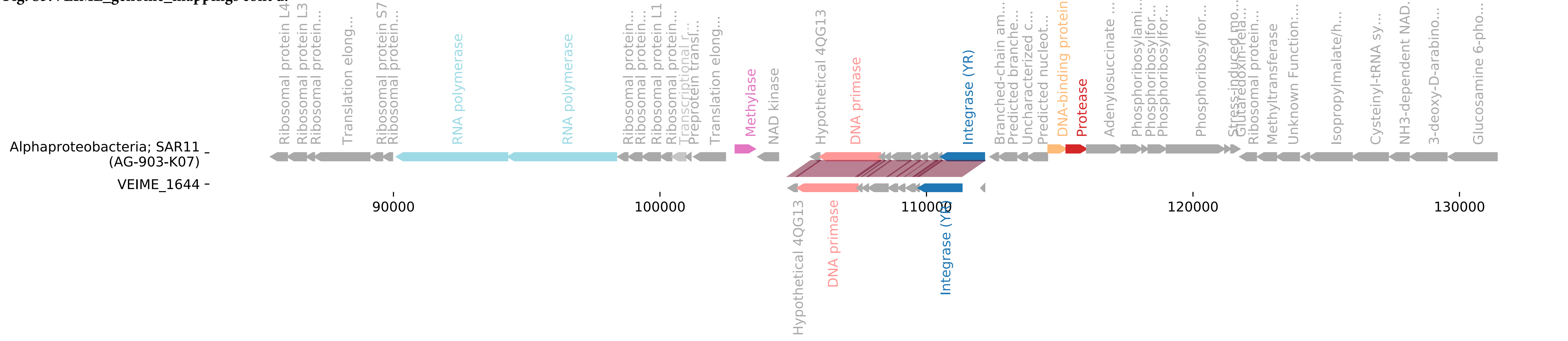

**Fig. S3.VEIME_genome_mappings cont'd.**

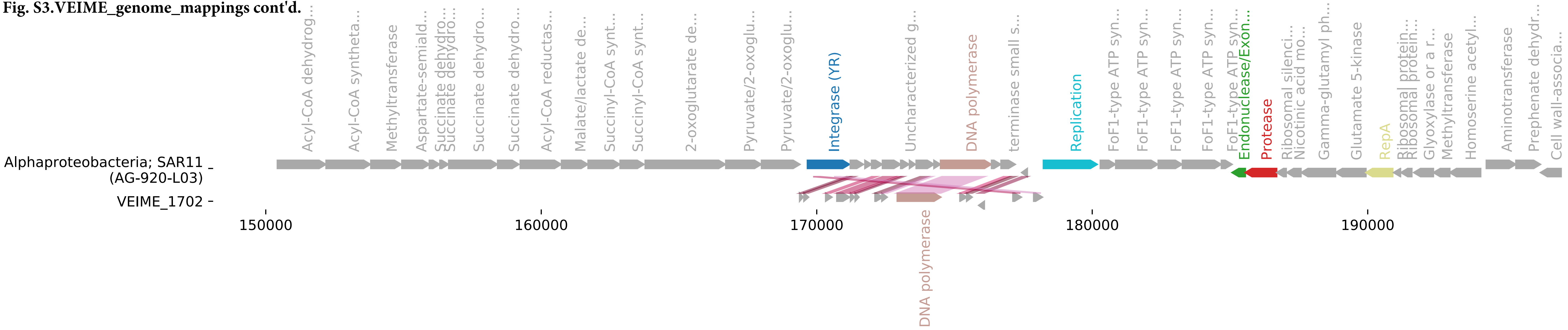

Fig. S3.VEIME_genome_mappings cont'd.

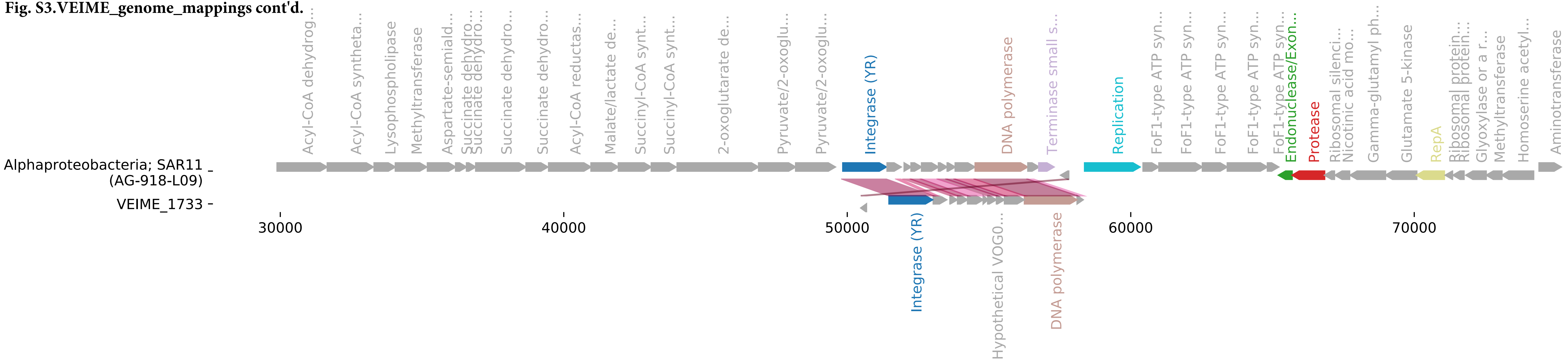

Fig. S3.VEIME_genome_mappings cont'd.

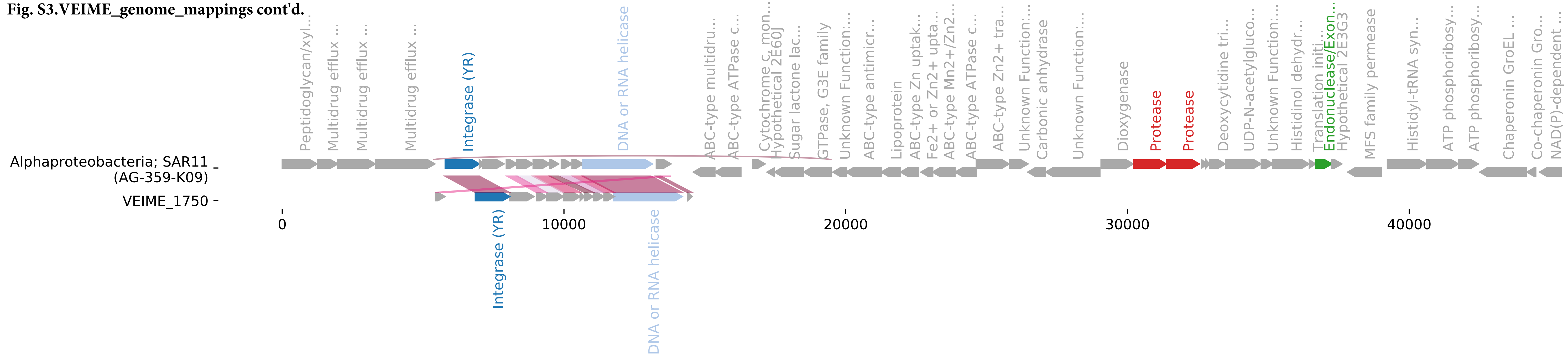

**Fig. S3.VEIME_genome_mappings cont'd.**

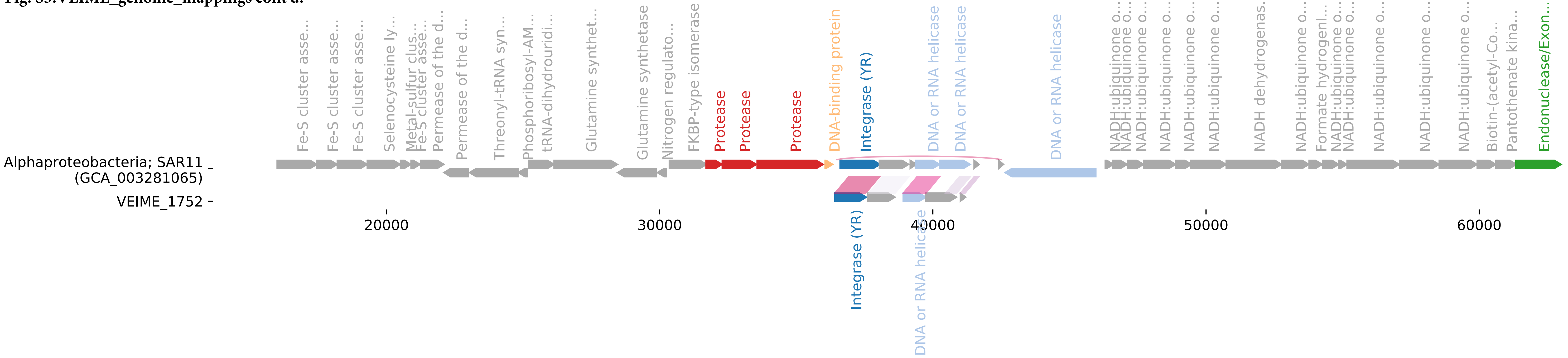

**Fig. S3.VEIME_genome_mappings cont'd.**

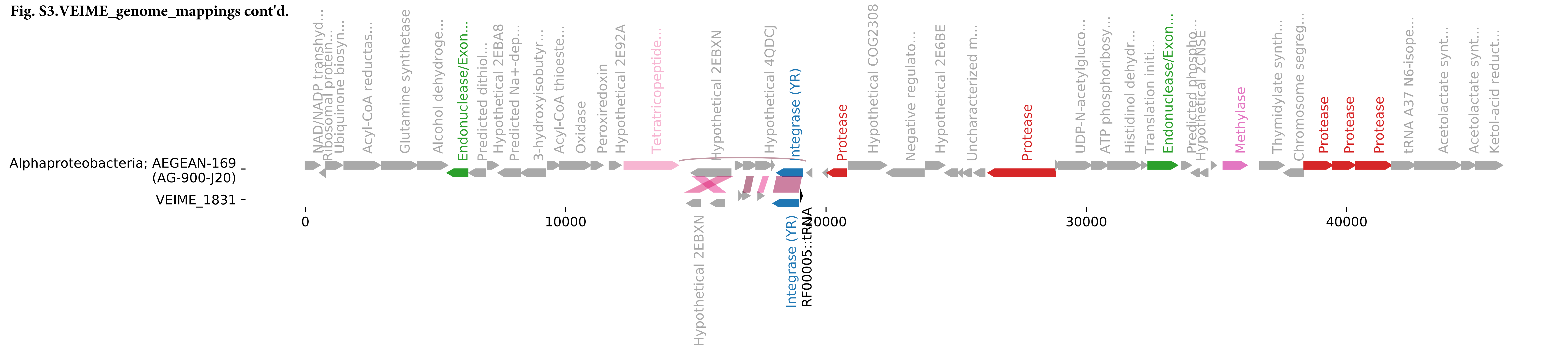

**Fig. S3.VEIME_genome_mappings cont'd.**

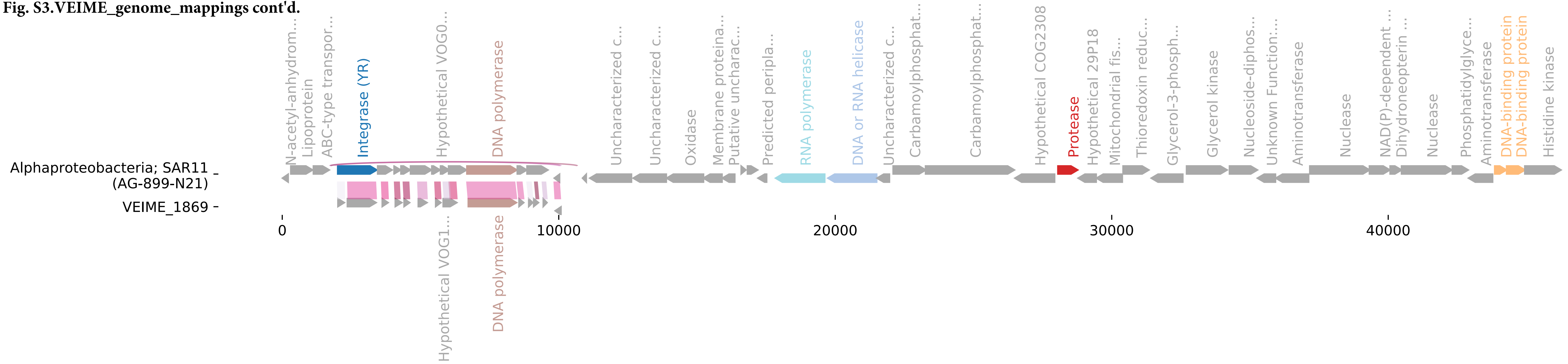

**Fig. S3.VEIME_genome_mappings cont'd.**

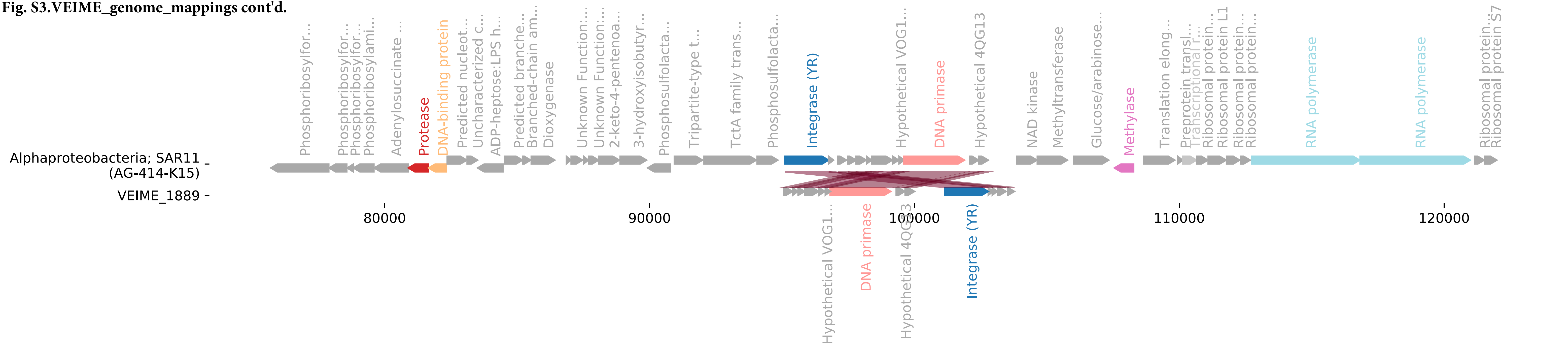

**Fig. S3.VEIME_genome_mappings cont'd.**

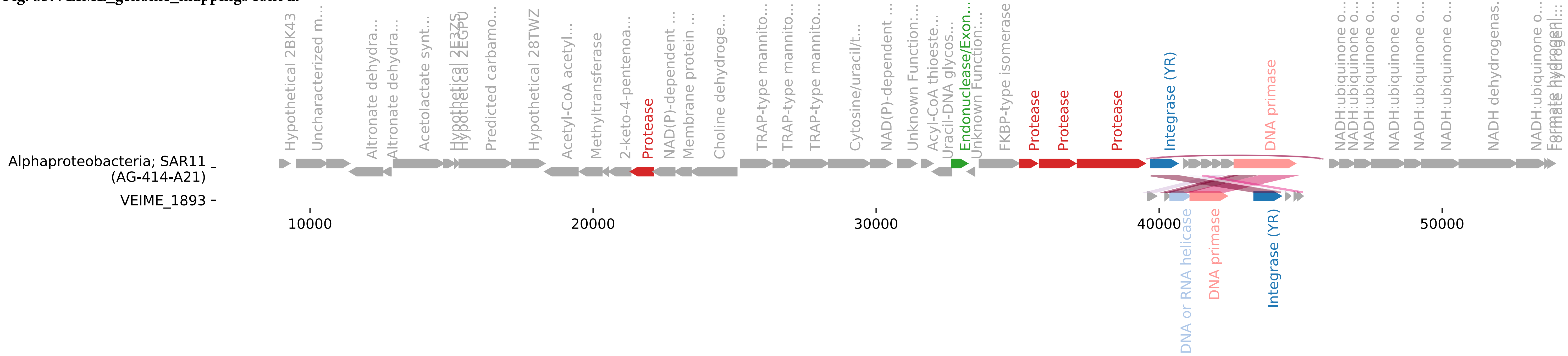

Fig. S3.VEIME_genome_mappings cont'd.

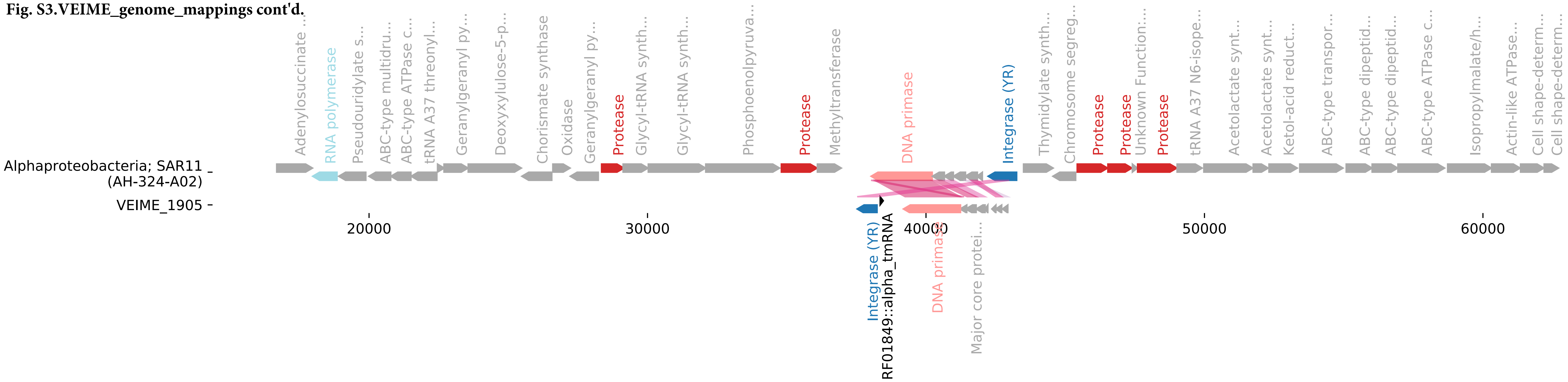

**Fig. S3.VEIME_genome_mappings cont'd.**

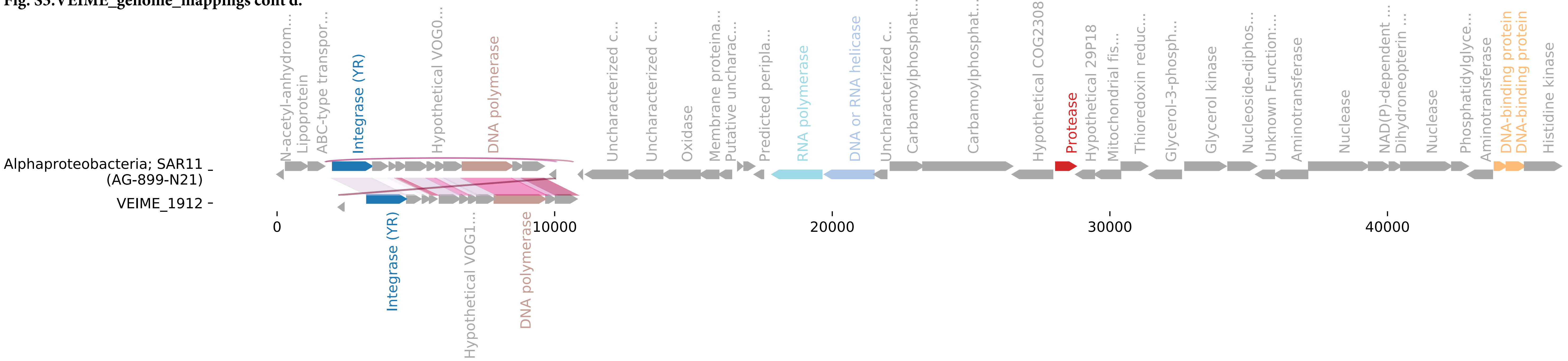

**Fig. S3.VEIME_genome_mappings cont'd.**

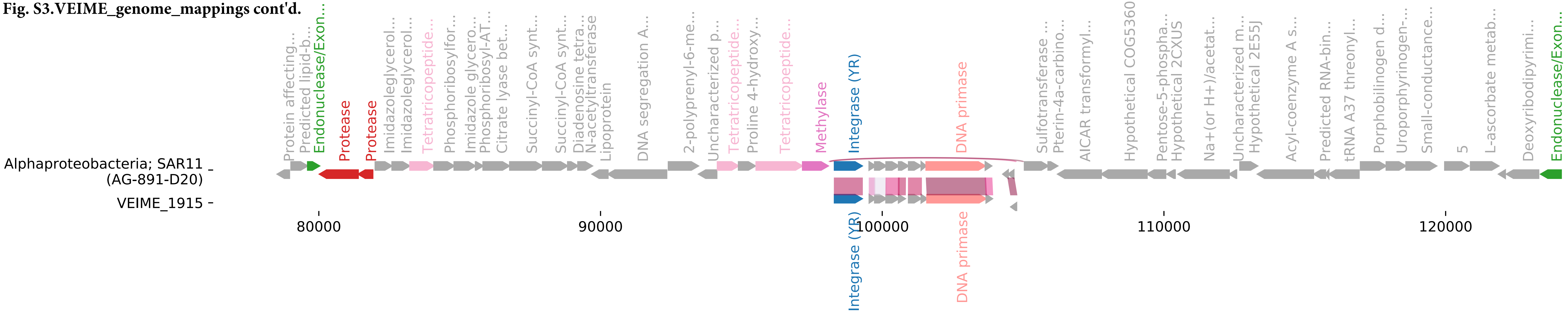

**Fig. S3.VEIME_genome_mappings cont'd.**

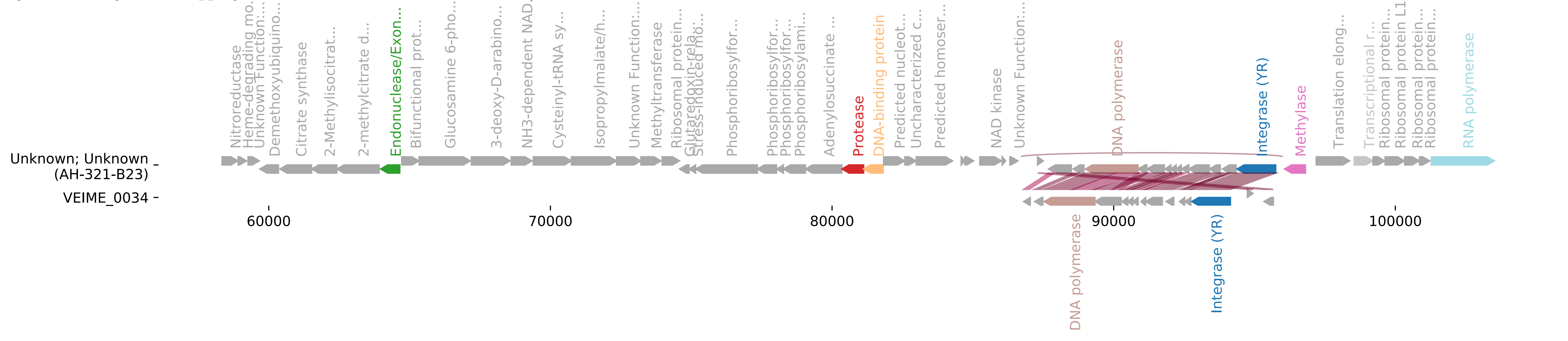

**Fig. S3.VEIME_genome_mappings cont'd.**

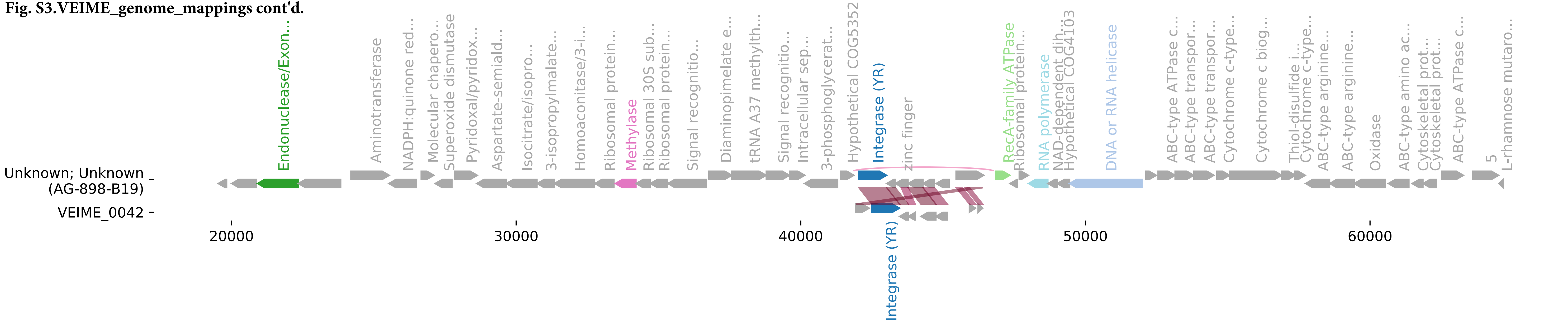

**Fig. S3.VEIME_genome_mappings cont'd.**

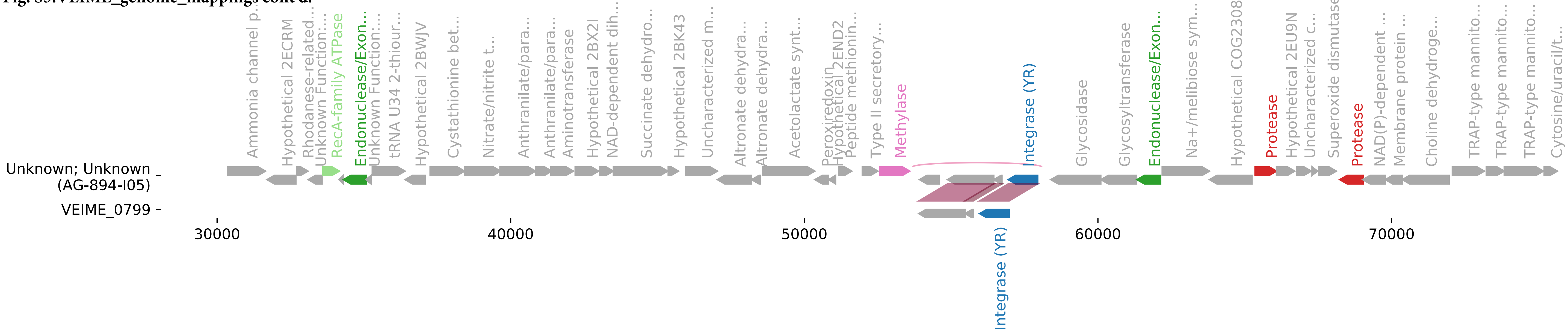

**Fig. S3.VEIME_genome_mappings cont'd.**

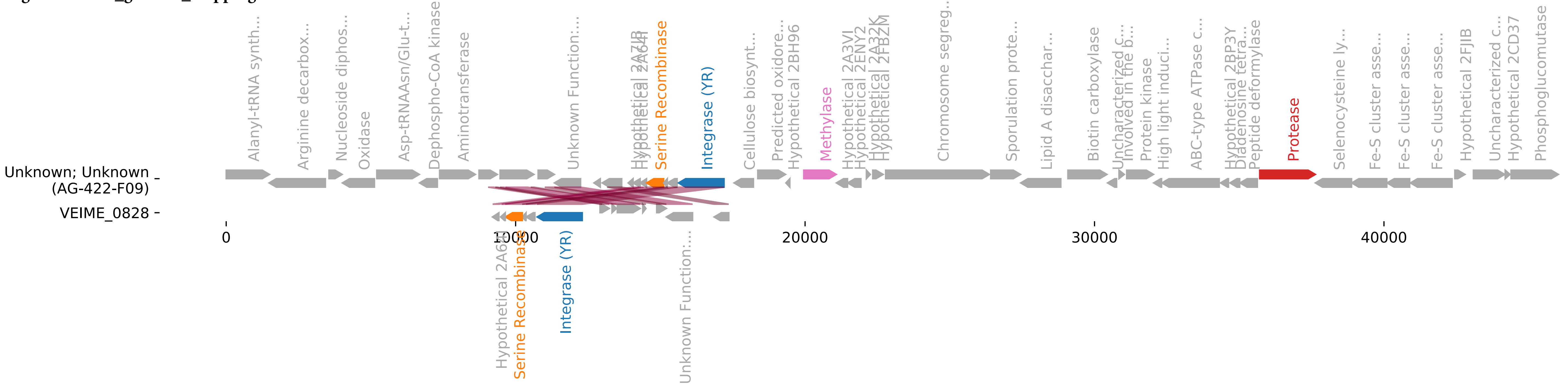

Fig. S3.VEIME_genome_mappings cont'd.

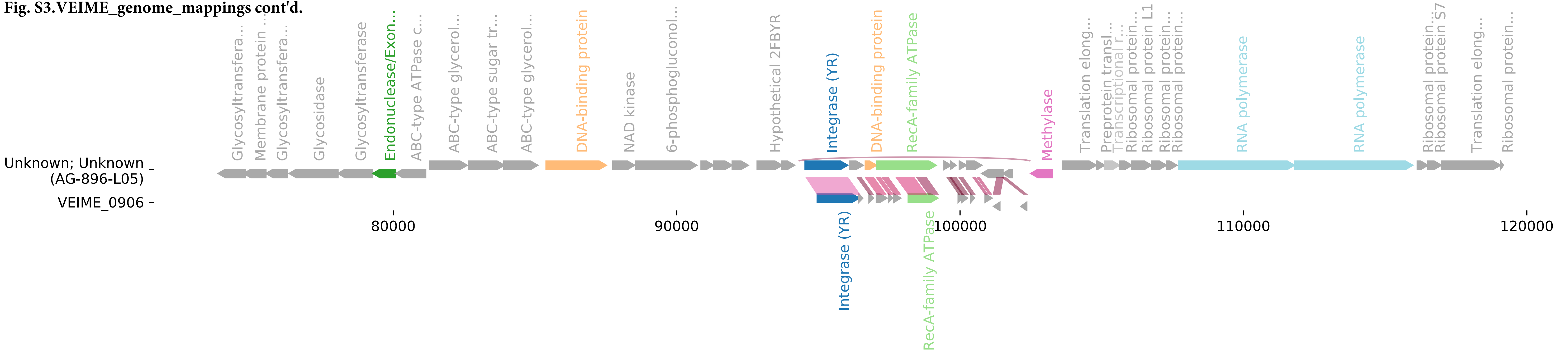

Fig. S3.VEIME_genome_mappings cont'd.

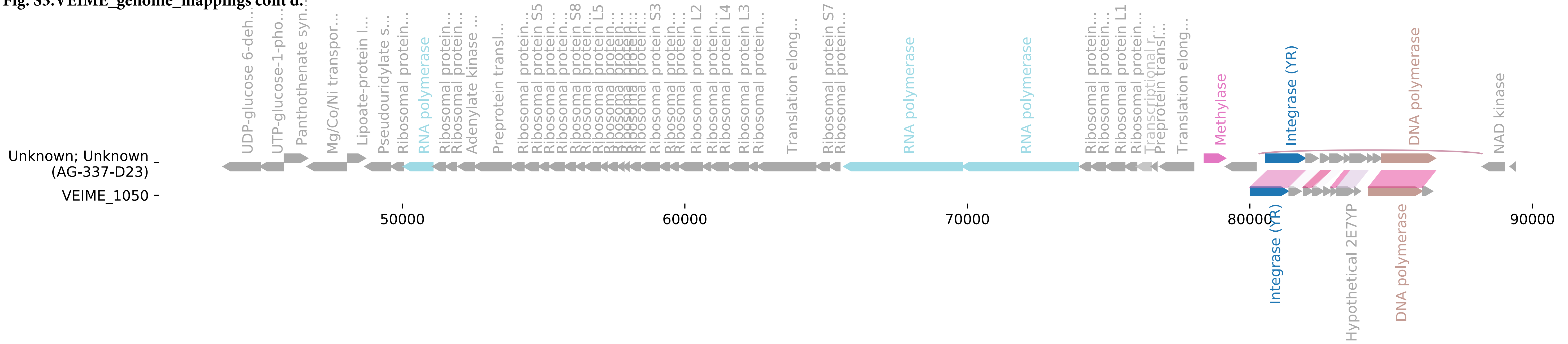

Fig. S3.VEIME_genome_mappings cont'd.

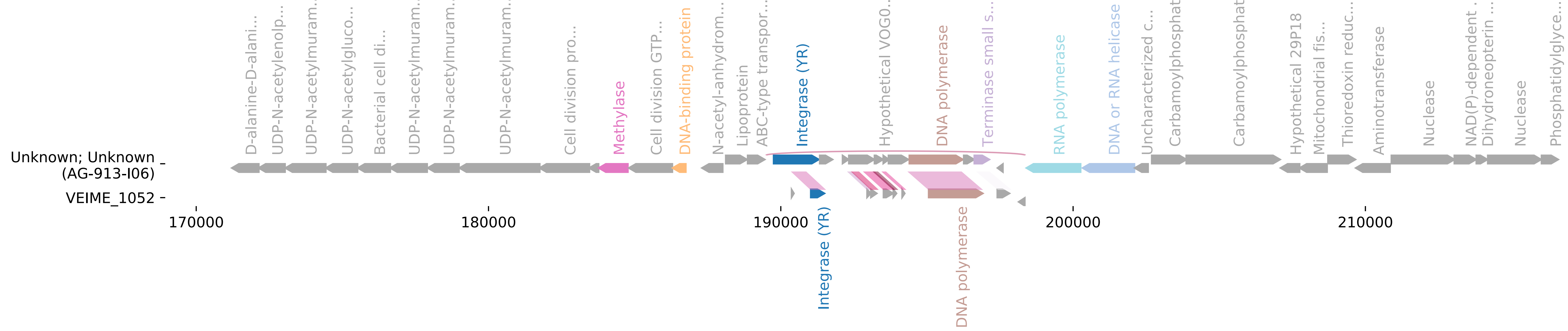

Fig. S3.VEIME_genome_mappings cont'd.

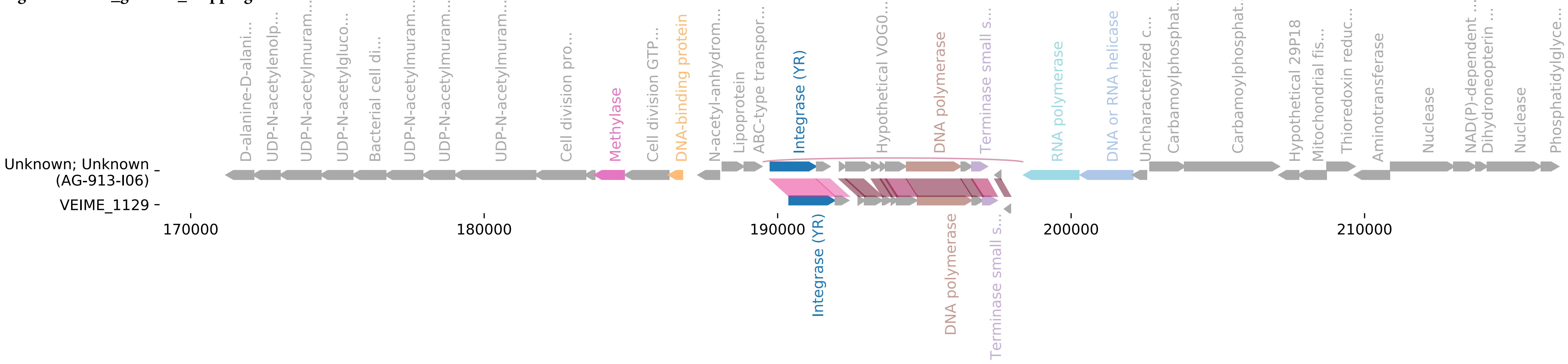

**Fig. S3.VEIME_genome_mappings cont'd.**

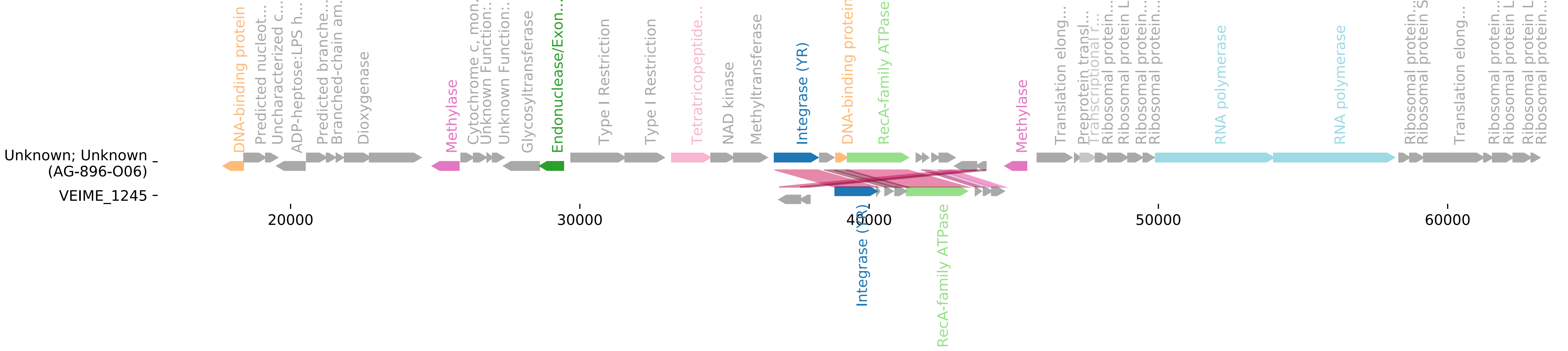

**Fig. S3.VEIME_genome_mappings cont'd.**

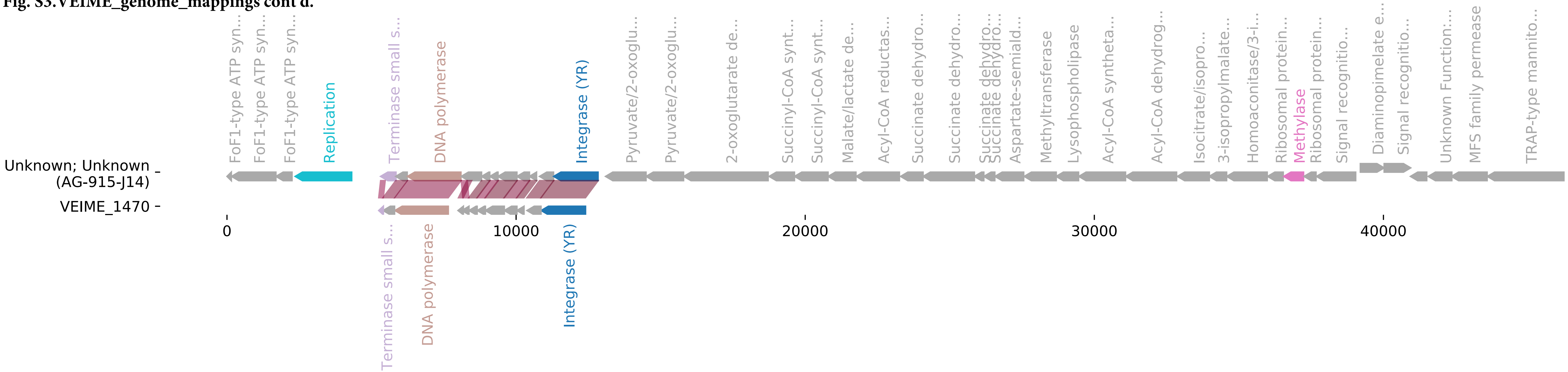

Fig. S3.VEIME_genome_mappings cont'd.

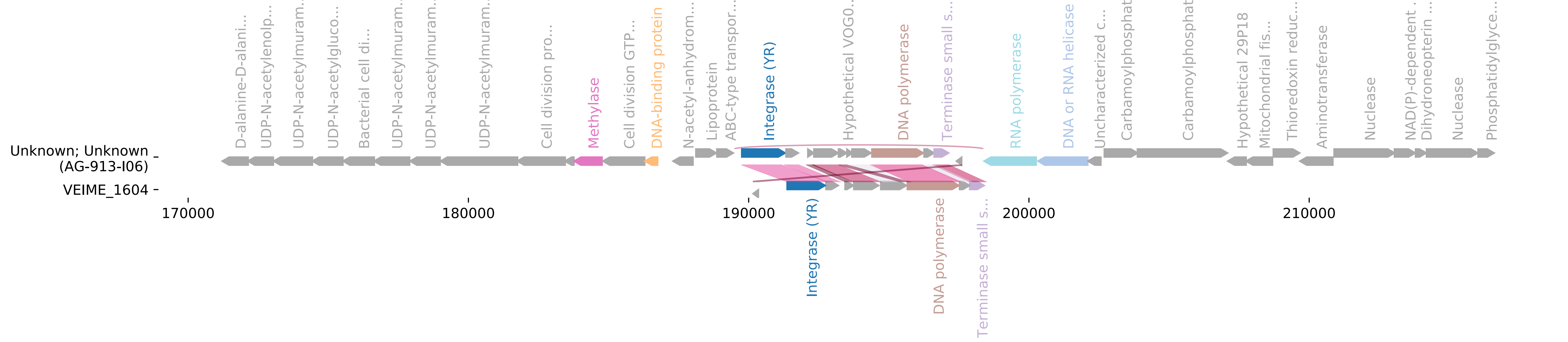

**Fig. S3.VEIME_genome_mappings cont'd.**

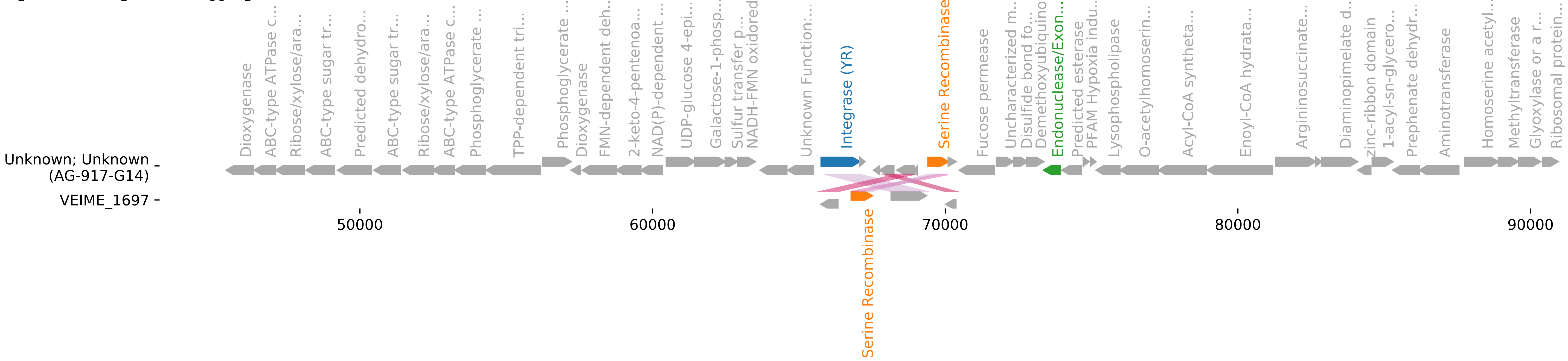

**Fig. S3.VEIME_genome_mappings cont'd.**

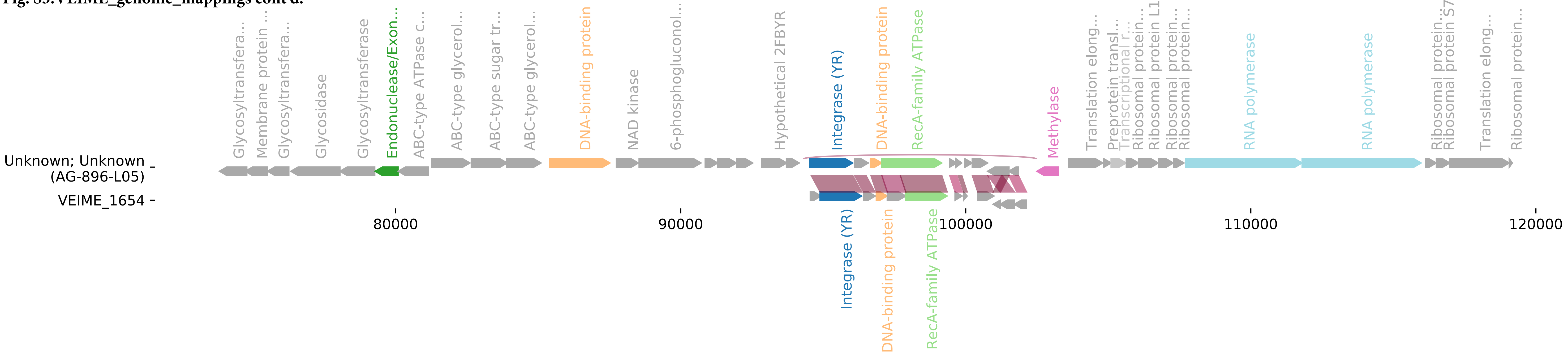

#### Figure S4: VEIME Genome Maps and annotations organized by tyrosine Integrase clades

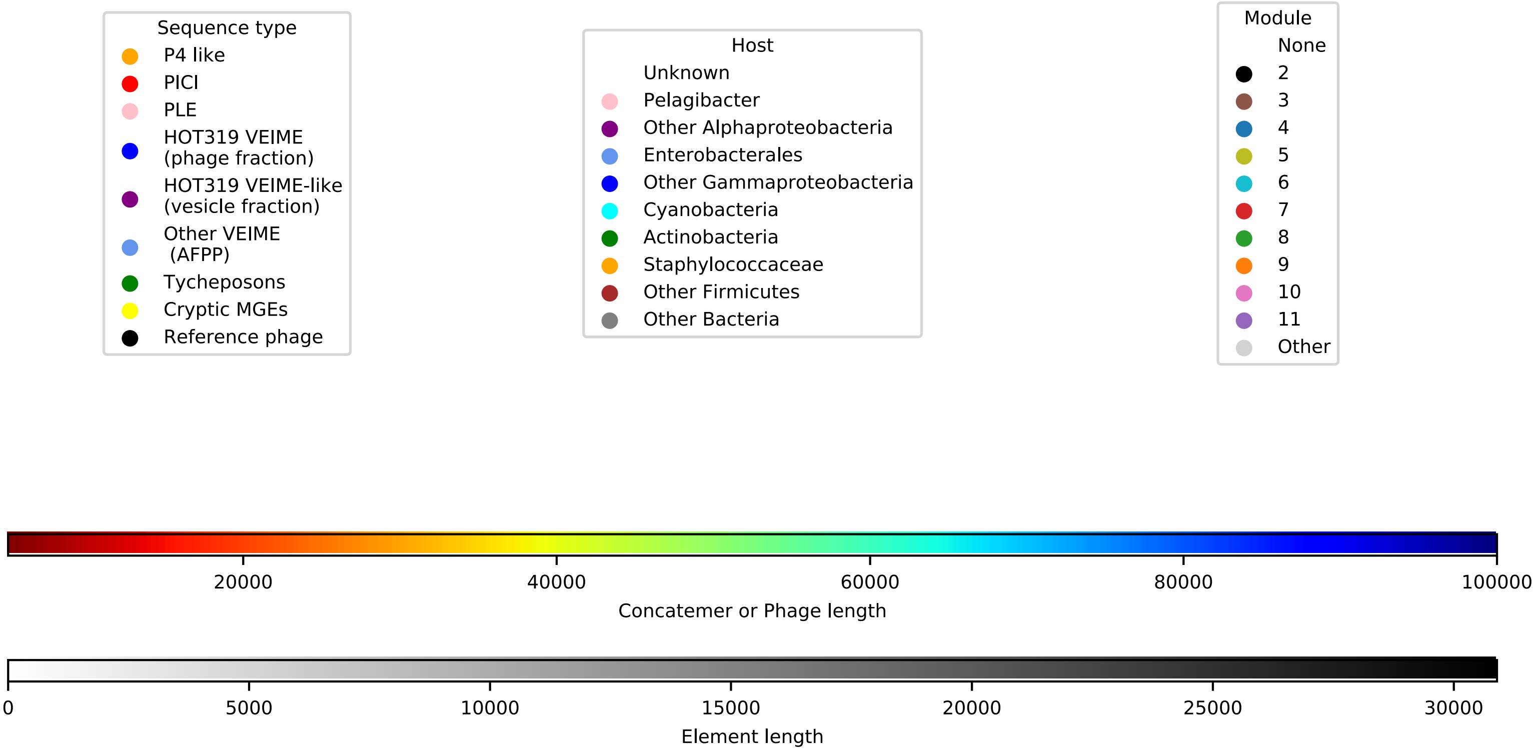


Figure S4: Integrase Tree Genome Maps
Gene annotation alignments (as in Figure S2) are drawn for selected clades from the integrase tree (Figure 3). Plot layouts are the same as Figure S2 except clustering dendrograms are replaced with maxium likelihood trees and moved to the lower left. Clade locations in larger tree are draw on the last page.

**Figure S4: VEIME Genome Maps and Annotations Organized by Tyrosine Integrase Clades, continued**

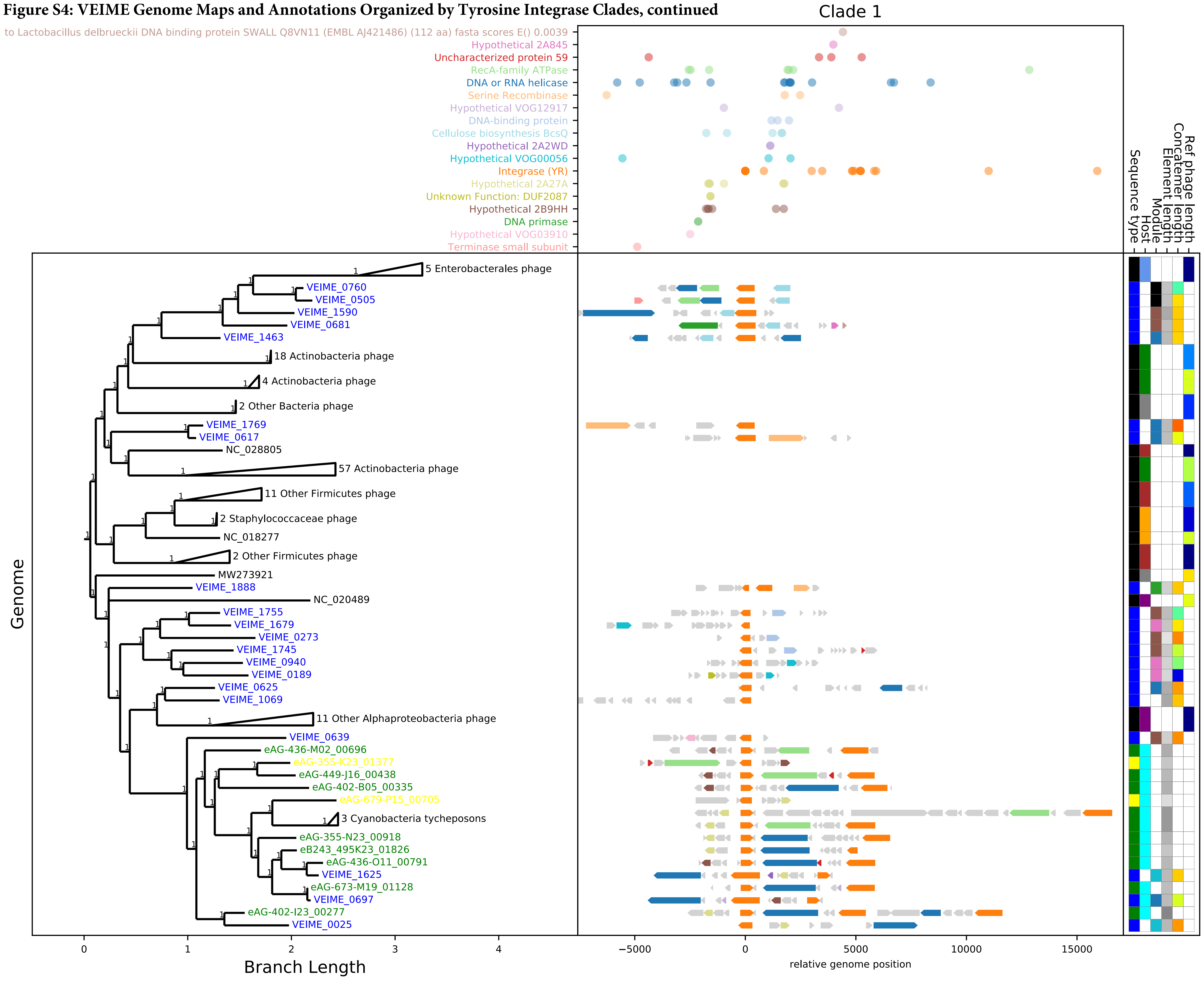

Figure S4: VEIME Genome Maps and annotations organized by tyrosine Integrase clades- continued

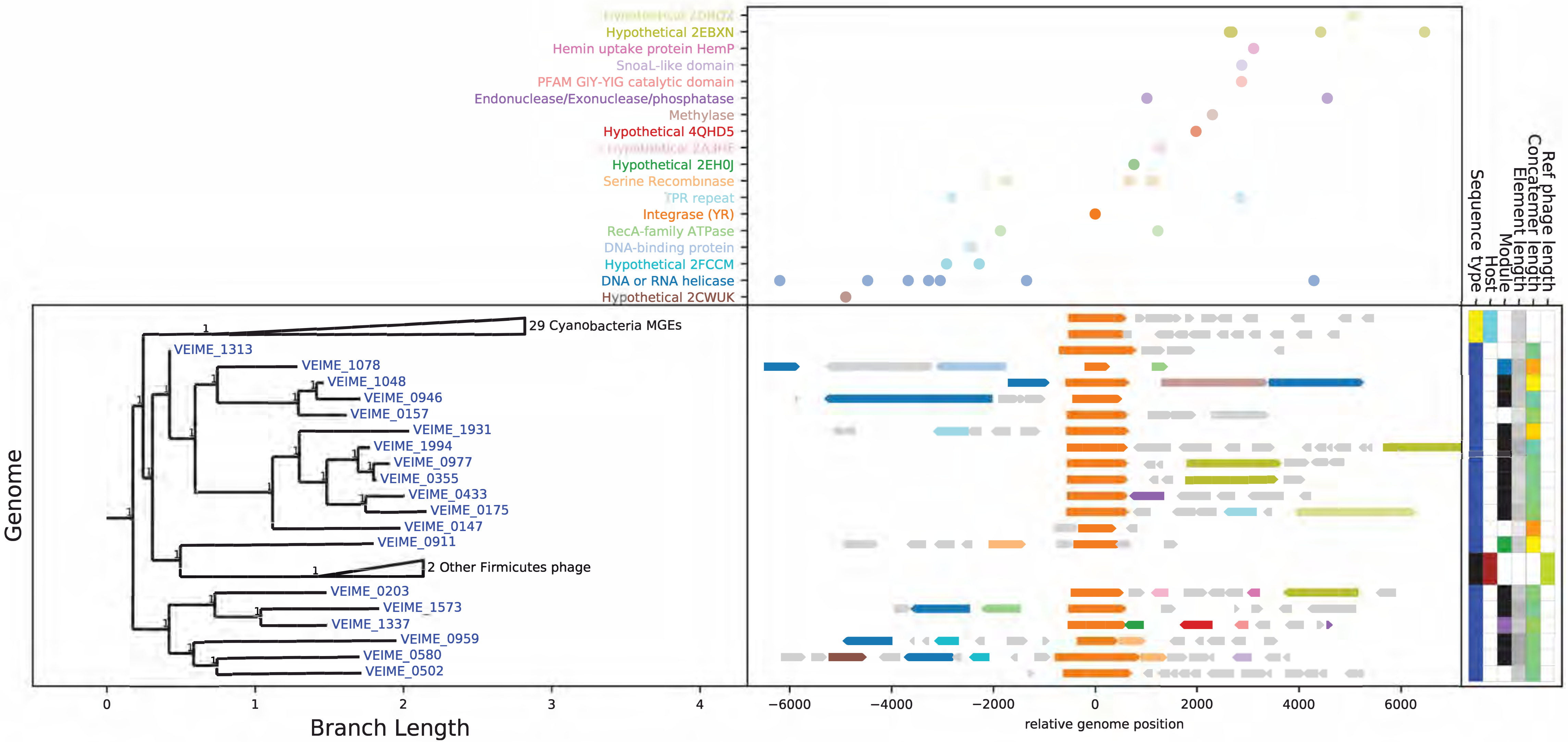

**Figure S4: VEIME Genome Maps and Annotations Organized by Tyrosine Integrase Clades, continued**

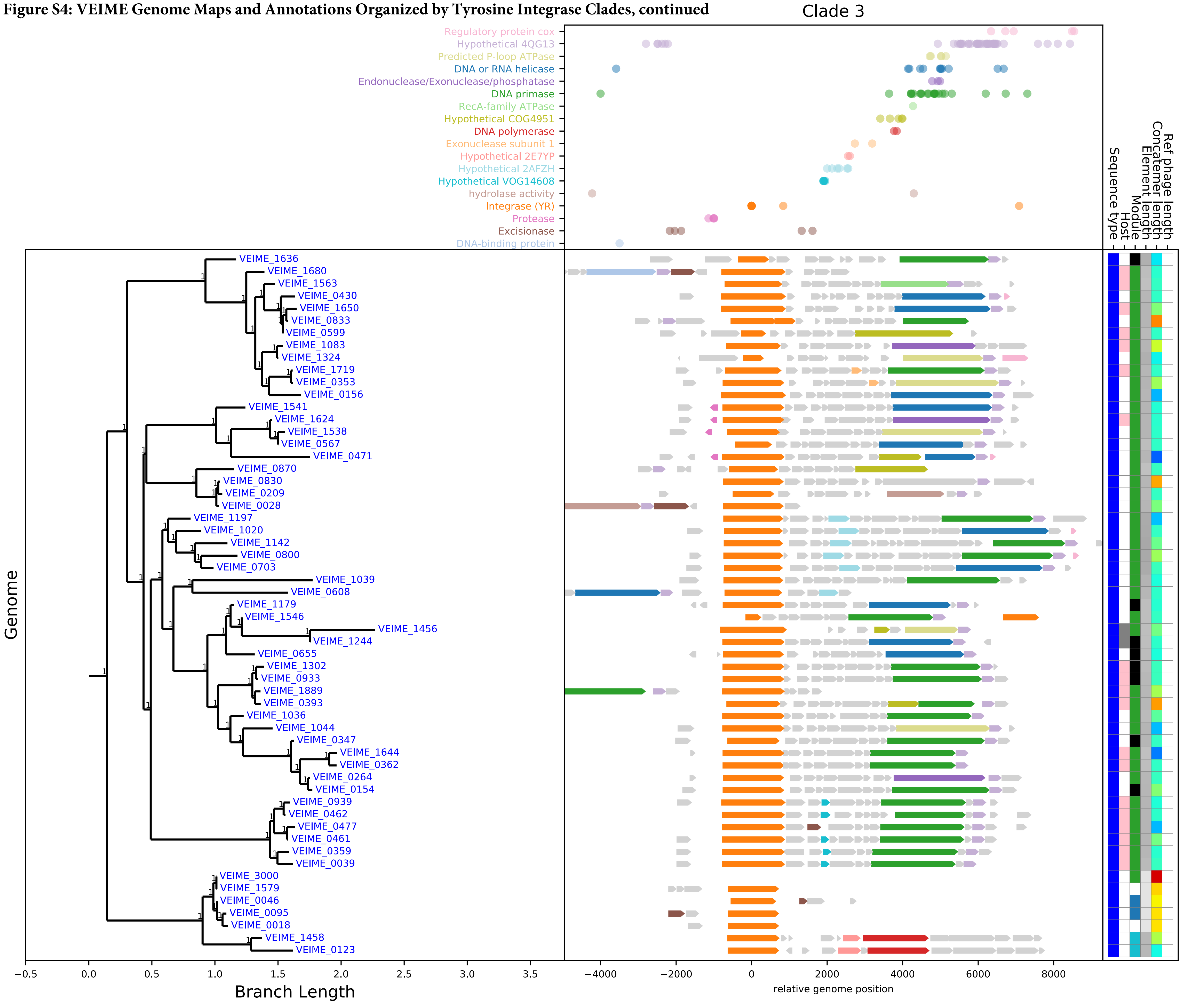

**Figure S4: VEIME Genome Maps and Annotations Organized by Tyrosine Integrase Clades, continued**

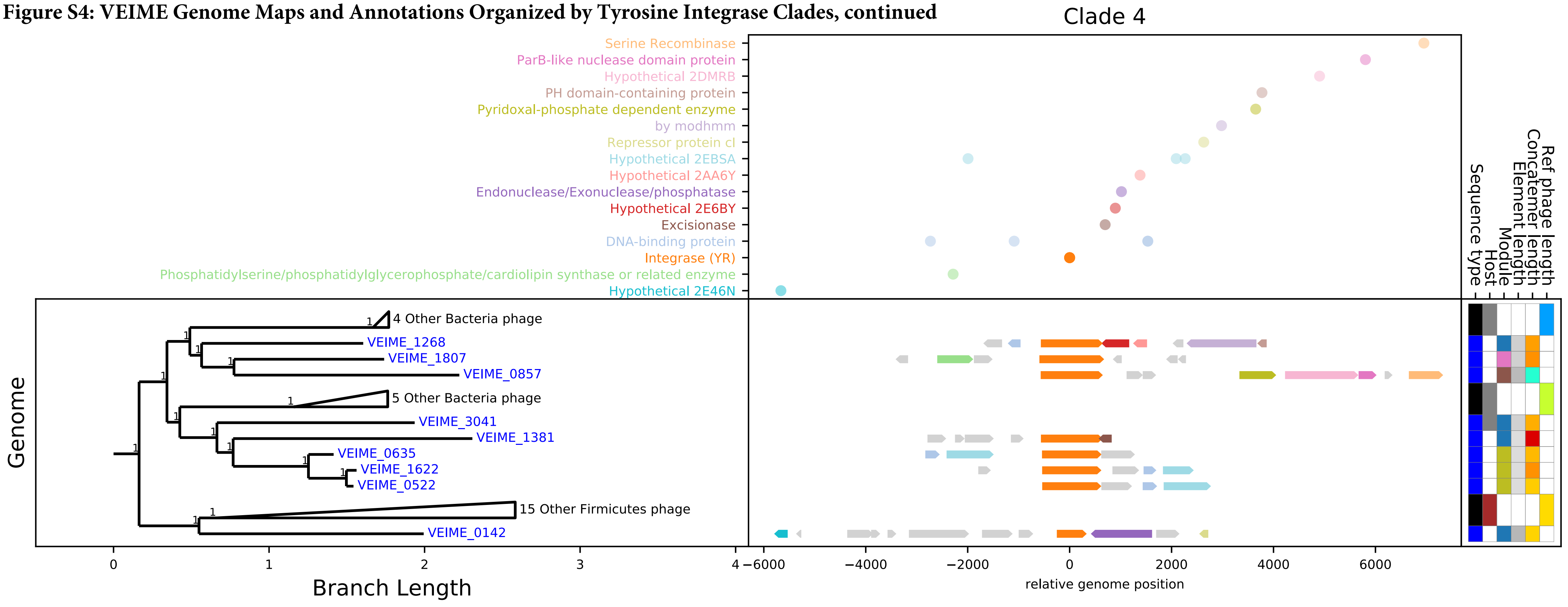

**Figure S4: VEIME Genome Maps and Annotations Organized by Tyrosine Integrase Clades, continued**

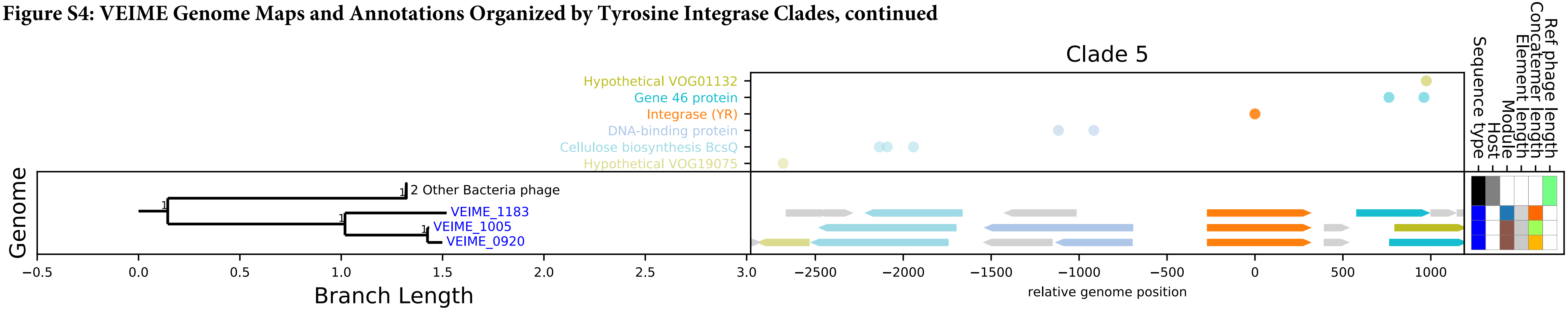

**Figure S4: VEIME Genome Maps and Annotations Organized by Tyrosine Integrase Clades, continued**

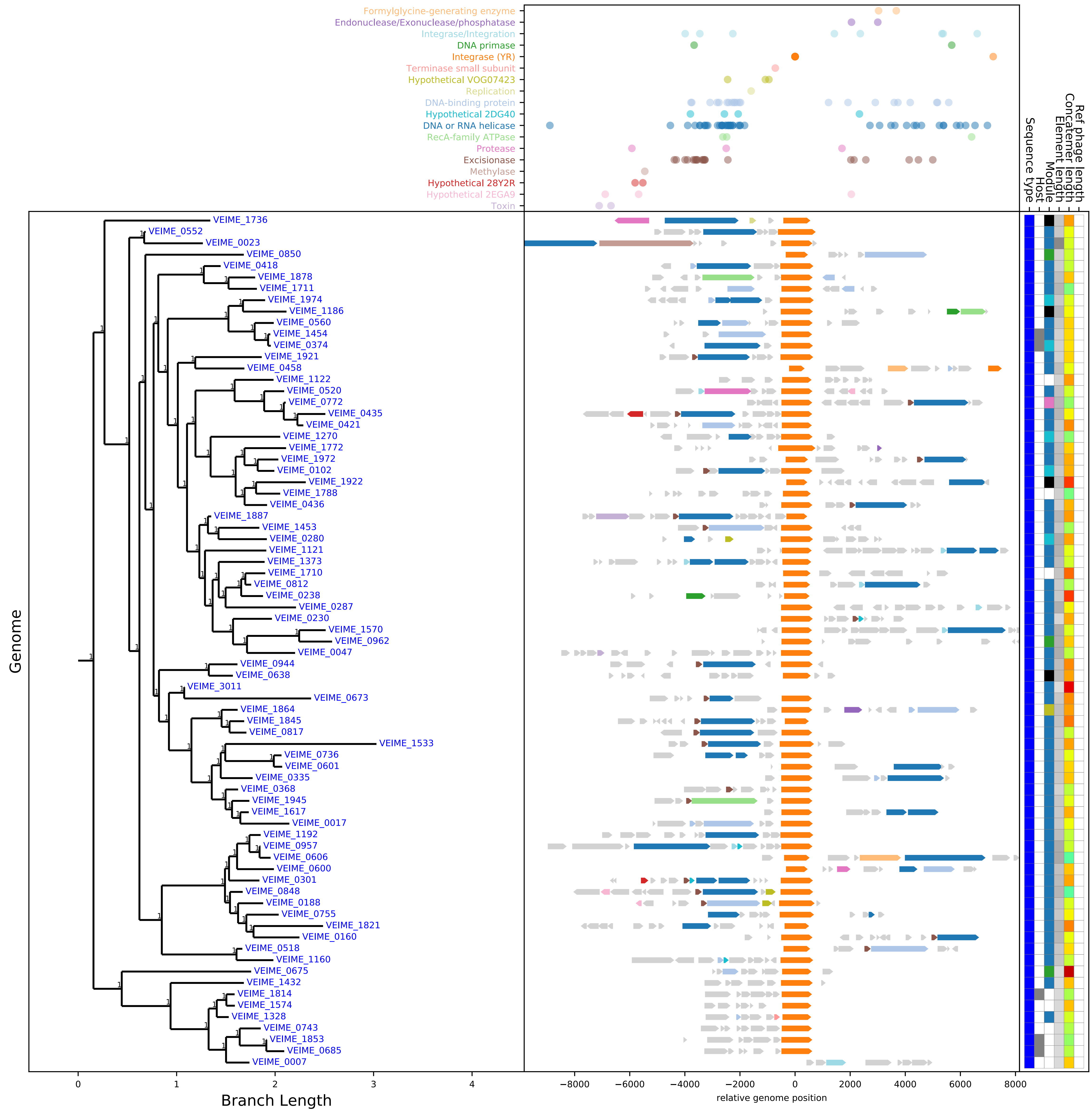

**Figure S4: VEIME Genome Maps and Annotations Organized by Tyrosine Integrase Clades, continued**

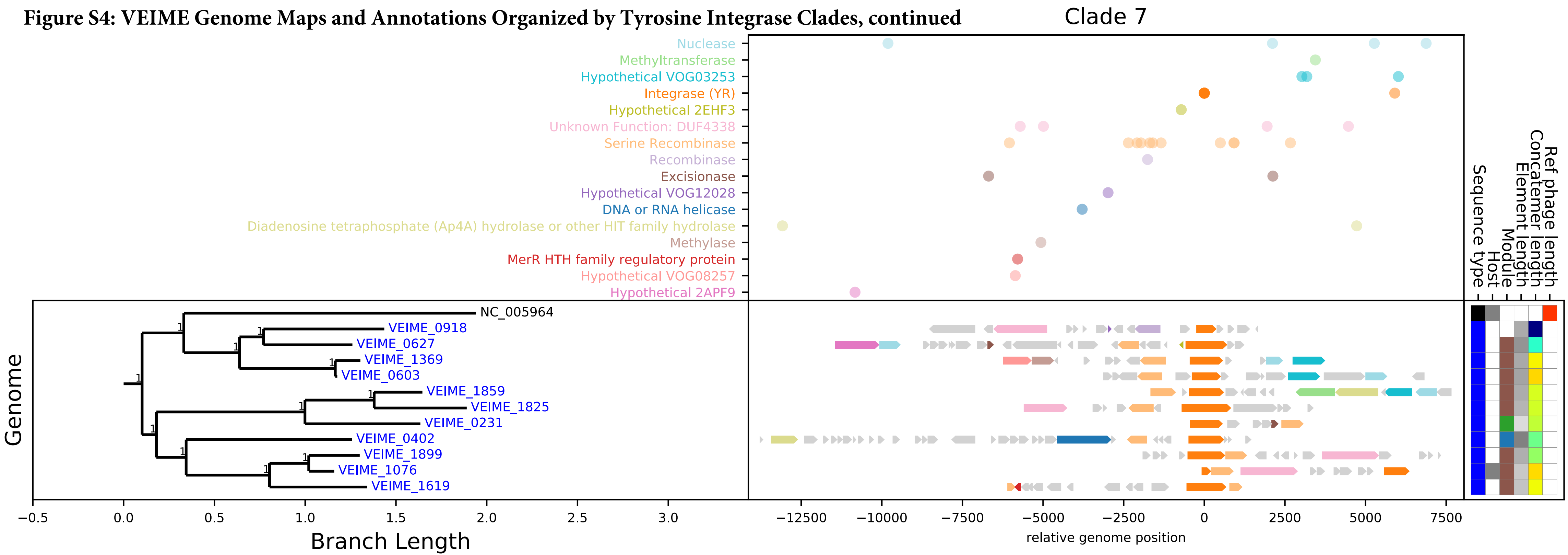

**Figure S4: VEIME Genome Maps and Annotations Organized by Tyrosine Integrase Clades, continued**

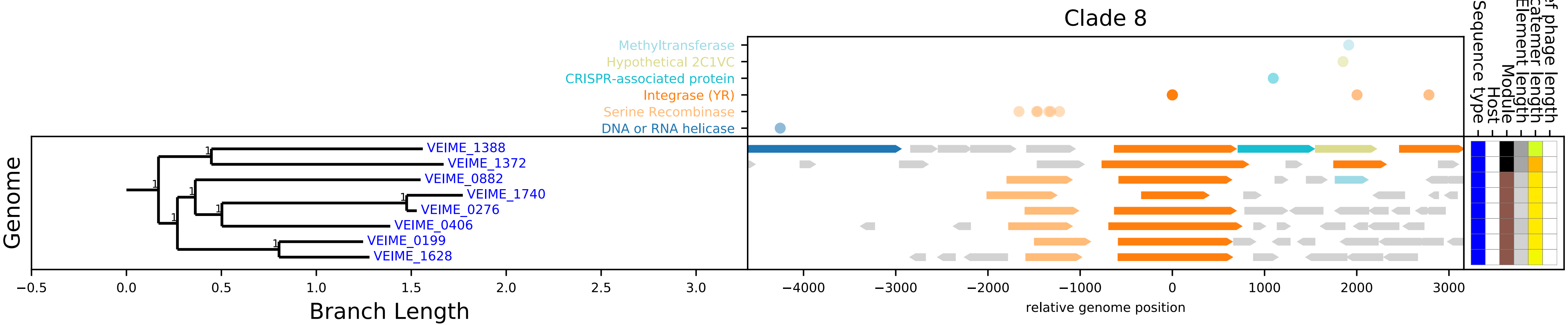

**Figure S4: VEIME Genome Maps and Annotations Organized by Tyrosine Integrase Clades, continued**

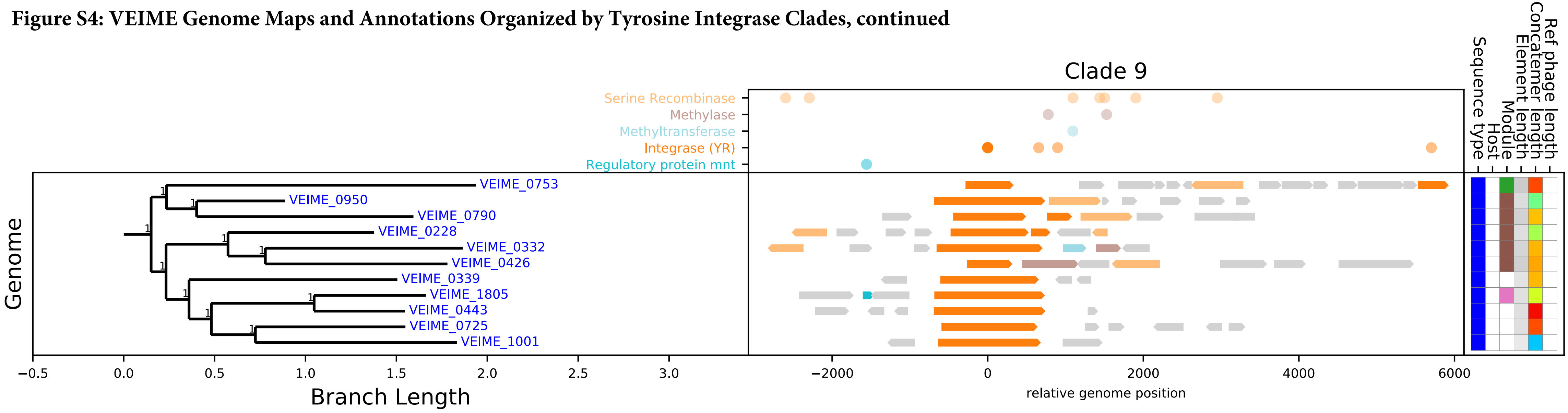

**Figure S4: VEIME Genome Maps and Annotations Organized by Tyrosine Integrase Clades, continued**

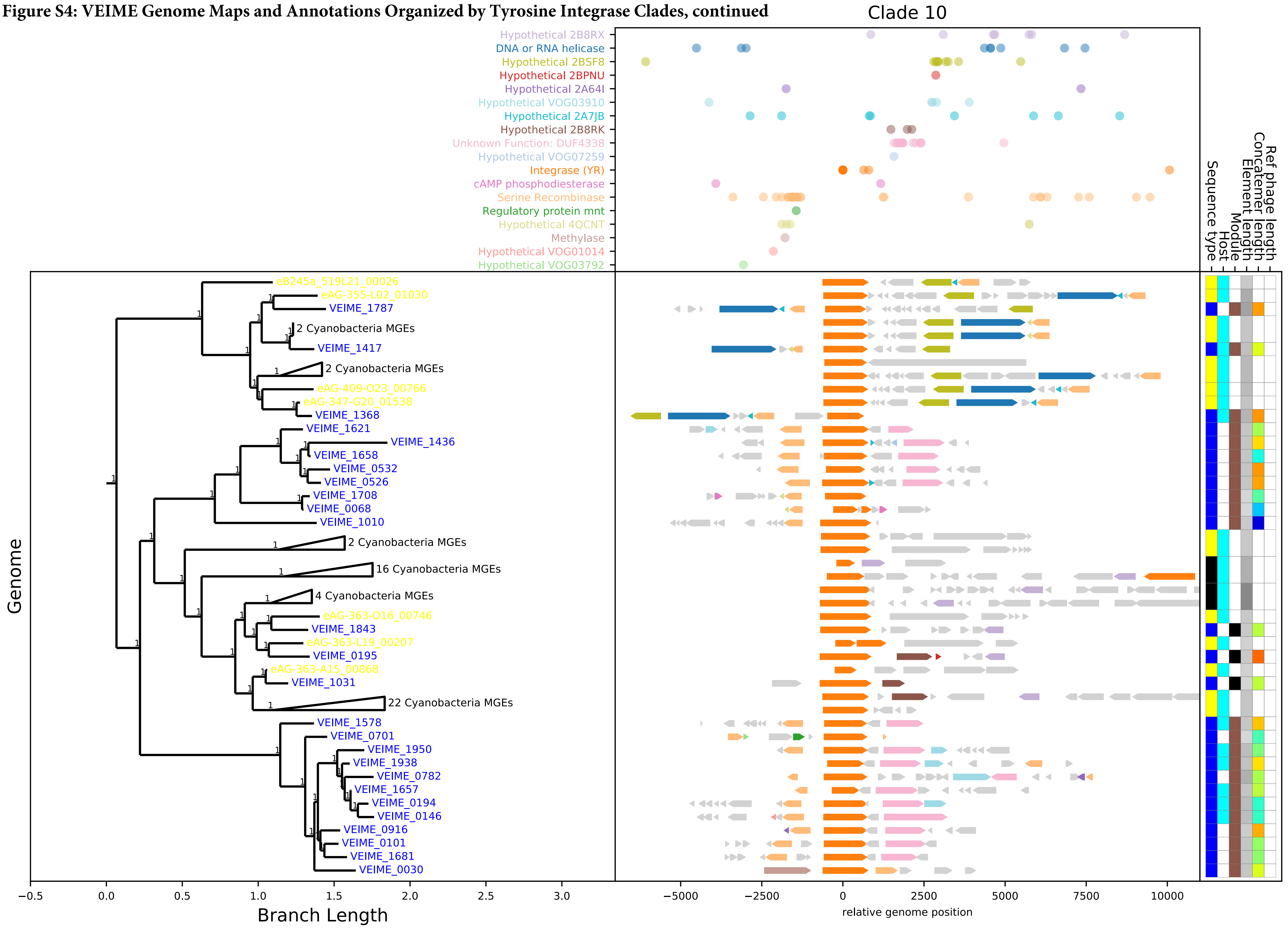

**Figure S4: VEIME Genome Maps and Annotations Organized by Tyrosine Integrase Clades, continued**

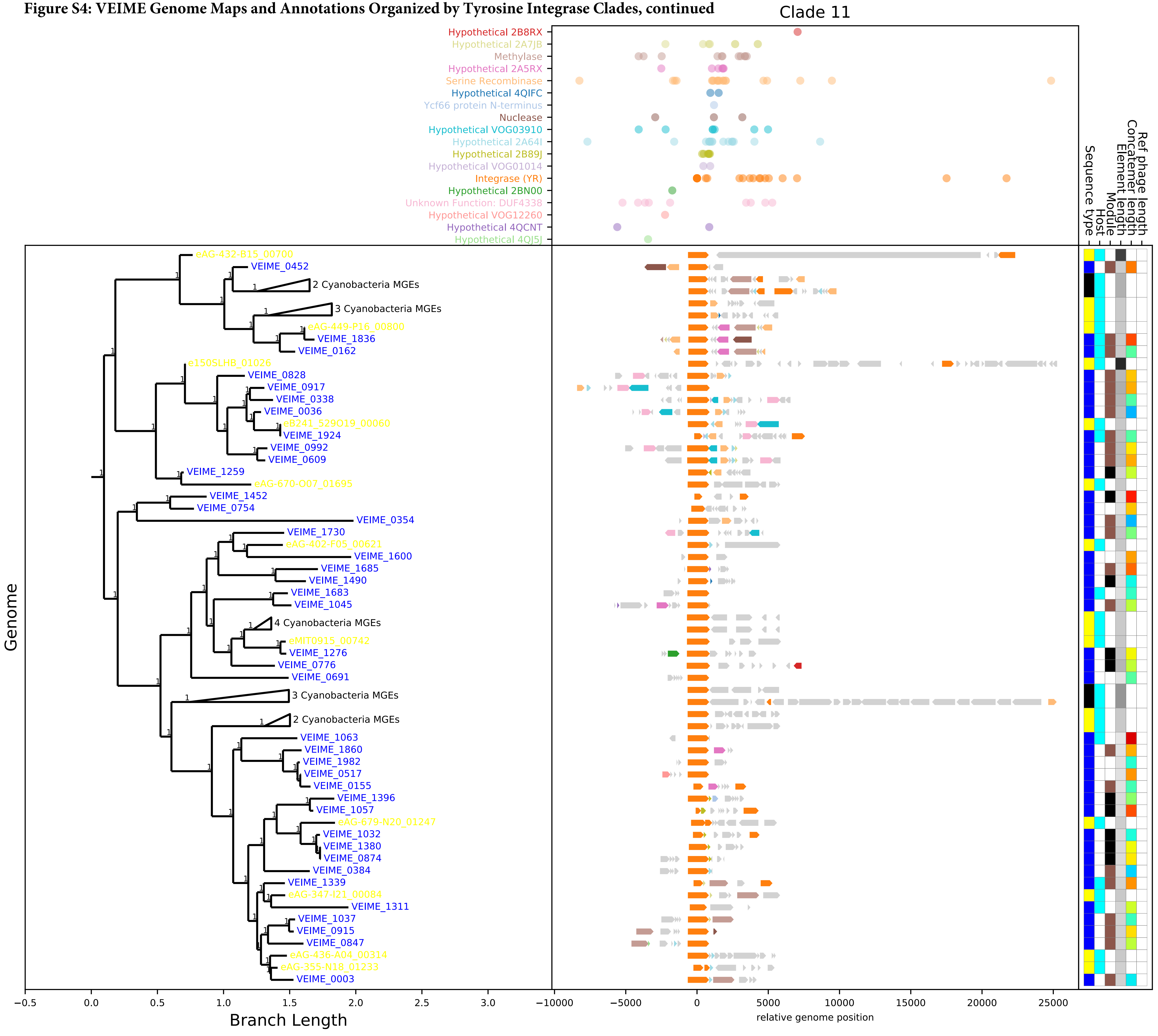

**Figure S4: VEIME Genome Maps and Annotations Organized by Tyrosine Integrase Clades, continued**

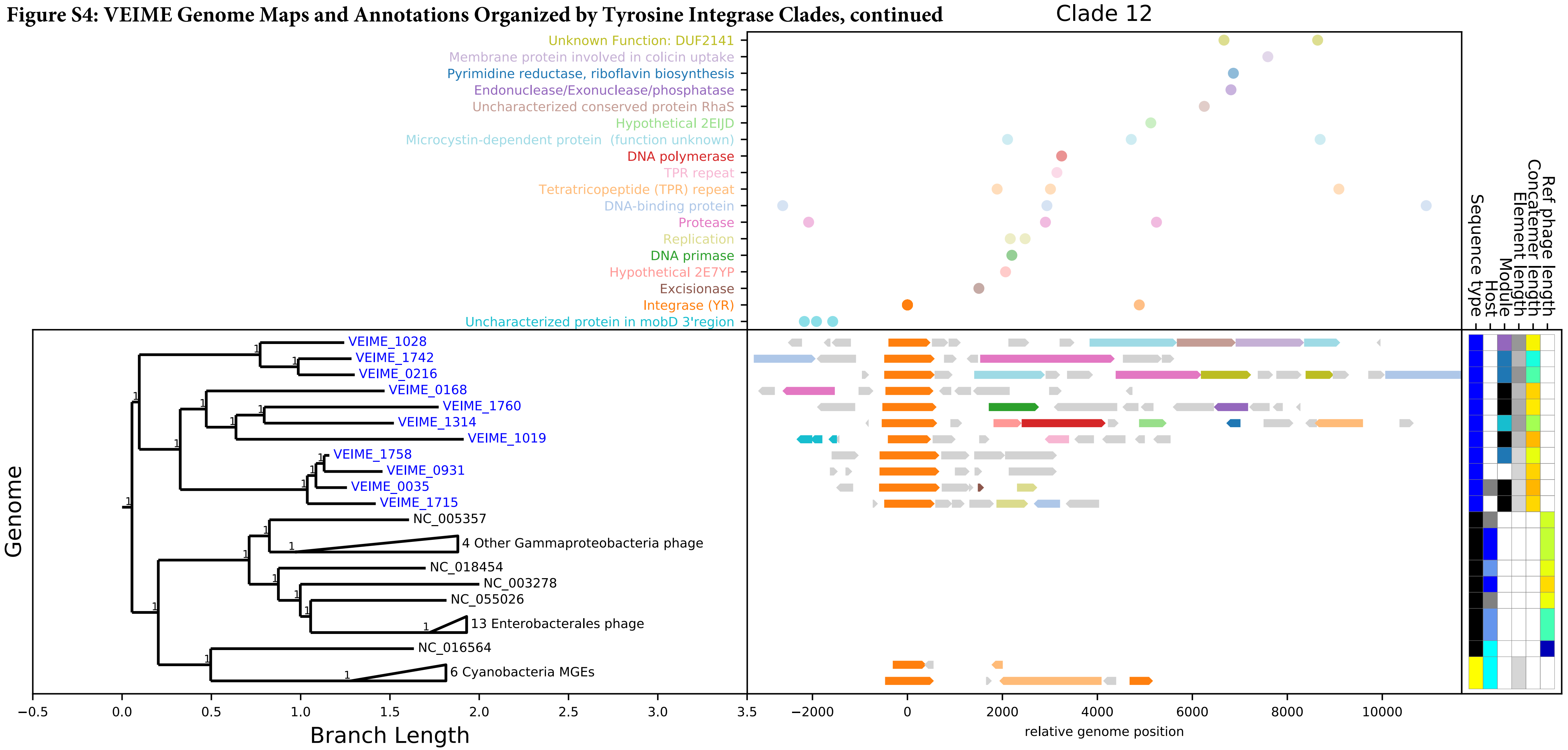

**Figure S4: VEIME Genome Maps and Annotations Organized by Tyrosine Integrase Clades, continued**

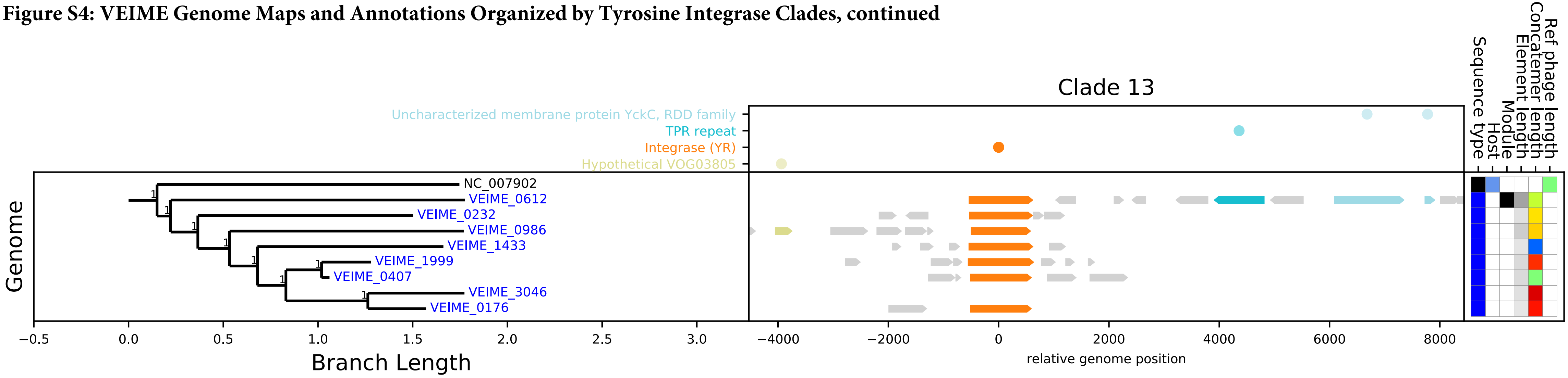

**Figure S4: VEIME Genome Maps and Annotations Organized by Tyrosine Integrase Clades, continued**

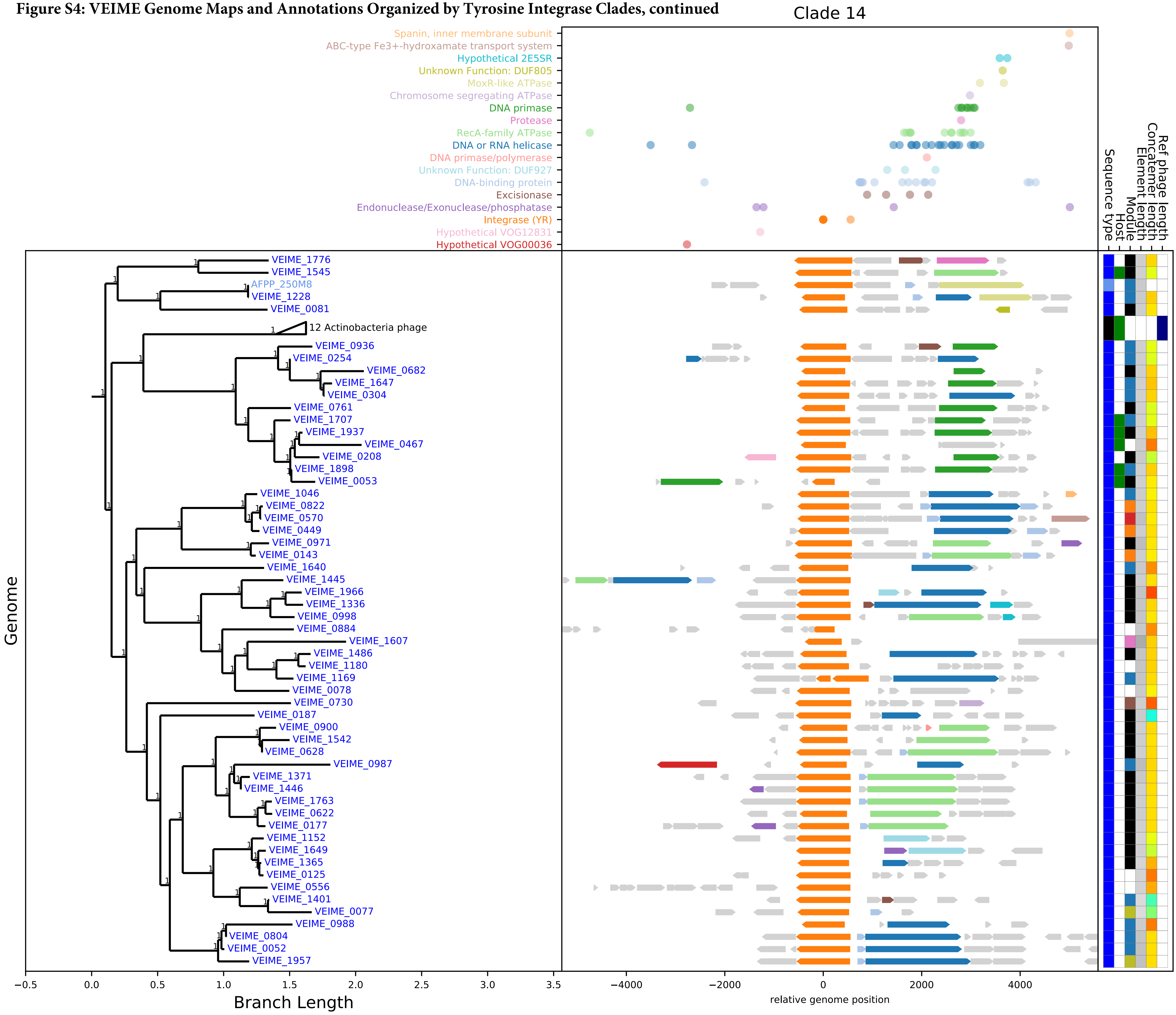

**Figure S4: VEIME Genome Maps and Annotations Organized by Tyrosine Integrase Clades, continued**

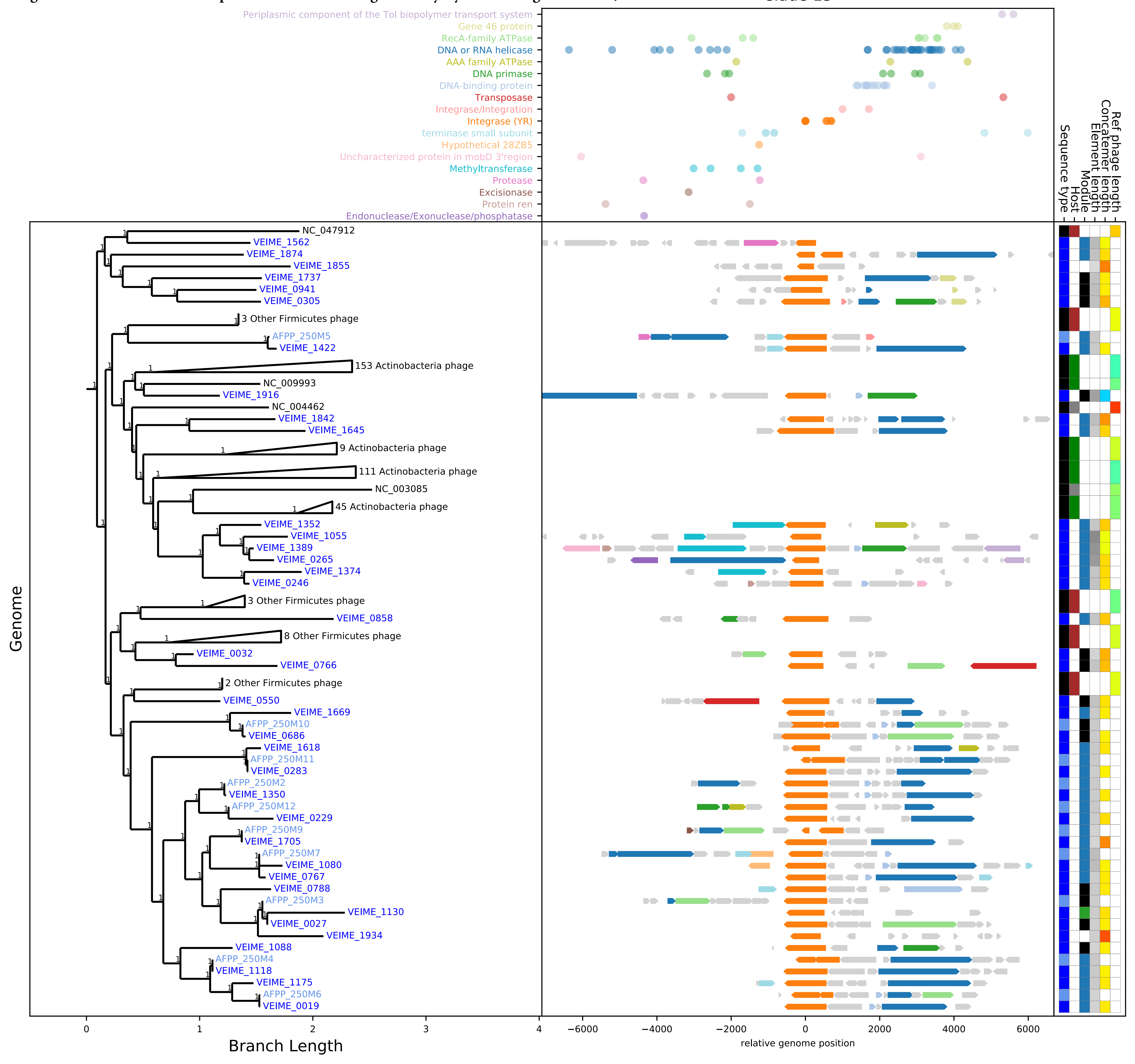

#### Figure S4: VEIME Genome Maps and Annotations Organized by Tyrosine Integrase Clades, continued

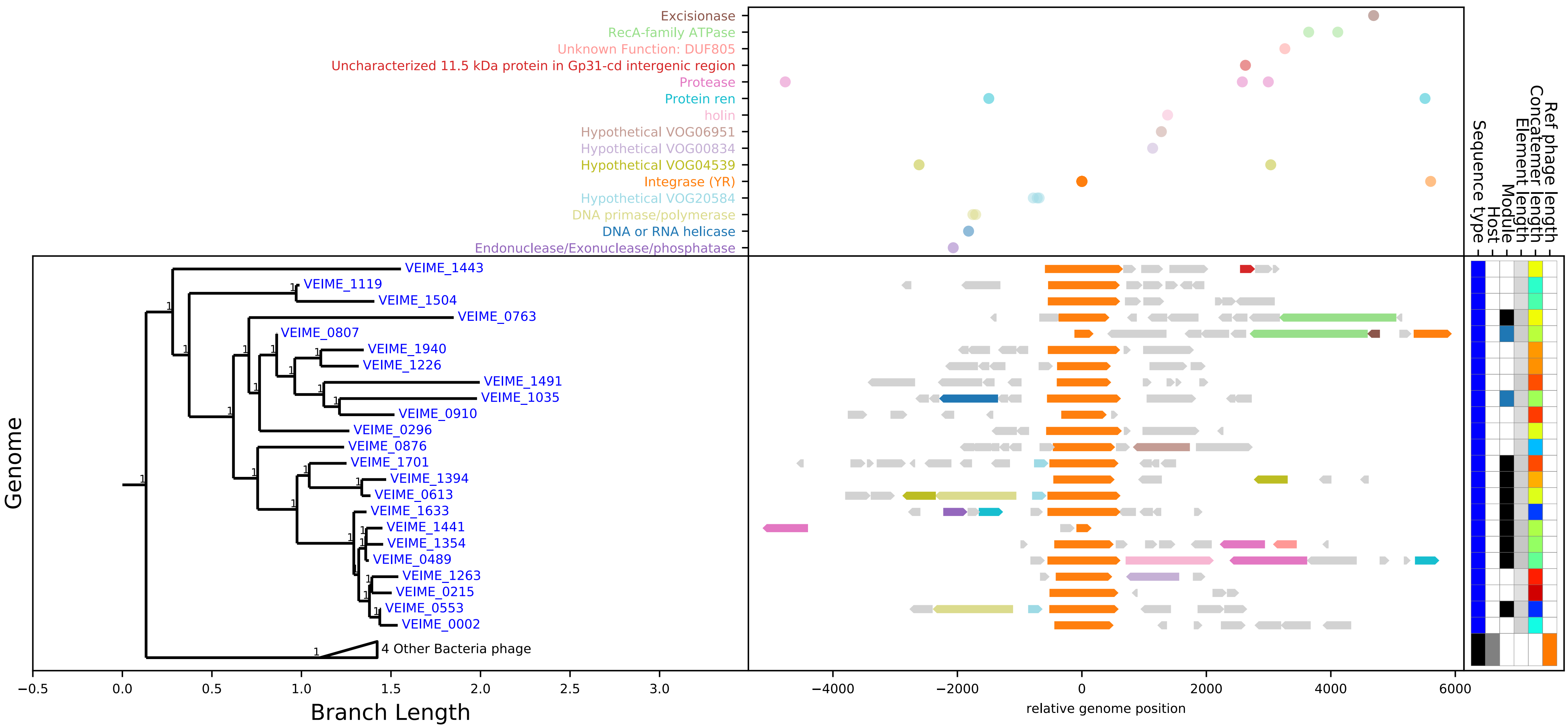

**Figure S4: VEIME Genome Maps and Annotations Organized by Tyrosine Integrase Clades, continued**

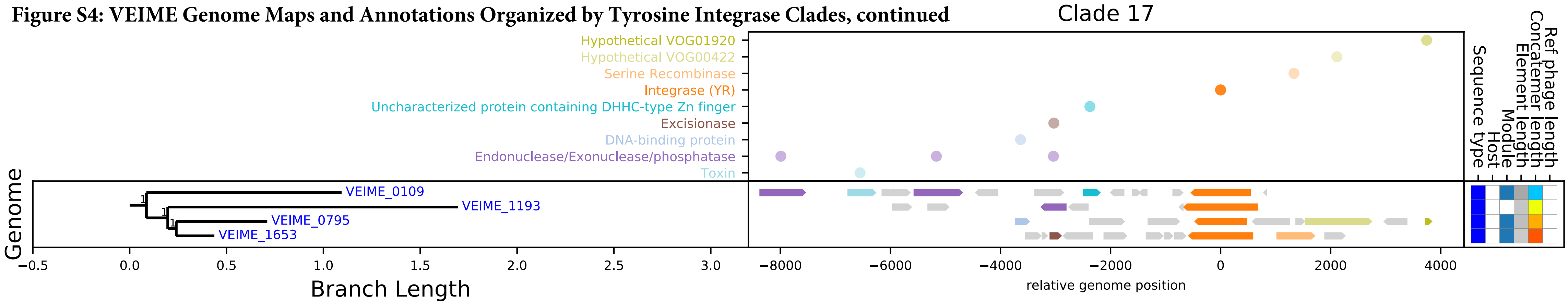

**Figure S4: VEIME Genome Maps and Annotations Organized by Tyrosine Integrase Clades, continued**

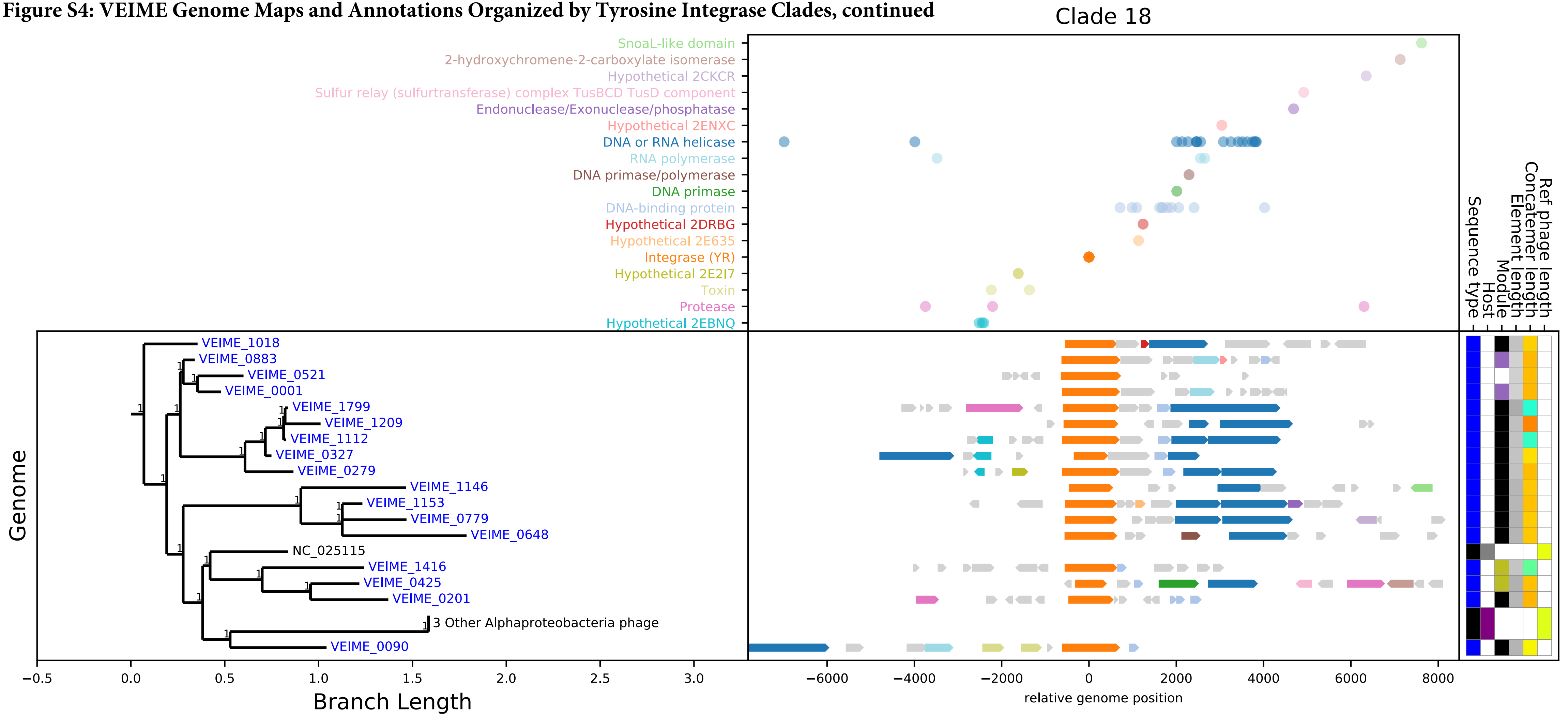

**Figure S4: VEIME Genome Maps and Annotations Organized by Tyrosine Integrase Clades, continued**

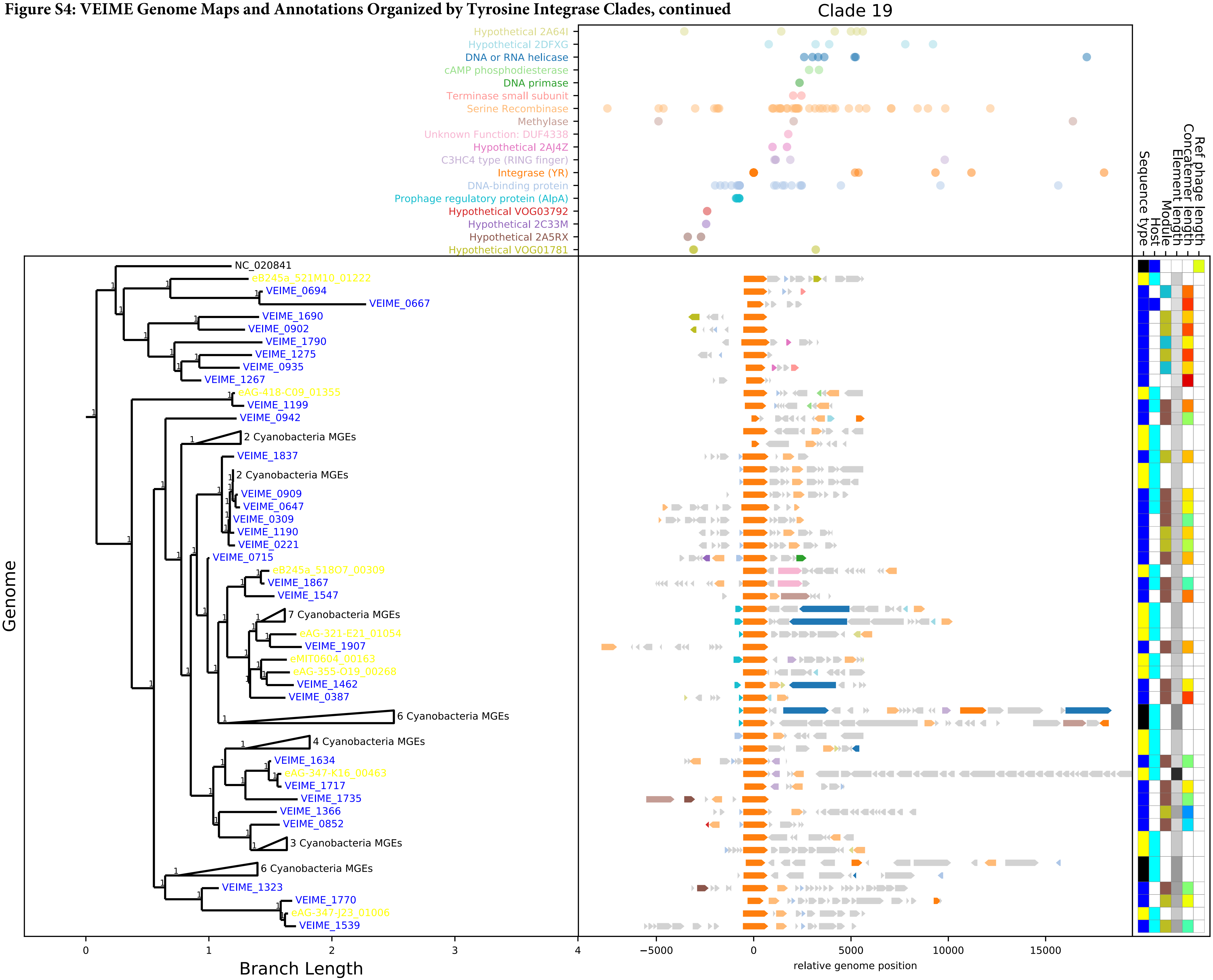

**Figure S4: VEIME Genome Maps and Annotations Organized by Tyrosine Integrase Clades, continued**

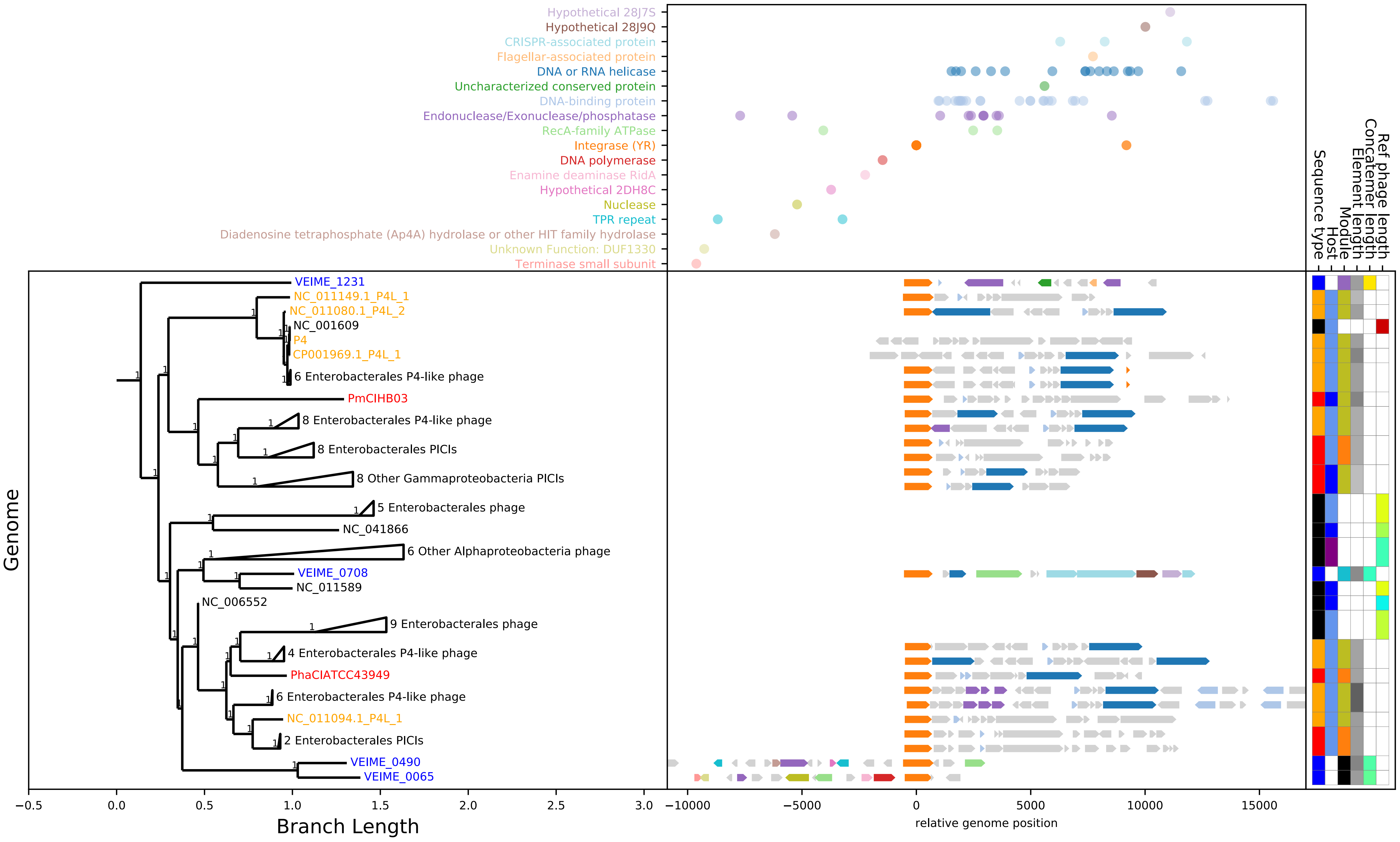

**Figure S4: VEIME Genome Maps and Annotations Organized by Tyrosine Integrase Clades, continued**

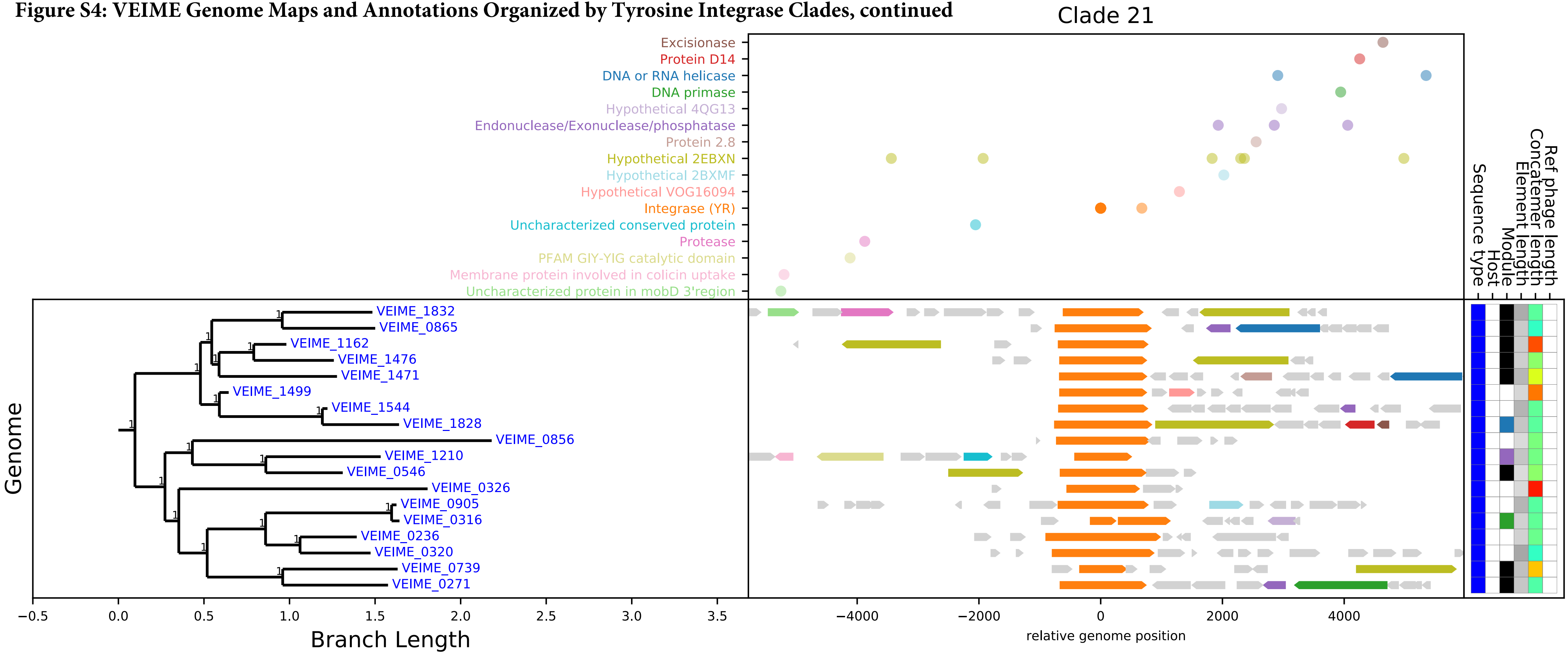

**Figure S4: VEIME Genome Maps and Annotations Organized by Tyrosine Integrase Clades, continued**

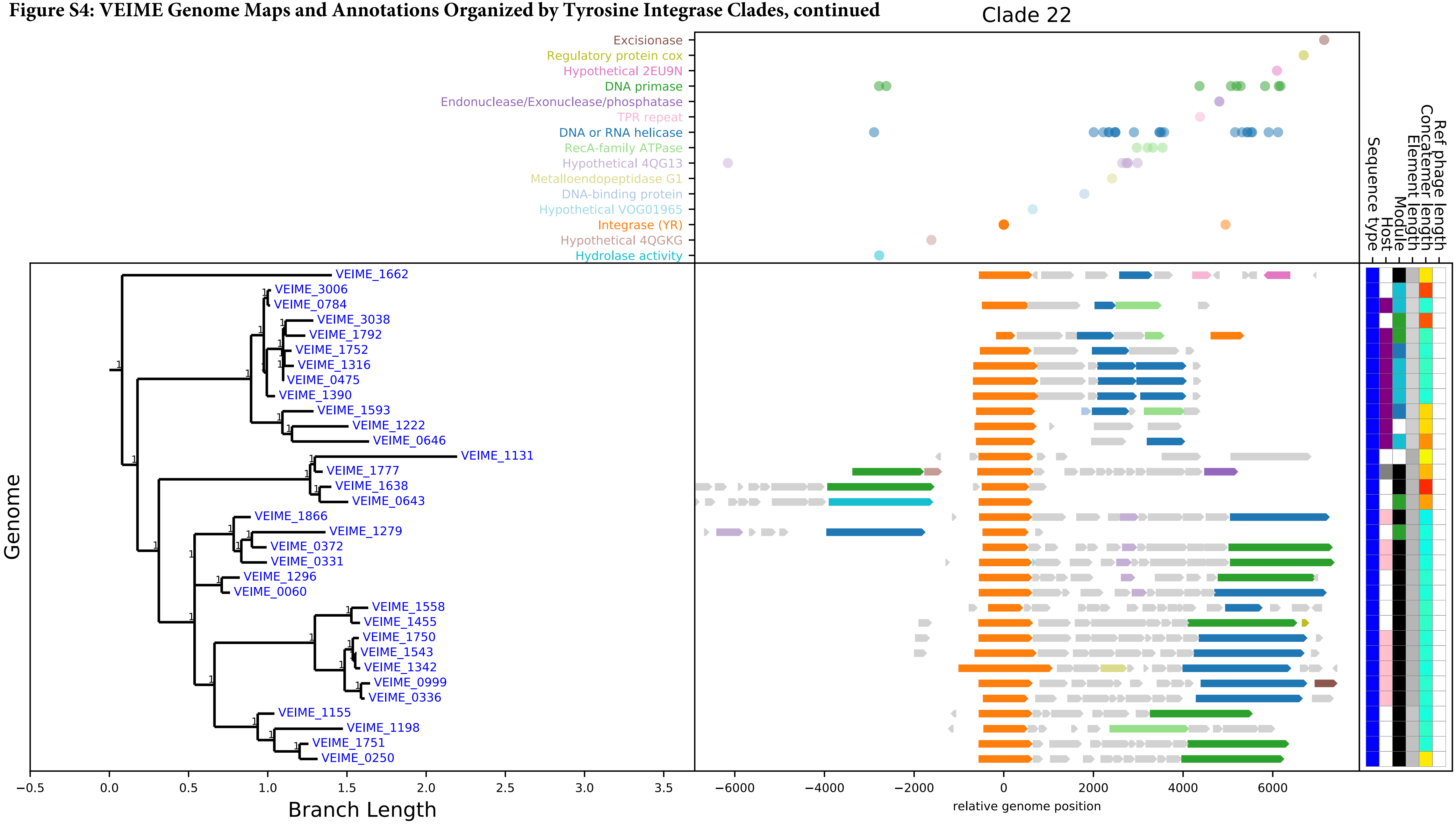

**Figure S4: VEIME Genome Maps and Annotations Organized by Tyrosine Integrase Clades, continued**

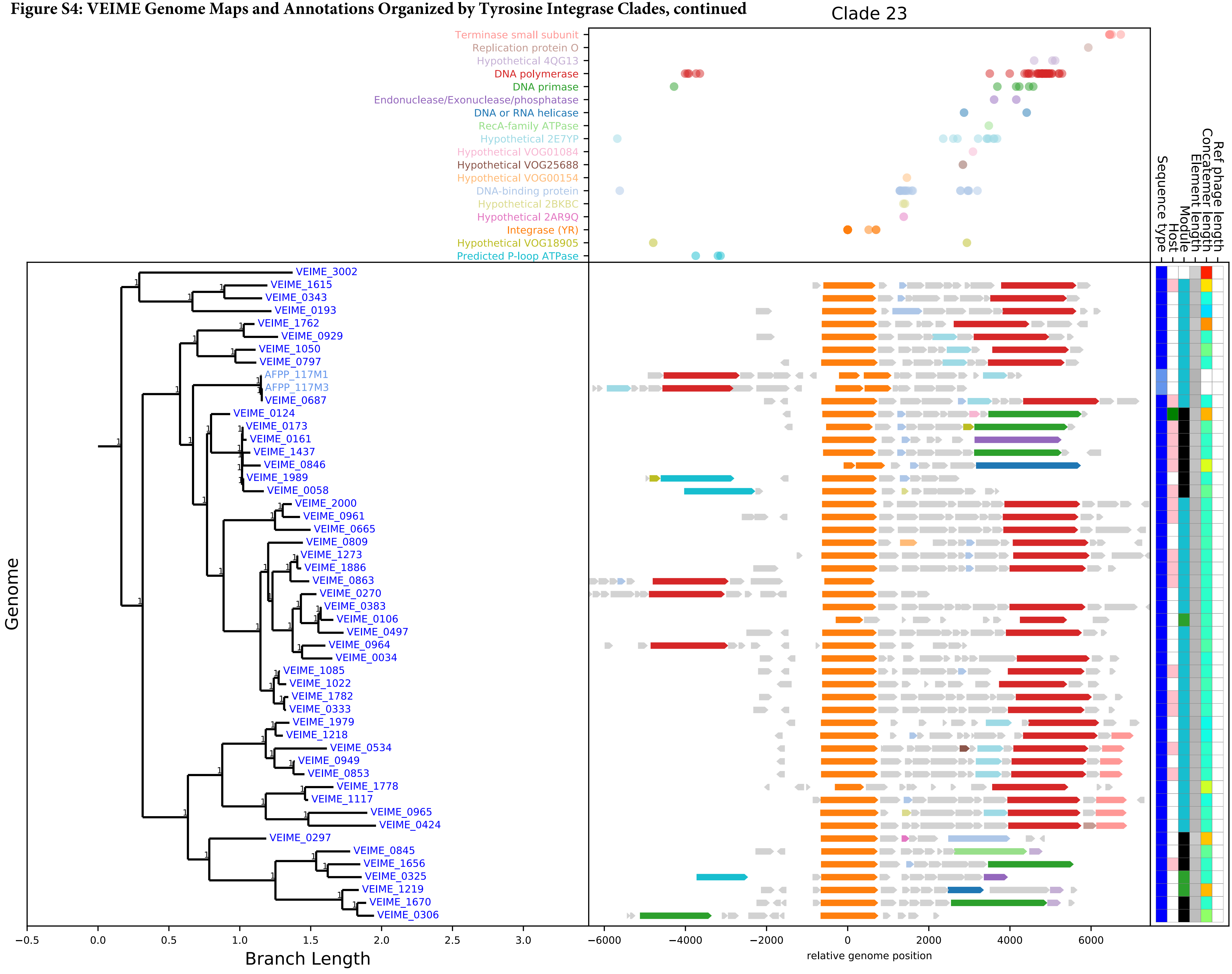

**Figure S4: VEIME Genome Maps and Annotations Organized by Tyrosine Integrase Clades, continued**

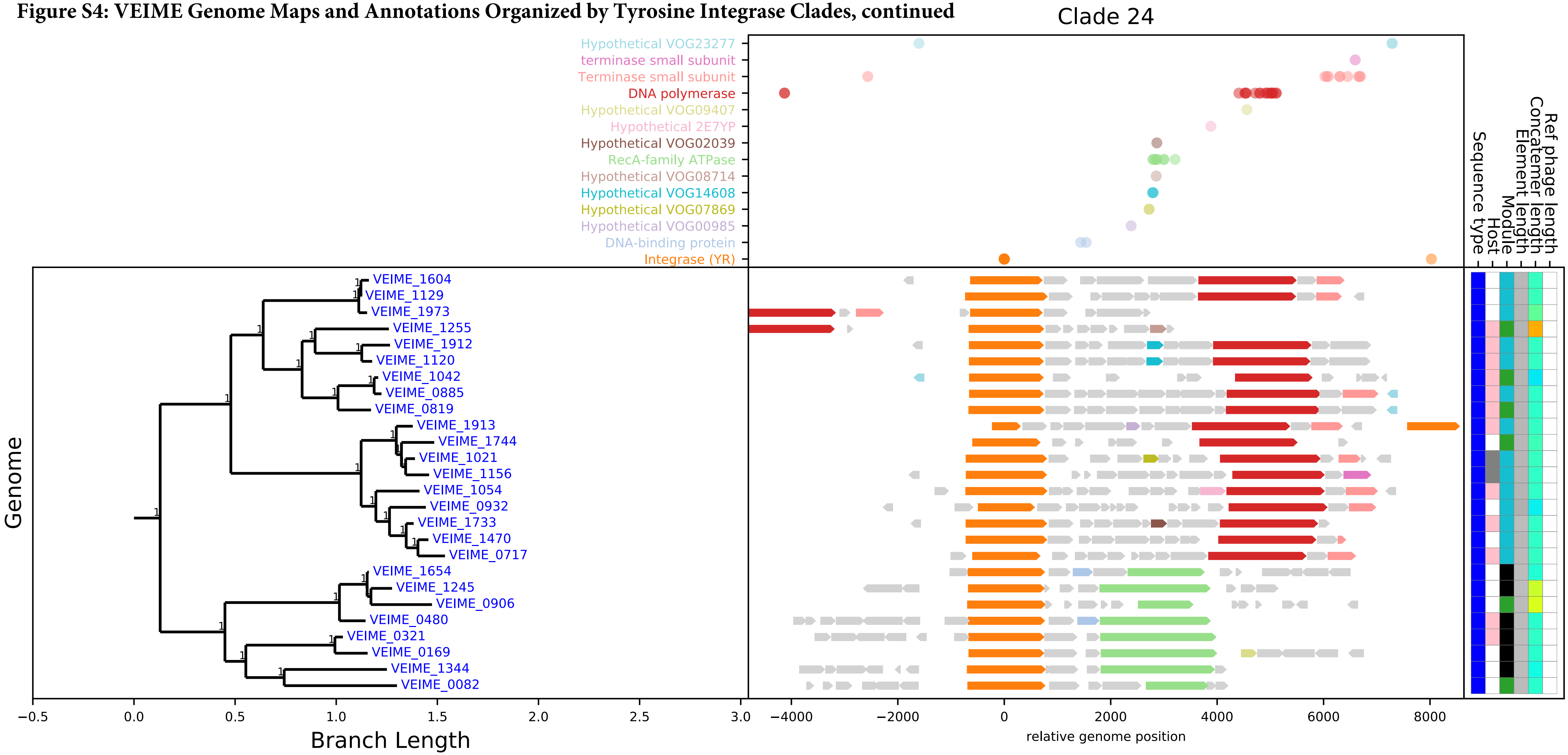

**Figure S4: VEIME Genome Maps and Annotations Organized by Tyrosine Integrase Clades, continued**

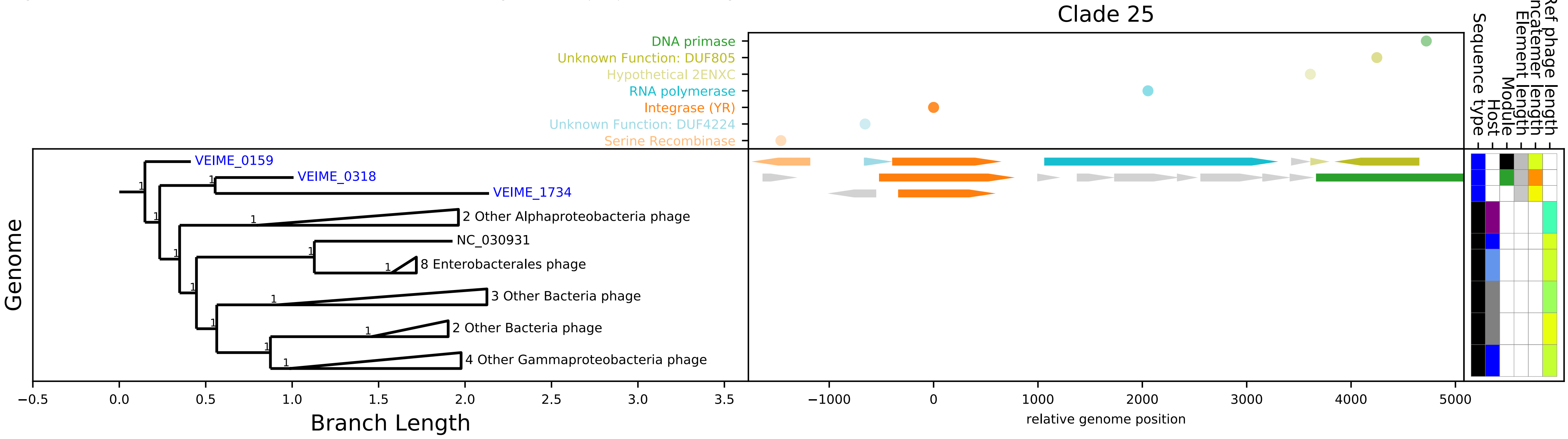

**Figure S4: VEIME Genome Maps and Annotations Organized by Tyrosine Integrase Clades, continued**

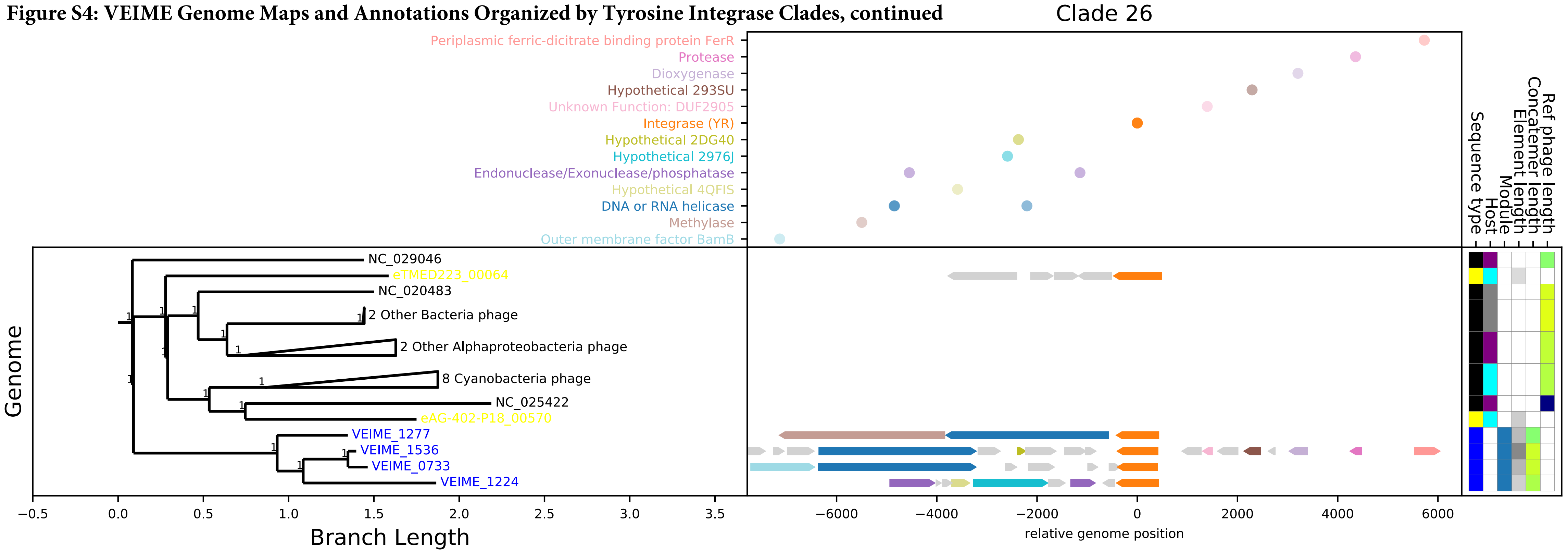

**Figure S4: VEIME Genome Maps and Annotations Organized by Tyrosine Integrase Clades, continued**

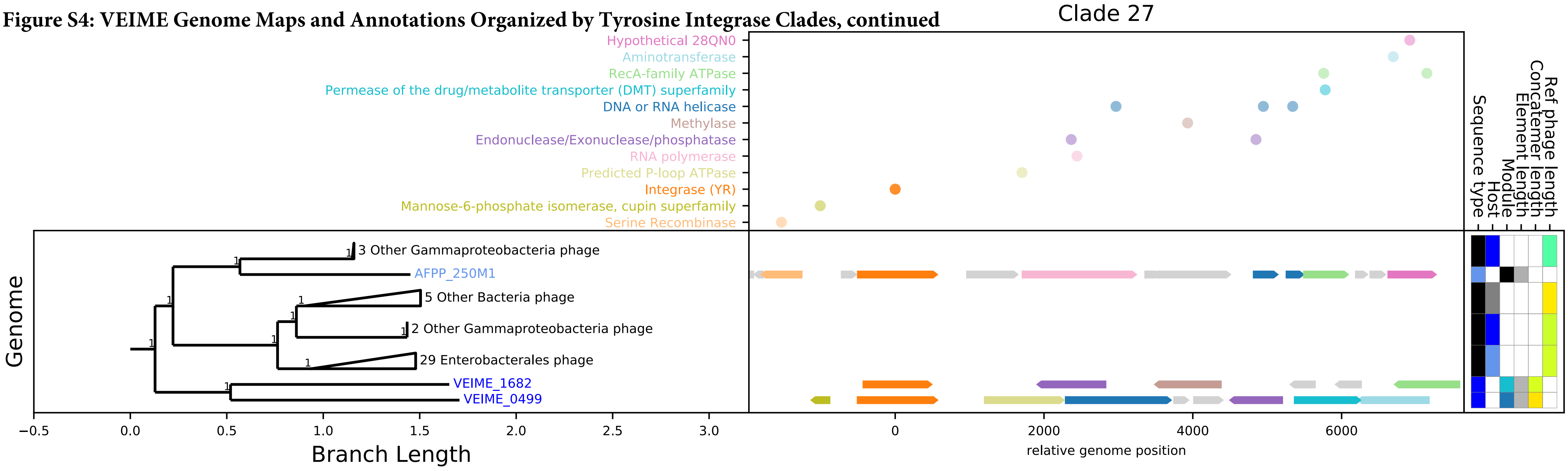

**Figure S4: VEIME Genome Maps and Annotations Organized by Tyrosine Integrase Clades, continued**

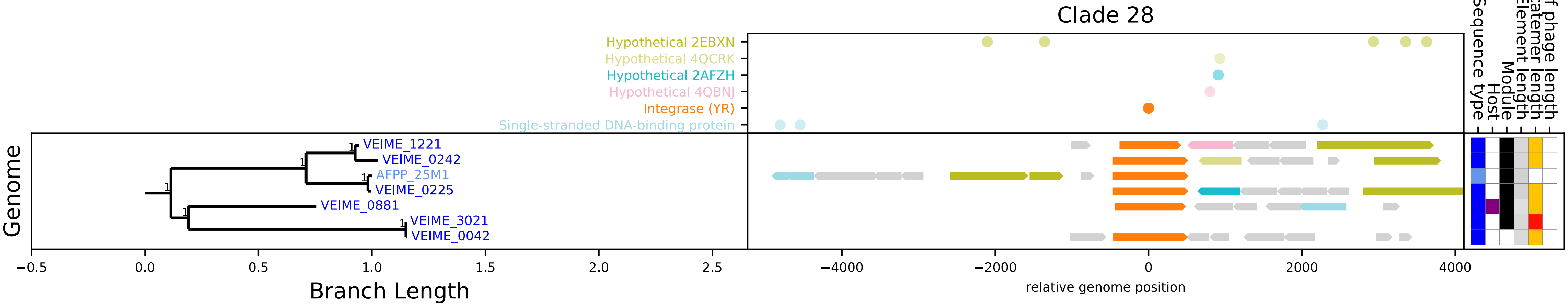

**Figure S4: VEIME Genome Maps and Annotations Organized by Tyrosine Integrase Clades, continued**

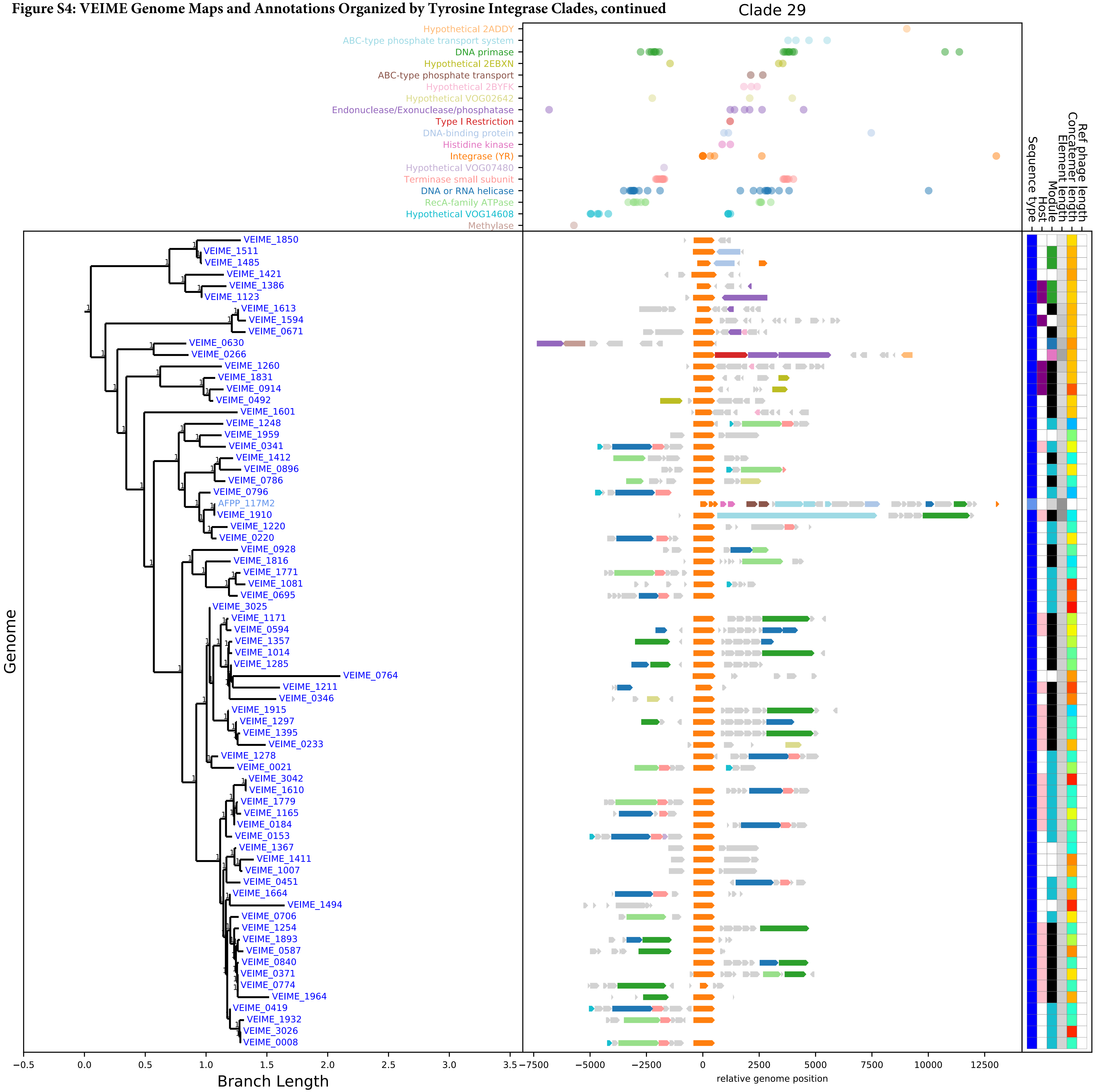

**Figure S4: VEIME Genome Maps and Annotations Organized by Tyrosine Integrase Clades, continued**

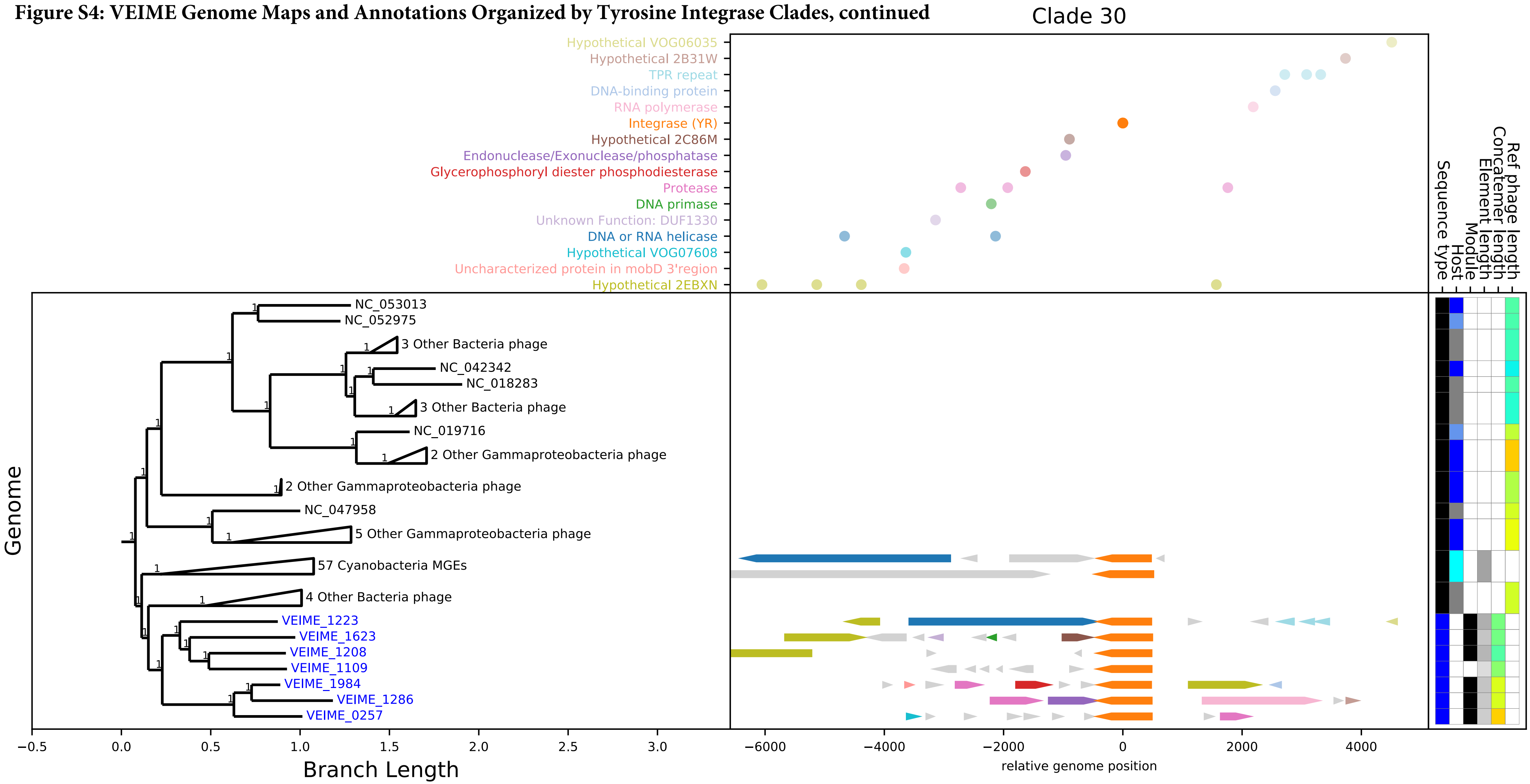

**Figure S4: VEIME Genome Maps and Annotations Organized by Tyrosine Integrase Clades, continued**

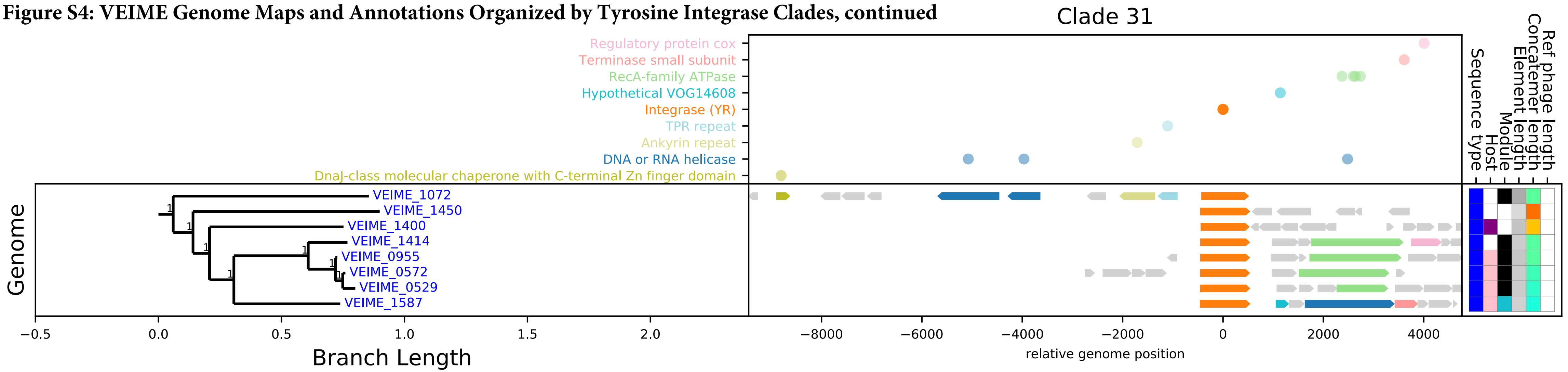

**Figure S4: VEIME Genome Maps and Annotations Organized by Tyrosine Integrase Clades, continued**

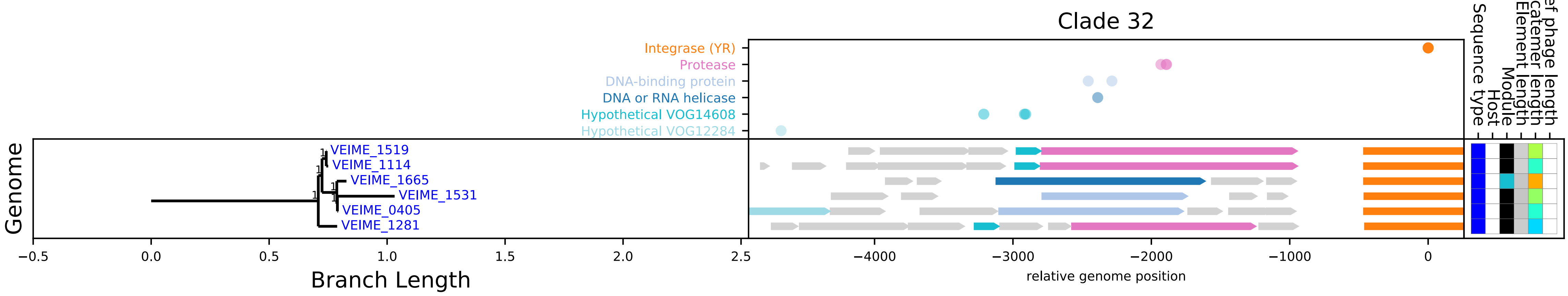

**Figure S4: VEIME Genome Maps and Annotations Organized by Tyrosine Integrase Clades, continued**

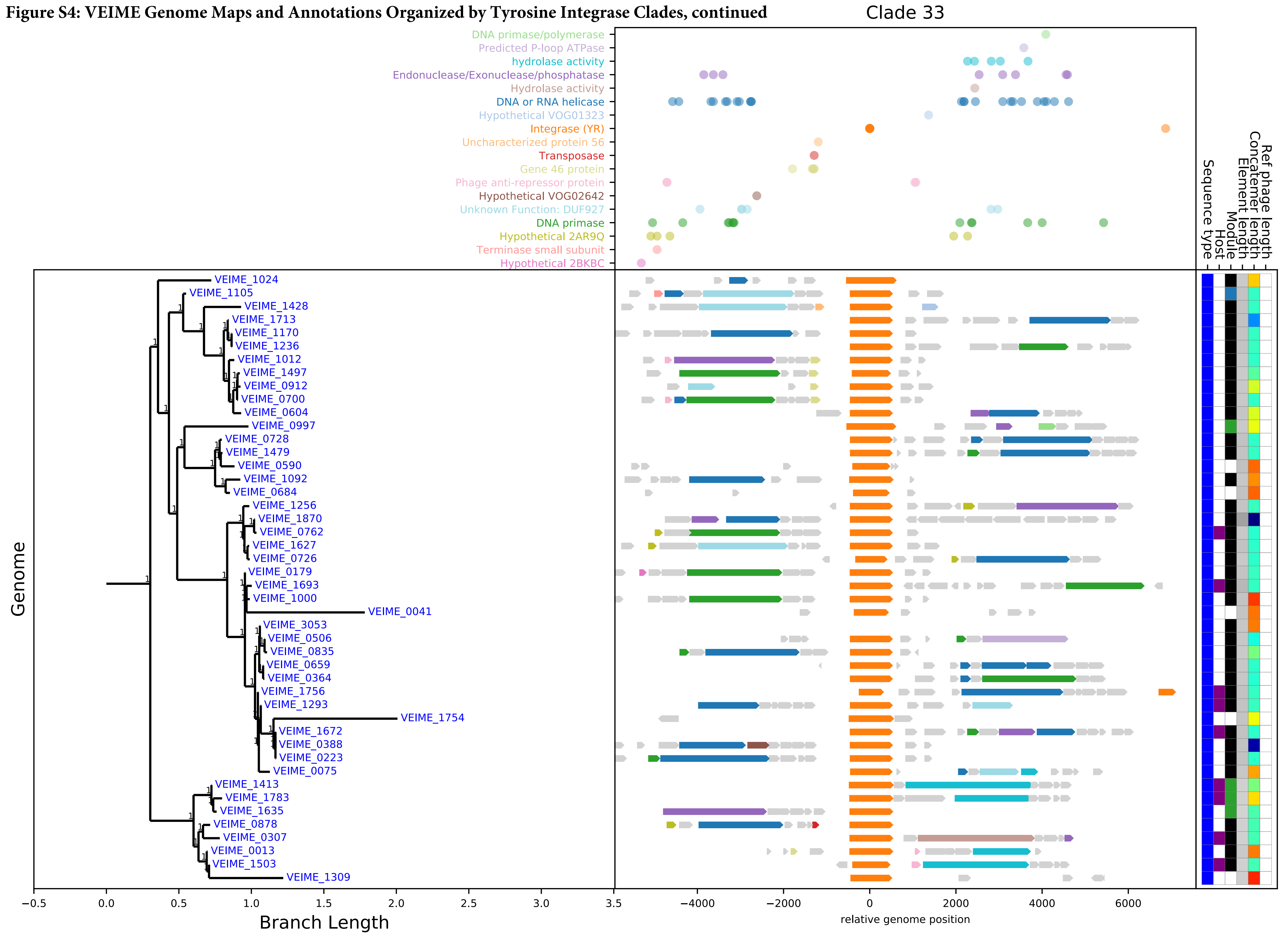

**Figure S4: VEIME Genome Maps and Annotations Organized by Tyrosine Integrase Clades, continued**

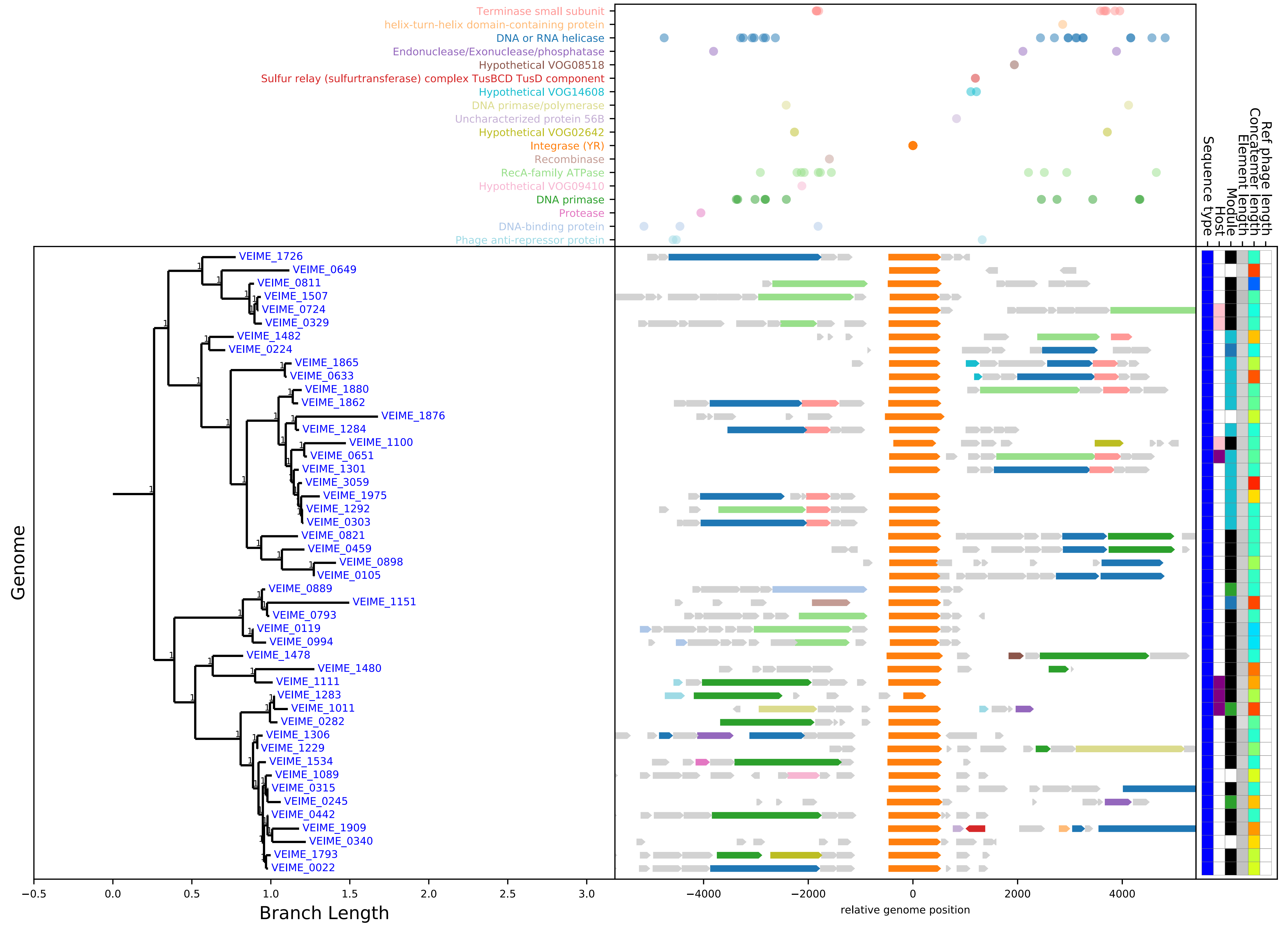

**Figure S4: VEIME Genome Maps and Annotations Organized by Tyrosine Integrase Clades, continued**

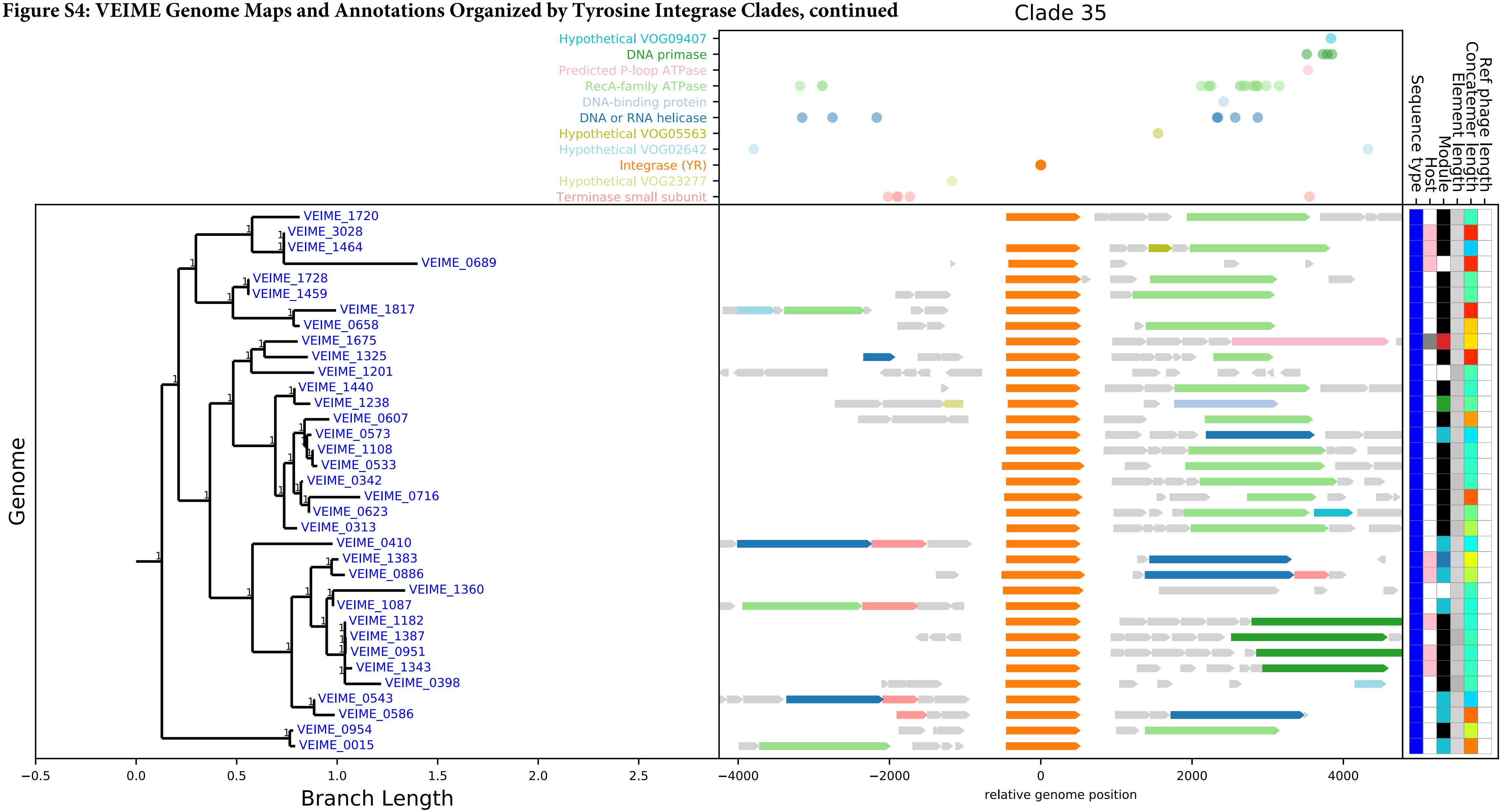

**Figure S4: VEIME Genome Maps and Annotations Organized by Tyrosine Integrase Clades, continued**

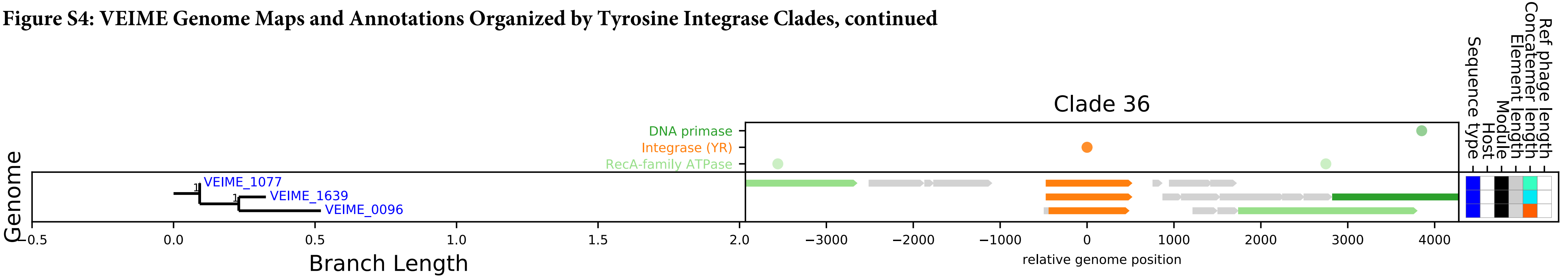

**Figure S4: VEIME Genome Maps and Annotations Organized by Tyrosine Integrase Clades, continued**

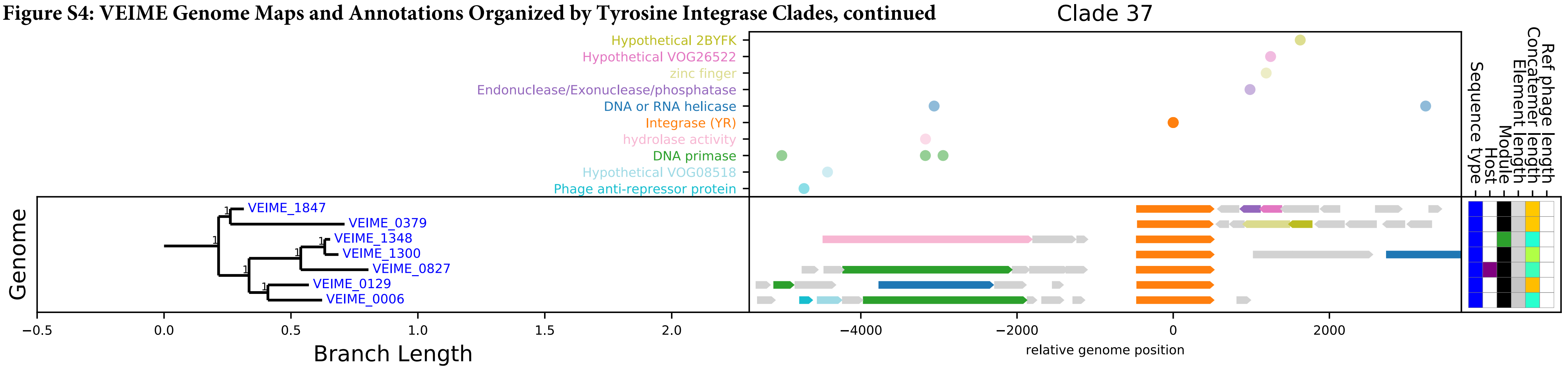

**Figure S4: VEIME Genome Maps and Annotations Organized by Tyrosine Integrase Clades, continued**

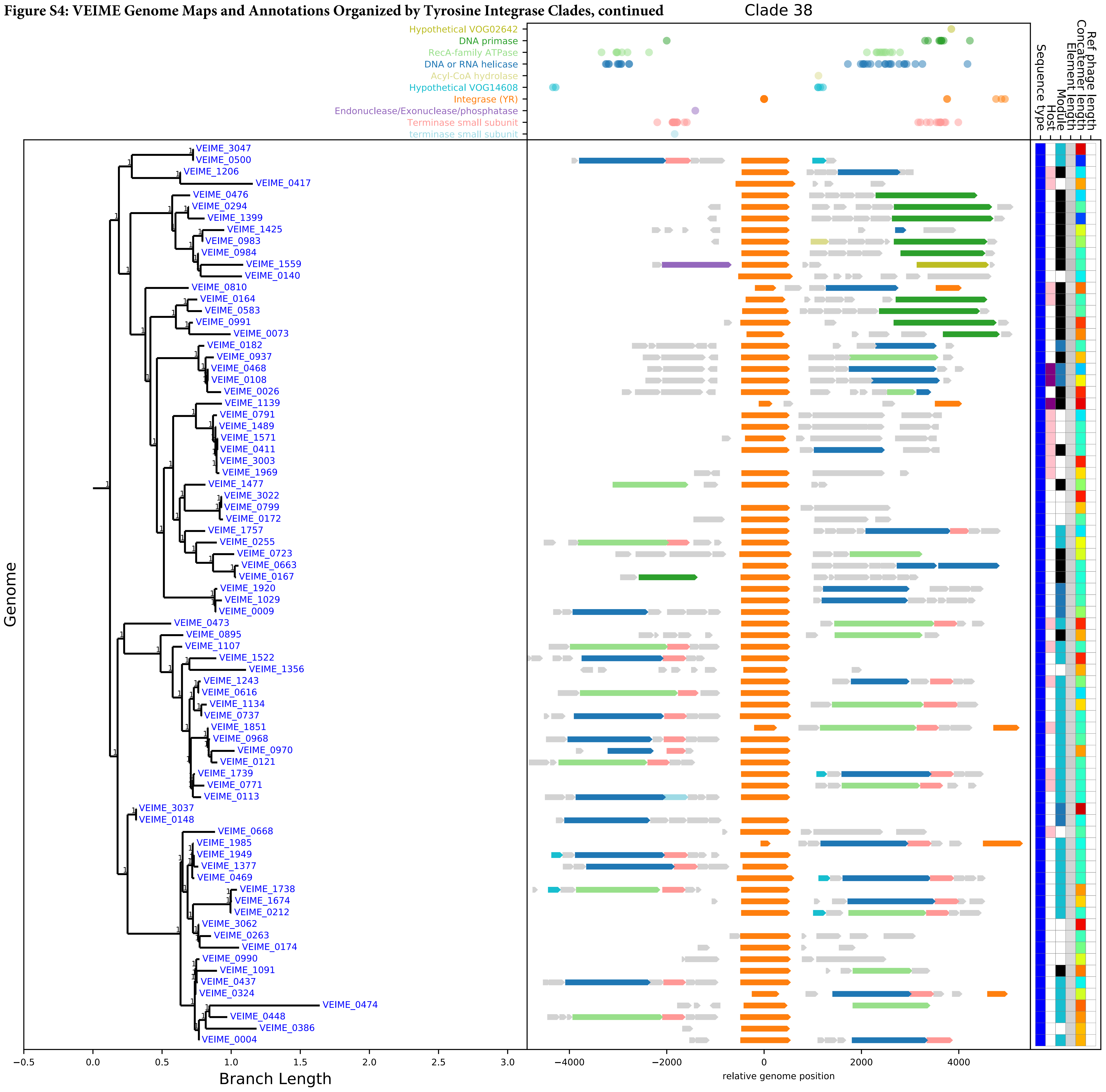

**Figure S4: VEIME Genome Maps and Annotations Organized by Tyrosine Integrase Clades, continued**

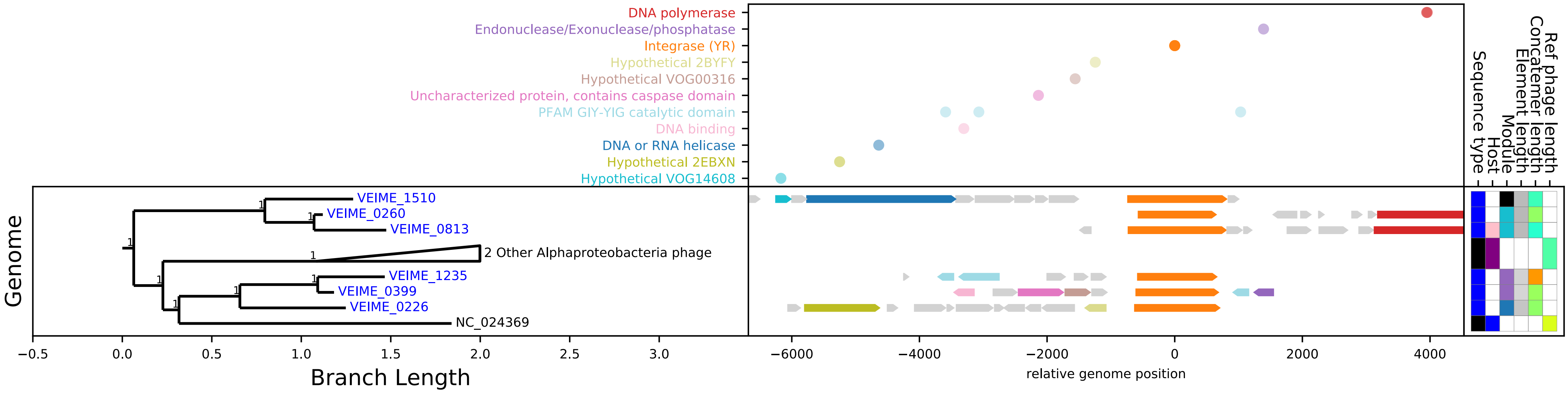

**Figure S4: VEIME Genome Maps and Annotations Organized by Tyrosine Integrase Clades, continued**

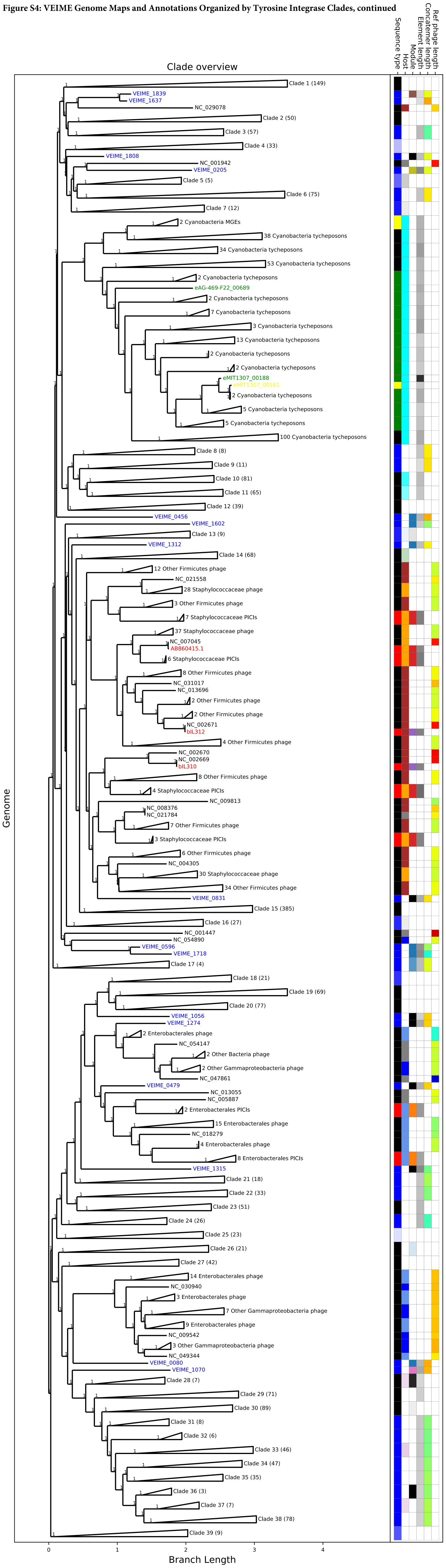

**Fig. S5 Tyrosine integrase phylogenetic tree**

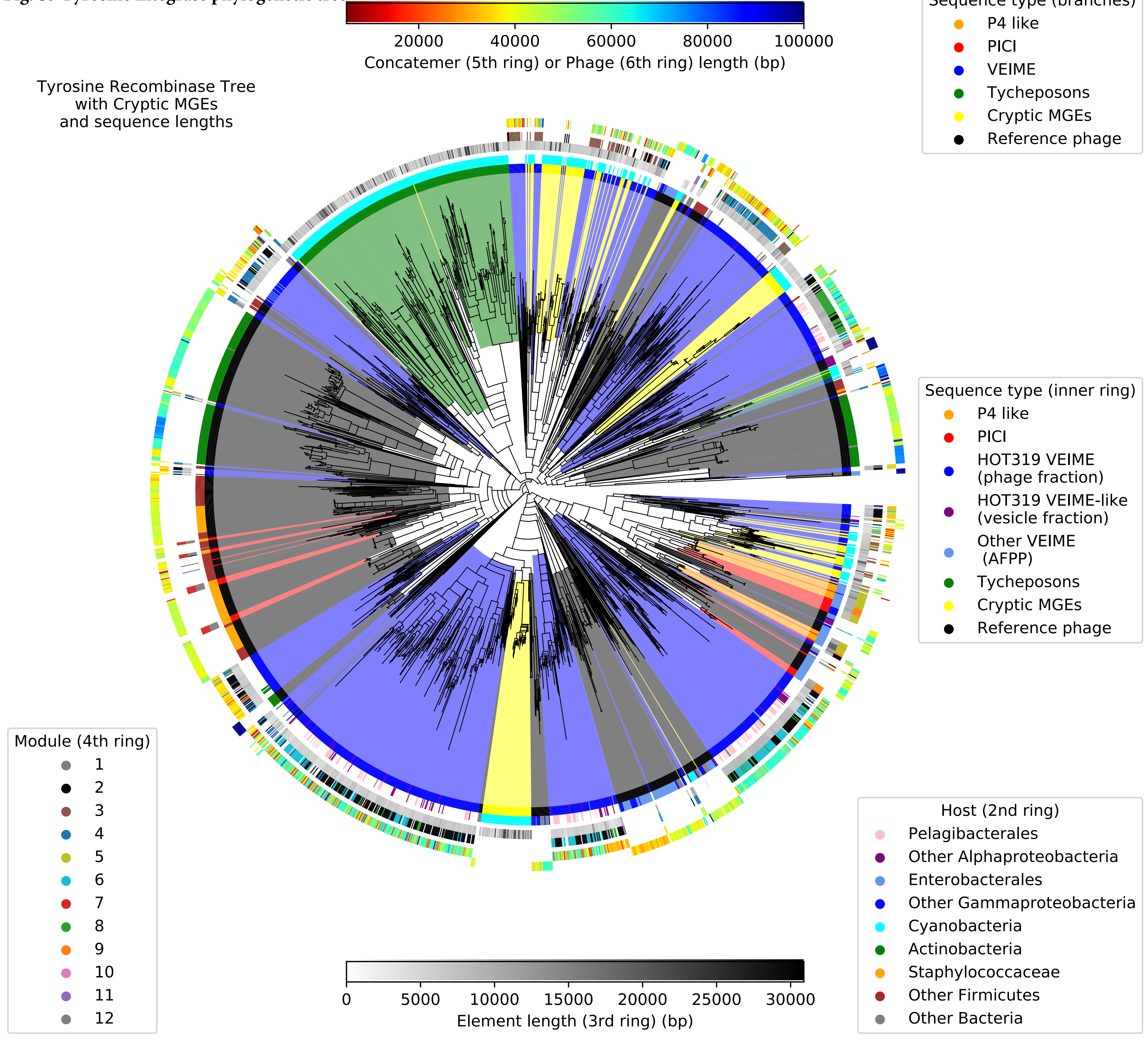

**Figure S6. VEIME Genome Maps and Annotations Organized by Serine Recombinase Clades**

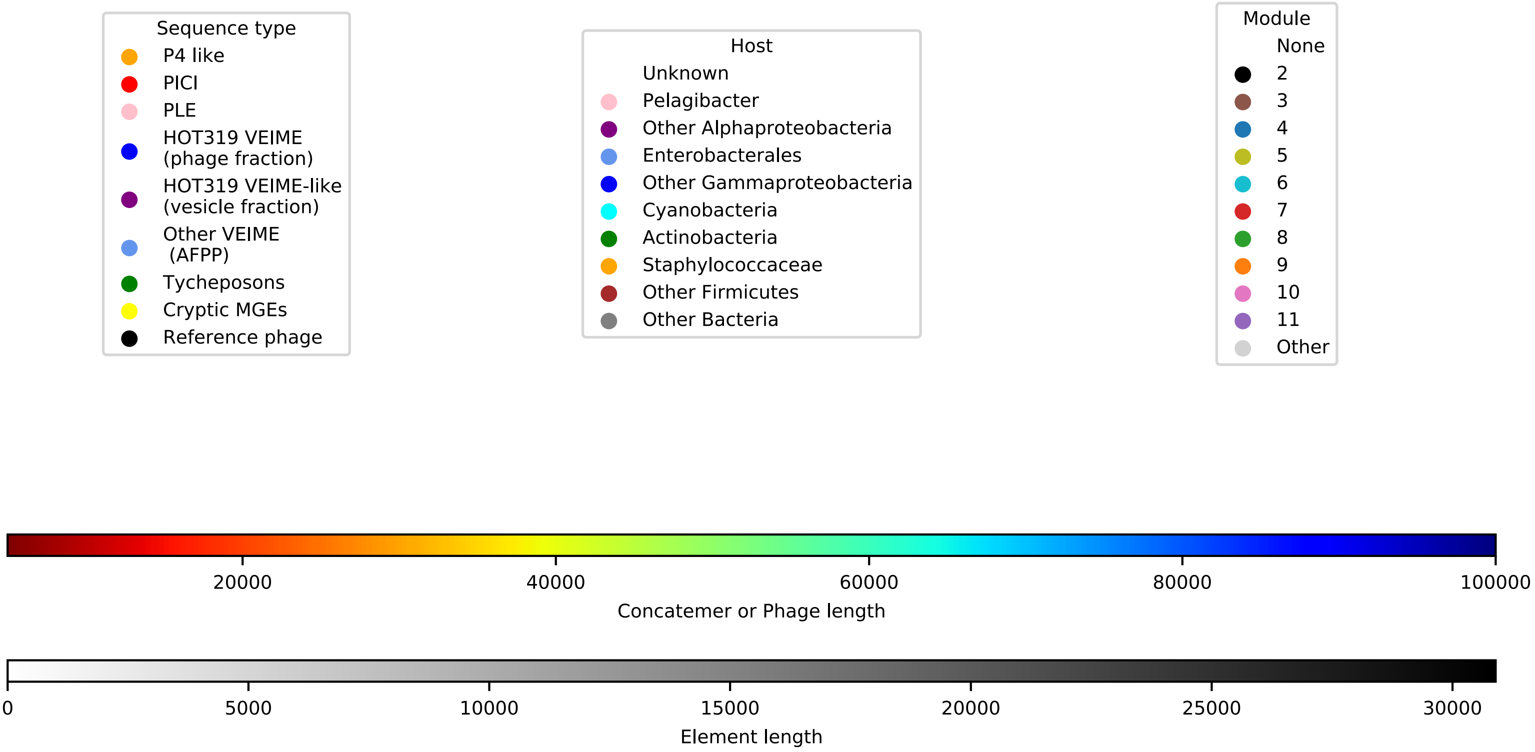


Figure S6: Serine Recombinase Tree Genome Maps
Gene annotation alignments (as in Figure S2 and S4) are draw for a Serine Recombinase maximum likelihood tree.

**Figure S6. VEIME Genome Maps and Annotations Organized by Serine Recombinase Clades, continued**

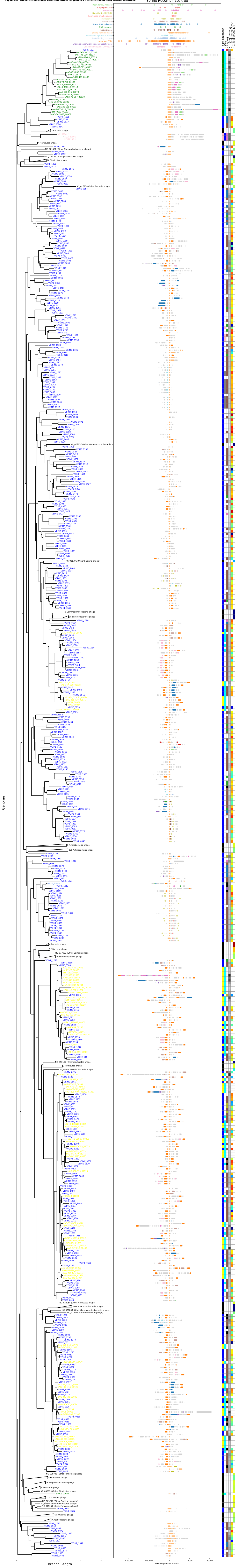

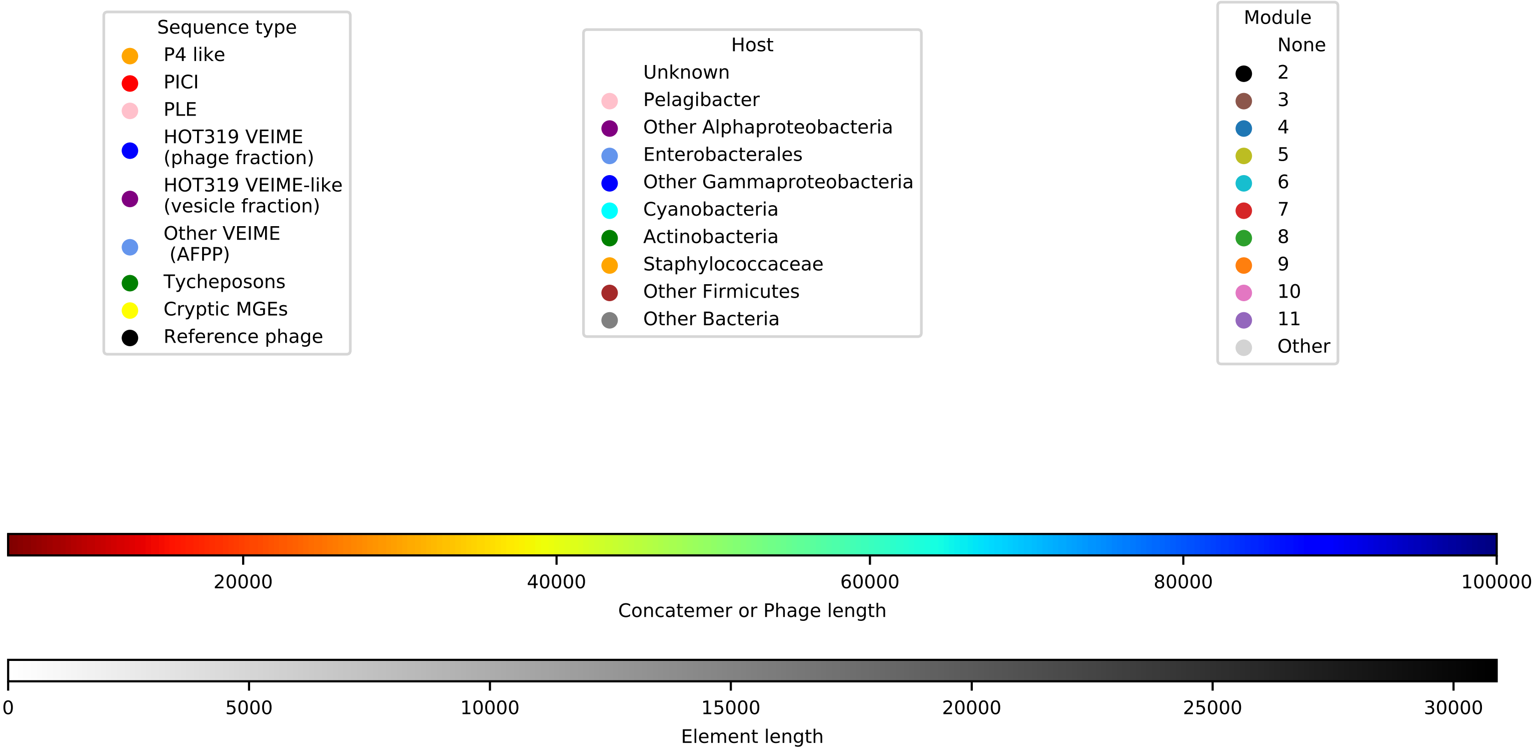


Figure S7: TerS Tree Genome Maps
Gene annotation alignments (as in Figure S4) are drawn for a tree built using the VOG06274 HMM (Terminase Small Subunit). TerS genes from bacteriophages are included for context.

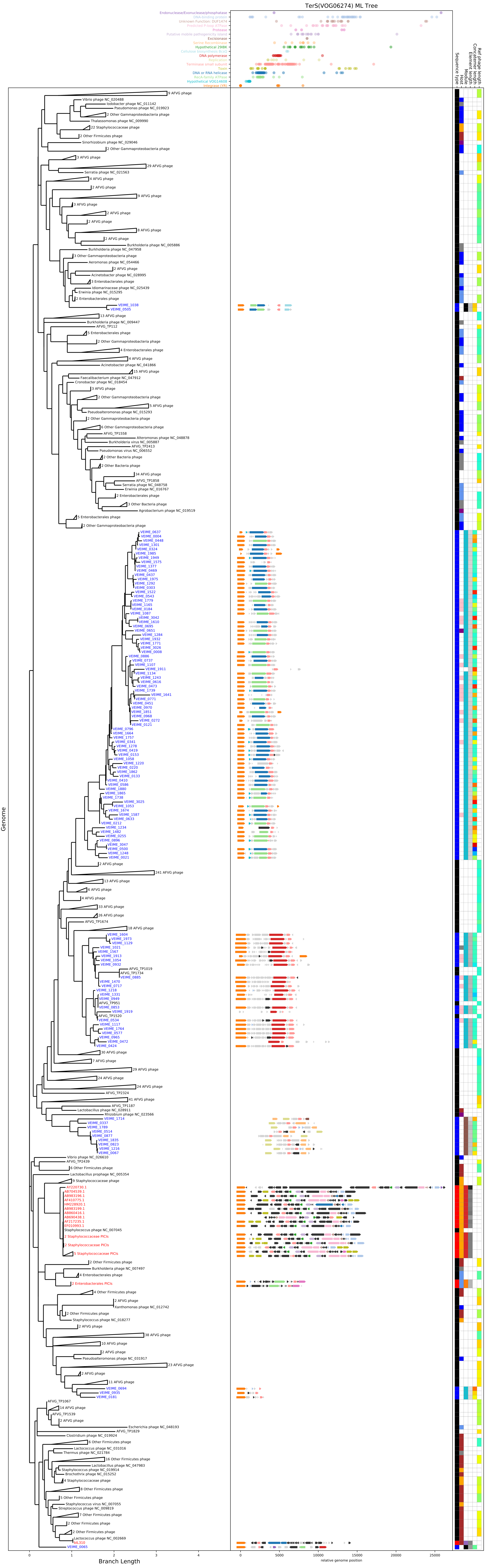

TerS(VOG06274) ML Tree
Endonuclease/Exonuclease/phosphatase
DNA-binding protein
Unknown Function: DUF1474
Predicted P-loop ATPase
Protease
Putative mobile pathogenicity island
Excisionase
Excisionase
Spore Recombinase
Hypothetical 29IBK
Cellulose biosynthesis BcsD
DNA polymerase
Replicase
Terminase small subunit
Toxin
DNA or RNA helicase
RecA-family ATPase
Hypothetical VDG14608
Integrase (YB)
Sequence type
Host
Module
Element length
Consensus length
Ref phage length
Genome
Branch Length
relative genome position
0
1
2
3
4
5000
10000
15000
20000
25000

Figure S8: Geographic distribution of VEIME abundance in TARA GOV2 samples.
Short read metagenomic covereage of VEIME alignments to GOV 2.0 contigs is avereaged for longitude bins (A) and latitude bins (B). Total coverage of all VEIMEs in each sample is averaged for combined latitude and longitude bins in (C).

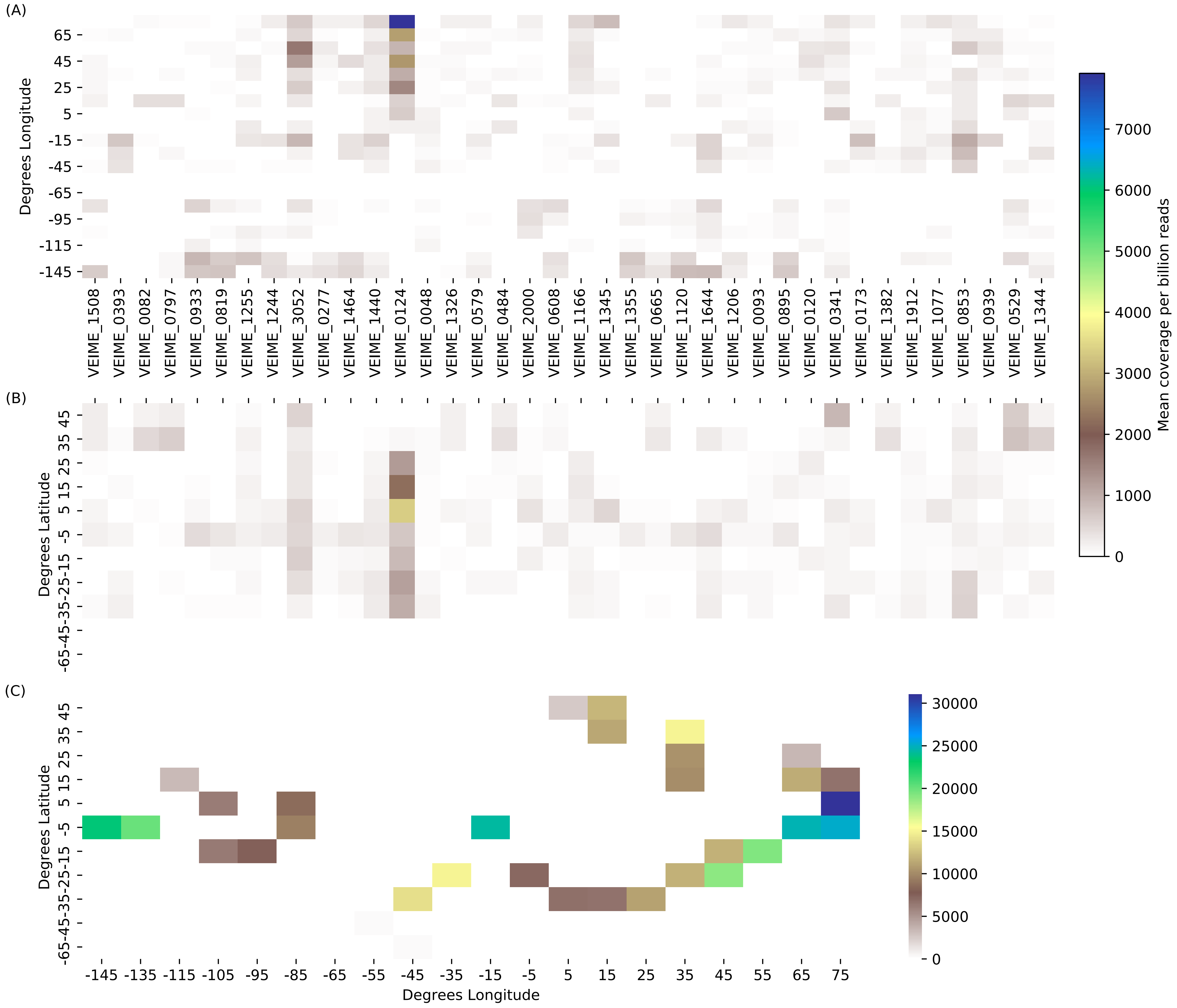

**Legends for Datasets S1 to S7**

| | |
|---|---|
| Data S1 | Bioinformatic software and databases used in these analyses. |
| Data S2 | Repeat size and number of copies for concatemeric Nanopore reads detected in the Tailed Phage and Membrane Vesicle fractions of 25m seawater collected at Station ALOHA on HOT cruise 319. |
| Data S3 | Metadata for polished VEIMEs, including those detected in the membrane vesicle fraction and those previously published as AFPPs. |
| Data S4 | Accessions, names, and host taxa of reference satellites and bacteriophage used in this analysis |
| Data S5 | Counts by module of sequence types, host taxa, and gene families, collected for all satellites (first 14 columns) and for VEIMEs only (second set of 13 columns). Gene descriptins and module assignments are in the last two columns. |
| Data S6 | Counts of gene functional annotations in VEIMEs by module, for functions present in at least 1% of VEIMEs, in at least one module |
| Data S7 | NCBI BioSample accessions for metagenomes used in this analysis. Samples from Station ALOHA were assembled as described in Materials and Methods. Assemblies for TARA samples were retrieved from iVirus. |
